# A primer on high-dimensional data analysis workflows for studying visual cortex development and plasticity

**DOI:** 10.1101/554378

**Authors:** Justin L. Balsor, David G. Jones, Kathryn M. Murphy

## Abstract

New techniques for quantifying large numbers of proteins or genes are transforming the study of plasticity mechanisms in visual cortex (V1) into the era of big data. With those changes comes the challenge of applying new analytical methods designed for high-dimensional data. Studies of V1, however, can take advantage of the known functions that many proteins have in regulating experience-dependent plasticity to facilitate linking big data analyses with neurobiological functions. Here we discuss two workflows and provide example R code for analyzing high-dimensional changes in a group of proteins (or genes) using two data sets. The first data set includes 7 neural proteins, 9 visual conditions, and 3 regions in V1 from an animal model for amblyopia. The second data set includes 23 neural proteins and 31 ages (20d-80yrs) from human post-mortem samples of V1. Each data set presents different challenges and we describe using PCA, tSNE, and various clustering algorithms including sparse high-dimensional clustering. Also, we describe a new approach for identifying high-dimensional features and using them to construct a *plasticity phenotype* that identifies neurobiological differences among clusters. We include an R package “v1hdexplorer” that aggregates the various coding packages and custom visualization scripts written in R Studio.

## 1. Introduction

More than 30 years ago, Artola & Singer (1) introduced the field of visual neuroscience to the central roles that NMDA and GABA_A_ receptors play in long-term potentiation in the visual cortex (V1) and Tsumoto et al. (2) showed the enhanced contribution of NMDARs during the critical period (CP). Thousands of experiments followed those studies, targeting specific pre- and post-synaptic proteins and providing an in-depth understanding of how neural proteins enhance or reduce experience-dependent development and plasticity in V1. More recently, proteomic and genomic studies are surveying thousands of proteins and genes to explore novel mechanisms regulating development and plasticity in V1 (3,4).

The shift from studying a few proteins to quantifying tens to thousands of proteins is changing our understanding of visual cortical development and plasticity but it also poses new challenges for data analysis. Here we describe two workflows for using high-dimensional analyses to study the development and plasticity of neural proteins (or genes) in V1. We take advantage of insights gained from previous studies about the role of different proteins in experience-dependent development and plasticity to select a targeted set of proteins to study. Furthermore, by working at the level of proteins, we can apply the same techniques to studying V1 in animal models (section 3) and humans (section 4).

Our aim is to describe workflows and provide examples for high-dimensional analysis of protein (or gene) data using the statistical software R. The examples address how to use the workflow to discover experience-dependent or lifespan changes in plasticity mechanisms. Also, we describe a novel approach to exploring and comparing the neurobiological features that characterize different rearing conditions or age groups that we call the *plasticity phenotype*. The goal of building the plasticity phenotype is to help uncover meaningful insights into V1 plasticity from the high-dimensional patterns of protein expression.

### 1.i) Getting started and contributions

The first challenge in developing the workflows was to determine which high-dimensional analyses were appropriate for our experimental designs. Our experiments include many proteins with known roles in neuroplasticity, and often the tissue samples come from multiple cortical regions. That experimental design has more variables (*p*) than conditions (*n*), so the data sets are *p>n* and are by definition high-dimensional. The data sets are also described as *sparse* because the distance between samples in the high-dimensional space is uneven. This sparse structure means that special consideration is needed in selecting methods for data analysis. First, the methods must support the discovery of clusters that differ on only a few proteins or combinations of proteins (features). Second, those discoveries need to guide meaningful insights into how V1 develops and changes with different forms of visual experience. To this end, we developed workflows that lead to the construction of plasticity phenotypes using features identified by high-dimensional analyses. Also, we implemented a visualization for the plasticity phenotype that facilitates the intuitive exploration of the data.

Here we describe and compare two approaches suitable for analyzing a targeted set of plasticity proteins (*p*) and comparing among rearing conditions (*n*) or developmental stages (*n*).

The intent of this paper is not to review high-dimensional analyses or to determine the “best” analysis, but rather to demonstrate and discuss two workflows appropriate to cluster and classify plasticity phenotypes in the developing visual cortex.

### 1.ii) Contributions of this paper

1. We provide two workflows with examples for identifying clusters in a sparse data set (*p≅n or p>n*). With each workflow we discuss considerations for selection of the analysis steps.
2. We show that it is necessary to explore different techniques to select the clustering method that is appropriate for the data set and research questions.
3. We combine tools for partitioning data points into clusters and identifying plasticity features. The features are used to create the plasticity phenotype that aids in meaningful interpretation of the data.

The rest of this paper is organized as follows. In section 1.iii, we review some of the high-dimensional data analysis methods that have been used in recent papers studying cortical development. In sections 3 and 4, we present the workflows with examples using PCA and tSNE, or sparse high-dimensional clustering, and describe how to build and use the plasticity phenotypes. Section 5 provides a brief summary and discussion.

### 1.iii) Past work using high-dimensional analysis

#### Principal Component Analysis

The most commonly used high-dimensional analysis for exploring gene or protein expression in the brain has been principal component analysis (PCA) (5,6). PCA transforms the gene or protein data, which is likely to include correlated genes or proteins, into a linear set of uncorrelated principal components that capture successively less of the variance in the data. Thus, individual cases can be visualized and analyzed within the transformed lower-dimensional space and that is often helpful for identifying clusters in the data (Figure 1). For example, a recent survey of human brain development using RNA and protein expression (6,529 proteins were found in all 7 brain regions) used PCA to reduce the dimensionality of the data and identify differences among brain regions (3). That analysis highlighted the separation of cerebellar samples from the other brain regions, but it is challenging to interpret the biological significance of the PC that differentiate the regions.

**Figure 1.**
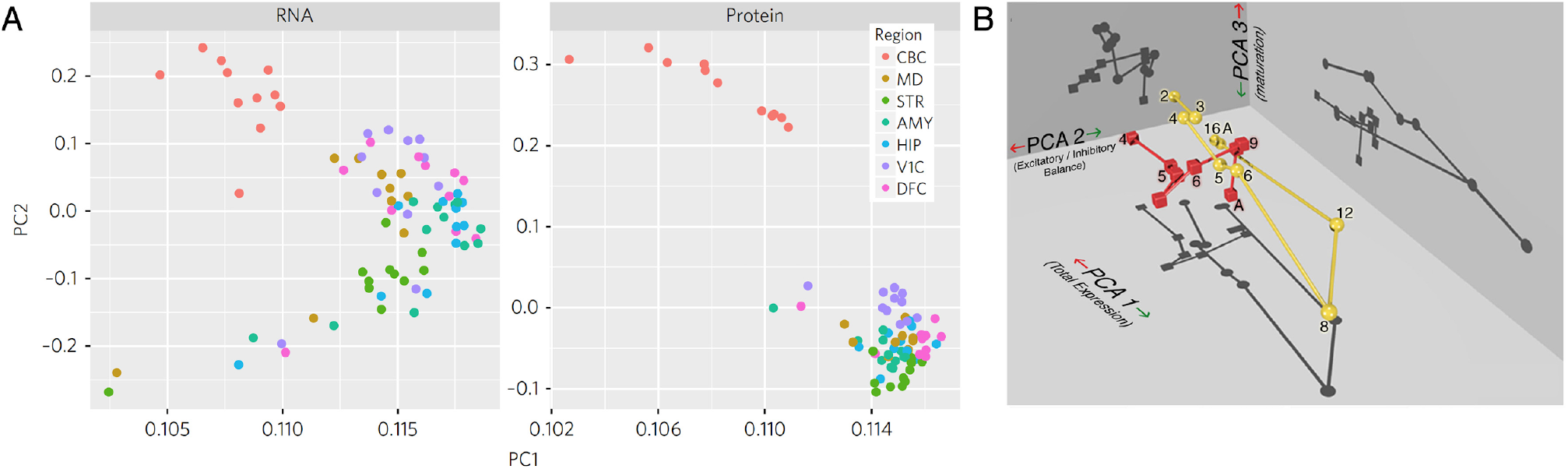
Interpreting PCA dimensions with either thousands or tens of genes or proteins: (**A**) Scatter plots for the first two dimensions from a PCA analysis of RNA (5141 genes) or protein (6529 proteins) expression for postmortem tissue samples from 7 regions of the human brain. The plots show a clear separation of cerebellum samples (CBC) from the other brain regions. It is challenging, however, to interpret the biological correlates that differentiate among the regions because the very large number of genes or proteins were reduced to just 2 unitless components. *Reprinted with permission from Becky C. Carlyle et al: Springer Nature, Nature Neuroscience, A multiregional proteomic survey of the postnatal human brain, Carlyle et al., 2017* (**B**) A plot of the first 3 dimensions from a PCA analysis of 7 proteins and 3 regions for normally developing (yellow spheres) or monocularly deprived animals (red spheres). The shadows projected on the three walls help to visualize differences between normal (circles) and deprived (squares) animals. Age (in weeks) is displayed beside each symbol and the connecting lines link the points by age. The biological correlates for each dimension were determined using the basis vectors -- PCA 1 reflects the sum of the protein expression, PCA 2 reflects an aspect of the E:I balance, and PCA 3 reflects the maturation of receptor composition. *Reprinted from Beston et al., 2010*.

The unitless dimensions of PCA components make it hard to identify the biological correlates when thousands of genes or proteins have been quantified and this often leads to the use of pseudo-units (e.g. pseudoage). In contrast, when a targeted set of genes or proteins with known functions are studied then the basis vectors for each component (the weights for each protein) can be used to attach biological significance to otherwise unitless dimensions. For example, after applying PCA to the expression of 7 synaptic proteins from animals reared with normal vision or monocular deprivation (MD), we used the basis vectors to infer that PC1 reflected the sum of the proteins, PC2 an aspect of the excitatory:inhibitory balance, and PC3 the maturational state of the subunit expression ((7); Figure 1B).

In section 3, we describe a two-step process for using PCA; first, the typical step for dimension reduction, and second, a new step that expands each dimension using the basis vectors to identify the biologically relevant features that account for variance in the data. Those features become the building blocks for the *plasticity phenotype* and facilitate interpretation by linking the features with known functions for regulating experience-dependent plasticity. Also, the overall pattern of features can be used to provide robust phenotypic information about the biological correlates that identify clusters in the data. Thus, the plasticity phenotypes help to discover meaningful insights into how V1 matures during normal development or is changed by abnormal visual experience.

#### t-Distributed Stochastic Neighbor Embedding

Another popular method for transforming and visualizing high-dimensional data is t-Distributed Stochastic Neighbor Embedding (t-SNE, (8)). tSNE measures the shortest distance between pairs of data points, then calculates pairwise probability estimates of similarity across *all* dimensions. Next, these estimates are mapped onto 2-dimensional (2D) space by scaling the distance between data points and positioning similar data points closer together. The new mapping preserves local and global patterns thereby representing the relationships among data points that may highlight clusters in the data. The artificial scaling makes it easier to identify clusters by either color-coding points based on a known attribute (e.g. cortical area), or by applying a clustering method to the tSNE XY coordinates. Furthermore, the unsupervised nature of tSNE is particularly useful when exploring data without strong *a priori* knowledge of the expression patterns that may differ among the conditions.

For example, a combination of PCA and tSNE was used to analyze the data from a recent study of single-cell mRNA expression in the developing human brain ((4), Figure 2A). In this example, PCA was used to reduce the dimensionality of the data and then tSNE to identify and visualize clusters (colour-coded dots) of samples (Figure 2B-D). Next, the tSNE plots were used to show clustering by lineage (e.g. CP or GZ)(Figure 2C) and cell type (e.g. DLX1 for MGE derived cells) (Figure 2D).

**Figure 2.**
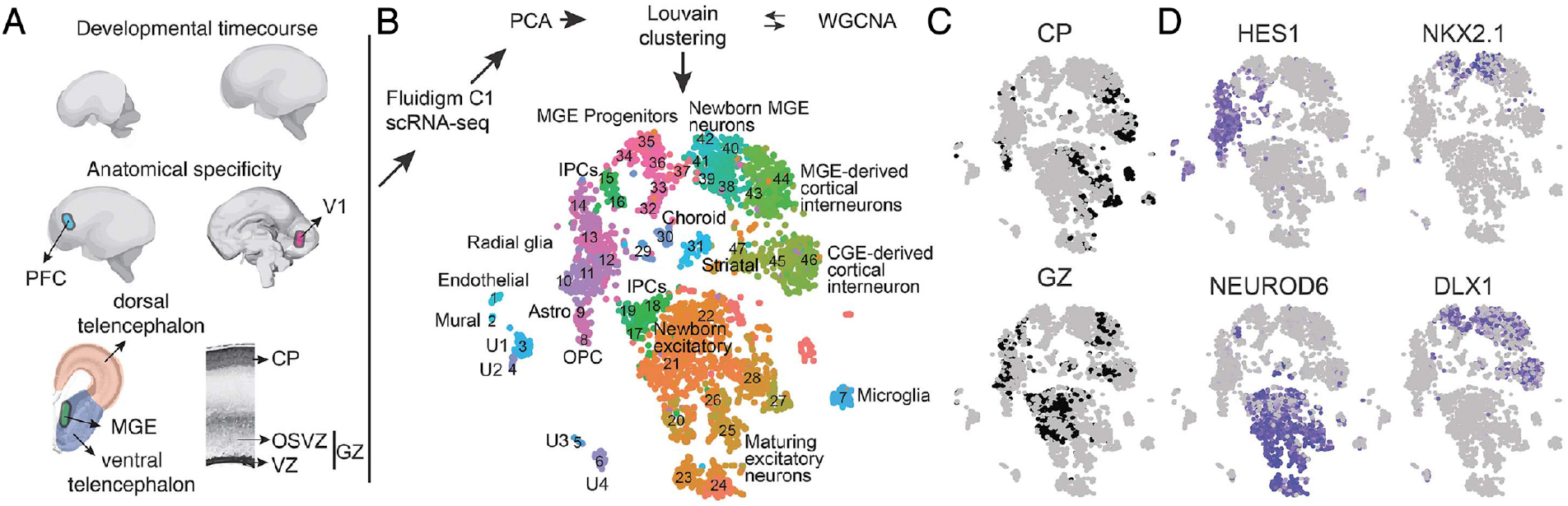
Applying tSNE to human brain cells identifies cell type clusters. Diverse cell types in human telencephalon development. (A) Schematic illustrating sample collection over time, region, and lamina. (B to D) Scatterplot of 4261 cells after principal components analysis and t-stochastic neighbor embedding (tSNE), coloured by (B) cluster, (C) cortical lamina source, and (D) maker gene expression. *From Nowakowski, T. J., Bhaduri, A., Pollen, A. A., Alvarado, B., Mostajo-Radji, M. A., Di Lullo, E., … & Ounadjela, J. R. (2017). Spatiotemporal gene expression trajectories reveal developmental hierarchies of the human cortex. Science, 358(6368), 1318-1323. Reprinted with permission from AAAS.*

This is a powerful workflow for analyzing and visualizing complex gene or protein data. It is a good illustration of a common approach that starts by reducing the dimensions down to 1-5 components and using those eigenvalues as input to the cluster analysis. However, care is needed when using dimension reduced data as input to clustering because the orthogonal components from PCA may not contain the features need to partition the data into clusters (9).

In section 3, we describe a modified workflow that uses tSNE to visualize clusters and PCA to select features for constructing plasticity phenotypes but does not use the transformed eigenproteins as the input to the tSNE analysis. We demonstrate this workflow using a data set comparing protein expression in V1 among a set of rearing conditions where animals had either normal visual experience, monocular deprivation (MD) or MD plus a subsequent treatment. We present the steps for this new workflow and explain how it can be used to explore and interpret cluster composition based on rebranding each sample by their plasticity phenotype.

#### Other clustering approaches

In addition to PCA and tSNE, there are many other algorithms that can be used to discover the natural clustering of samples based on similar patterns of features. Two major classes of clustering algorithms are hierarchical and partitional techniques. Hierarchical clusters are built through a top-down (divisive) or bottom-up (agglomerative) approach: top-down begins with all the data in a single cluster that is recursively divided into smaller clusters, or bottom-up begins with each data point in individual clusters that are recursively merged. Hierarchical clustering continues until a designated threshold is met, usually the number of clusters (*k*). One of the strengths of that approach is the use of a dendrogram to graphically represent clusters by ordering samples with similar features nearby in a tree structure. For example, the matrix in Figure 3B used hierarchical clustering to order the strength of pairwise correlations between brain areas for a set of 123 proteins that are differentially expressed across human development (Figure 3B; (3)). The strength of the correlations was colour-coded, and the hierarchical ordering made it easy to see the clustering of cerebellar samples based on their poor correlations (blue) with the other areas. It can be difficult, however, to see more subtle differences in the pattern of correlations among the other areas and to select a height in the tree to define more subtle clustering.

**Figure 3.**
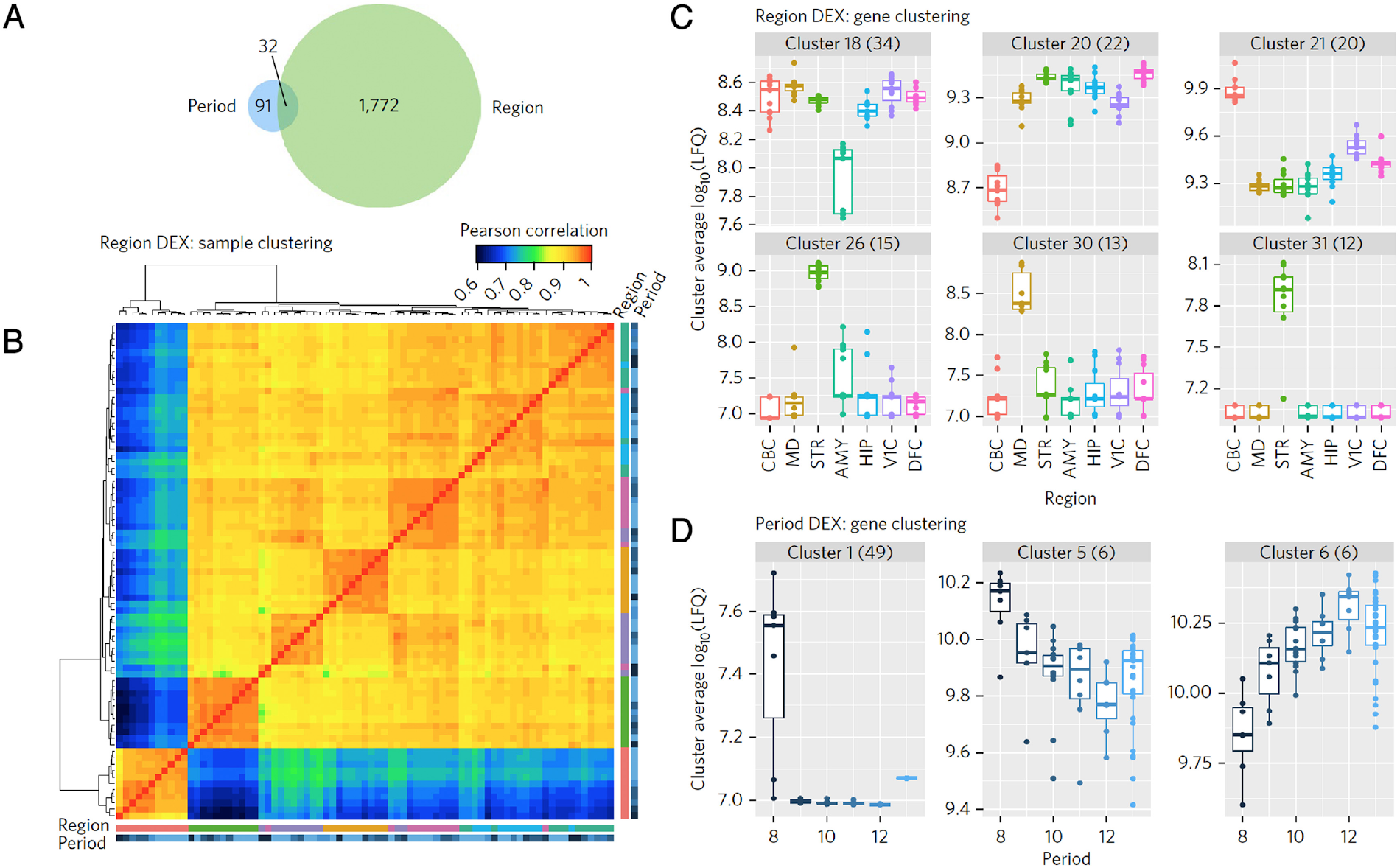
Hierarchical clustering of samples in a correlation matrix. **a**, Venn diagram showing the number of genes that were significantly and differentially expressed across developmental periods (left, blue) or brain regions (right, green). **b,** Hierarchical clustering revealed major divisions between brain regions, but less distinct classification based on developmental period. **c-d**, Exploring clusters by region (**c**) reveals region-specific enrichment or depletion and exploring across developmental periods (**d**) reveals developmentally enriched period of protein expression shortly after birth (period 8). Each data point is shown as a dot. *Reprinted with permission from Becky C. Carlyle et al: Springer Nature, Nature Neuroscience, A multiregional proteomic survey of the postnatal human brain, Carlyle et al., 2017*

The other major class of clustering algorithms, partitional, does not impose hierarchical structure to the data and instead finds all of the clusters at the same time as a partition of the data. *K*-means is the best known and most widely used partitional clustering method. It starts by partitioning samples into *k* clusters where the number of clusters is chosen using the gap statistic to estimate the number of clusters in the data set (10). The data are then iteratively repartitioned into different clusters to minimize the within-cluster sum of squares. Since this method is not hierarchical, the clusters cannot be represented in a dendrogram, so *K*-means or other partitional clustering methods (e.g. Louvain, Infomap, etc) are often visualized using tSNE. For example, in the workflow diagram from Nowakowski et al (2018) the color-coded clusters in the tSNE plot were identified using Louvain partitional clustering.

Many clustering approaches work well when there is good separation among the features in the protein or gene data set. In contrast, when samples differ on a small fraction of the features or there are more subtle changes in protein expression the standard clustering methods begin to fail and sparse clustering algorithms are needed. The advantage of sparse clustering is that it uses an adaptive selection of a subset of features within hierarchical or *K*-means clustering where the selection of features is iteratively optimized using a regression-style analysis (e.g. lasso) (11).

The data set used for section 3 had good separation of the features because the different rearing conditions drove large changes in protein expression. In the second workflow (section 4), however, we used an example data set with more subtle variations in protein expression that required application of a modern sparse clustering algorithm to uncover age-related clustering in development of human V1.

Finally, whether clustering is done with hierarchical, partitional, or sparse algorithms the same challenge remains -- how to link the holistic exploration of the data with the biological features that differentiate the groups (e.g. ages, cortical areas, rearing conditions). In previous studies, the task of pinning down those features was often done by sorting through the clusters using a series of plots and univariate analyses aimed at finding genes/proteins that are over- or under-expressed in a group (3,12). That approach, however, focuses on measurements from just one variable per sample and thereby loses sight of differences that arise from higher order combinations of protein or gene expression. That problem prompted us to develop a method for discovering combinations of proteins that represent high-dimensional features and using those to construct the *plasticity phenotype*. While the idea of a phenotype is not new, our approach to extracting features from the protein data and using them to analyze and visualize the plasticity phenotypes is a novel application in this field of study.

## 2. Example data sets and preparation of the data

In this paper, we used two data sets of neural protein expression in V1 obtained using Western blotting and quantification with densitometry. Each data set was organized into a matrix with *n* rows of observations (e.g. cases, cortical regions, and number of Western blot runs) and *p* columns of variables (e.g. # of proteins). The first data set included results from animal studies examining changes in glutamatergic and GABAergic receptor subunit expression during normal development, monocular deprivation (MD), or treatment after MD (7). The second data set was from a series of studies examining the development of human V1 by measuring expression of a collection of neural plasticity proteins in postmortem tissue samples from cases that range in age from neonates to older adults (13-18).

Prior to beginning the high-dimensional analyses described in this paper it is important to inspect and organize the raw data set. First, if using Western blotting ensure that the quantification of the bands did not include artifacts (e.g. bubbles, spots) or poorly labeled bands that could skew the results. Those data points should be omitted, and the missing data can be filled by imputation. A variety of imputation functions have been implemented in R and a recent package *impute* was developed for microarray data that imputes missing gene or protein expression data using a nearest-neighbor analysis (19).

## 3. Using PCA & tSNE to study experience-dependent changes in visual cortex: Data reduction, feature identification, clustering, plasticity phenotypes

The first workflow describes using PCA to identify features and then clustering of the features with tSNE. A novel aspect of this approach is using the features to construct plasticity phenotypes and applying those phenotypes to rebrand clusters into meaningful groups.

The data set comes from two animal studies of visual cortical development and plasticity (7) with an *n*x*p* matrix comprised of *n*=567 rows of observations and *p*=7 columns of protein variables (Tables 1&2). The final matrix had 3,969 cells of data and after omitting 602 cells with poor labelling, the final number of data points was 3,367.

**Table 1.** Observations (*n*)

| **Categories** | **Specific** | **Total** |
| --- | --- | --- |
| Rearing Condition | Normal (9), MD (8), Reverse Occlusion (1), Binocular Deprivation (1), Binocular Vision (5) | 24 |
| Regions | Central (2), Peripheral (8 or 10), Monocular (2) | 12 or 14 |
| WB Runs | 1,2 | 2 |
|  | **Sum** | **567** |

**Table 2.**
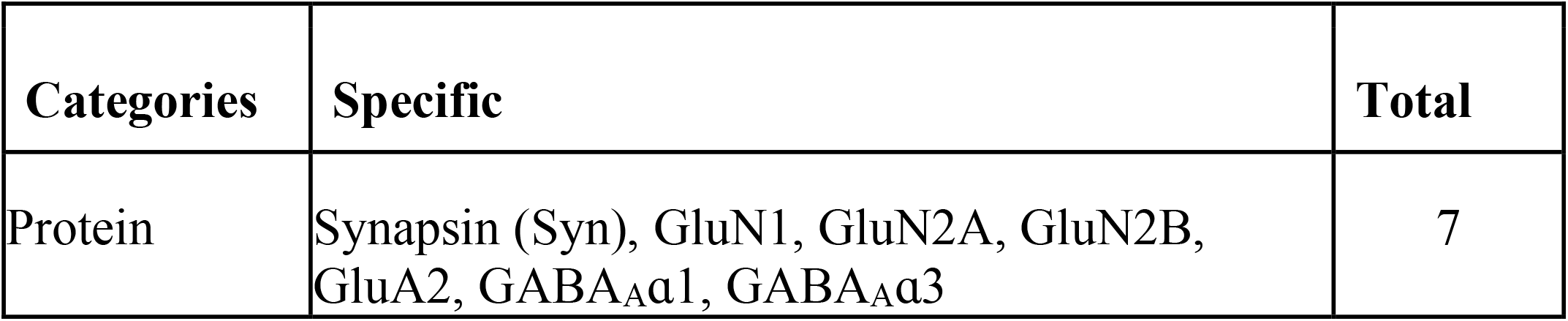
Variables (*p*)

### 3.i) Quantification and analysis

The natural first step in the analysis workflow is to find extremes in protein expression that identify significant differences between conditions. Using a series of univariate analyses, however, becomes overwhelming very quickly as the number of genes or proteins quantified increases (Figure 4B). Furthermore, such an approach does not realize the potential of the high-dimensional data set since it is not inclusive of the full repertoire of proteins available. Instead, holistic approaches that examine all proteins can identify patterns in the data that suggest how the biological functions might have changed.

**Figure 4.**
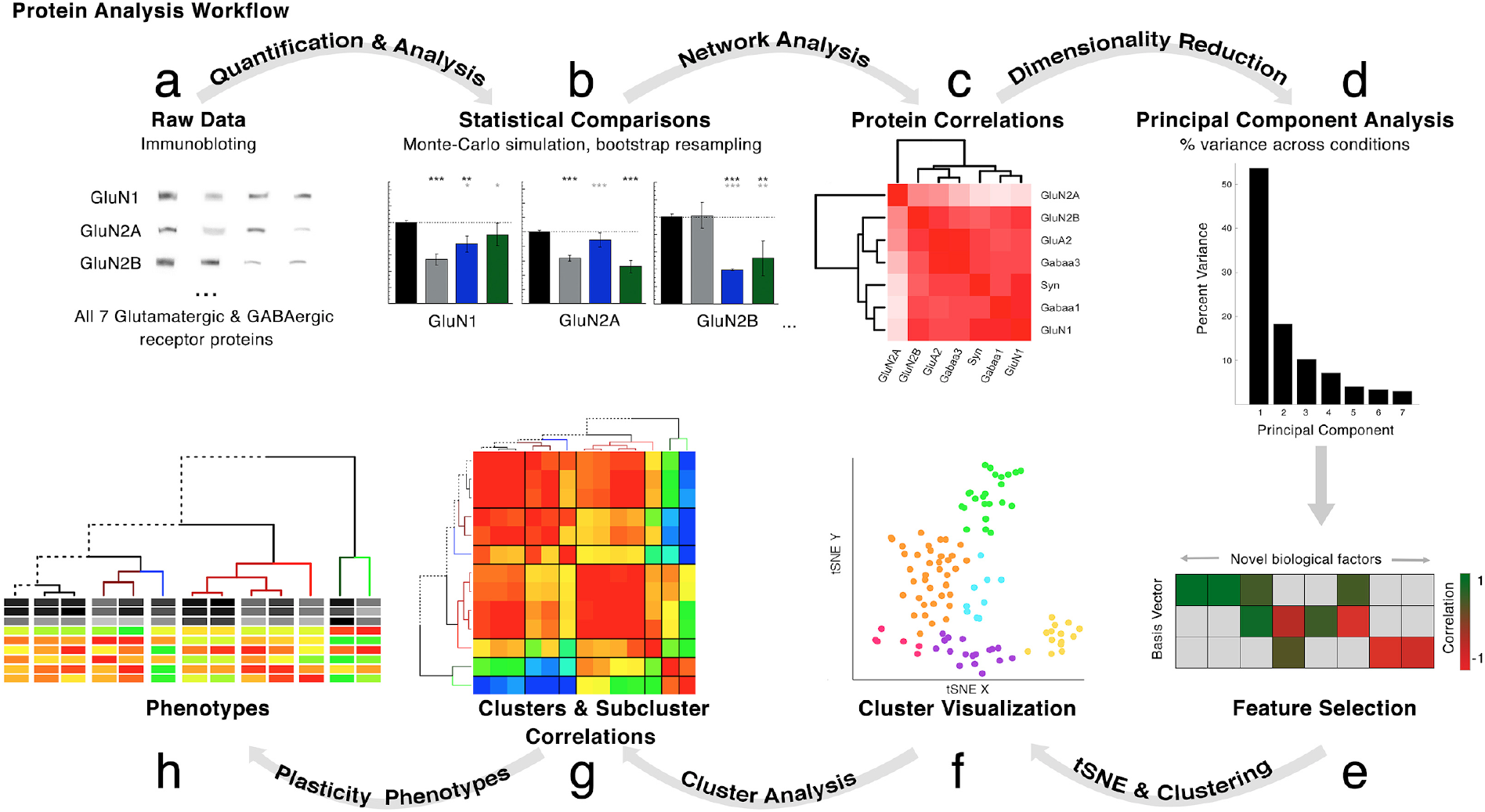
Protein analysis workflow for p<10 and p≅n. **a**. Protein expression was collected across 7 proteins using immunoblotting (N=4232). **b**.Standard univariate analyses to identify significant differences among the 9 rearing conditions. **c**. Network analyses calculated using Pearson’s R between the 7 proteins in each rearing condition. **d.** PCA to explore and **e**. transform features. **f**. tSNE to represent the data in low-dimensional space (2D), then clustering algorithms were applied to the low-dimensional representation of the data. Clusters were annotated and subclusters identified (coloured dots) using known information about the tissue samples (rearing conditions). **g**. Network effects assessed using Pearson’s R correlations between clusters & subclusters **h**. Plasticity phenotypes to visualize similarities/differences among subclusters.

Next, we describe using pairwise correlations and hierarchical clustering to visualize patterns in the data using a 2D correlation heatmap. The organization of positive and negative correlations provides insight into the network of protein expression in visual cortex and how different visual rearing conditions changed it.

### 3.ii) Network analysis

Visualizing pairwise correlations between proteins was the first step to beginning a more holistic analysis of the data. The order of the proteins was sorted by hierarchical clustering so that proteins with similar patterns of correlations were nearby. A dendrogram was used to visualize the tree of protein clusters in the data (Figure 4C).

The analysis was done in R Studio using a series of packages available for download at the Comprehensive R Archive Network (CRAN). These packages include: *Harell Miscellaneous, stats, dendextend, seriation,* and *gplots*.

First, the data file was read in to an object called `my.data`.

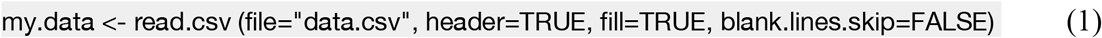

Then the data were parsed into subsets representing each of the rearing conditions. The following R coding example (2) demonstrates how to parse the data for the Normal rearing condition (Normal):

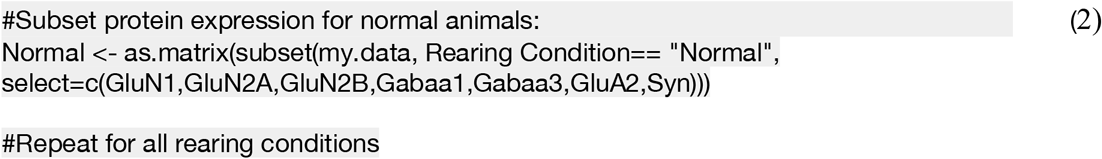

A correlation matrix was calculated for each subset of data. There are many packages in R that can calculate Pearson’s R correlations. We used the *rcorr* function in the *Harell Miscellaneous* package (*Hmisc*, (20)) since it computes a matrix of Pearson’s R or Spearman’s Rho rank correlation coefficients for all possible pairs of columns of a matrix. The following coding example (3) calculates the Pearson’s R for the Normal subset of data:

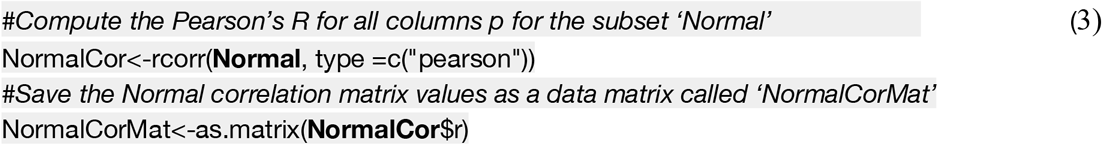

The matrix of Pearson’s R correlations was then converted into distance values since those values will be used for the hierarchical clustering. The distance matrix is the inverse of the correlation matrix, so that values represent dissimilarity rather than similarity. This step is necessary for generating the dendrograms that will represent cluster hierarchies. The conversion from a correlation matrix into a distance matrix was done in R using the *dist* function in the *stats* package (21).

This coding example (4) converts the correlation matrix for Normal animals (`NormalCorMat`) to a distance matrix:

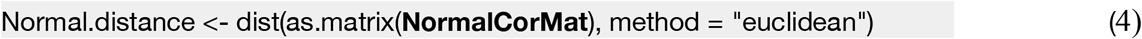

The distance matrix for the proteins was used was used for hierarchical clustering with the *hclust* function in the *stats* package. Clusters were visualized using a dendrogram to show the branching pattern and distances between proteins. Proteins with similar patterns of correlations were separated by shorter branch distances (y-axis) and fewer branch points.

This coding example (5) calculated the hierarchical clustering of the `Normal.distance` matrix and generated the dendrogram in Figure 5:

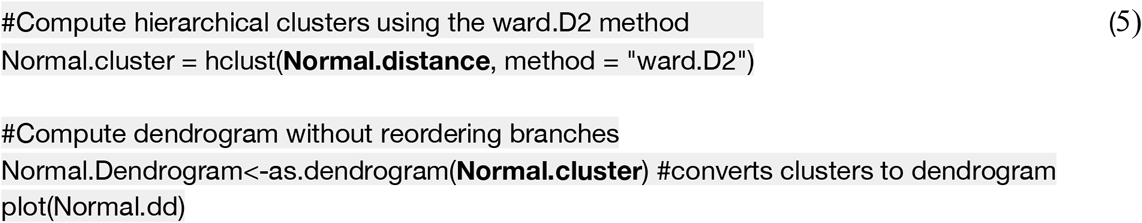

**Figure 5.**
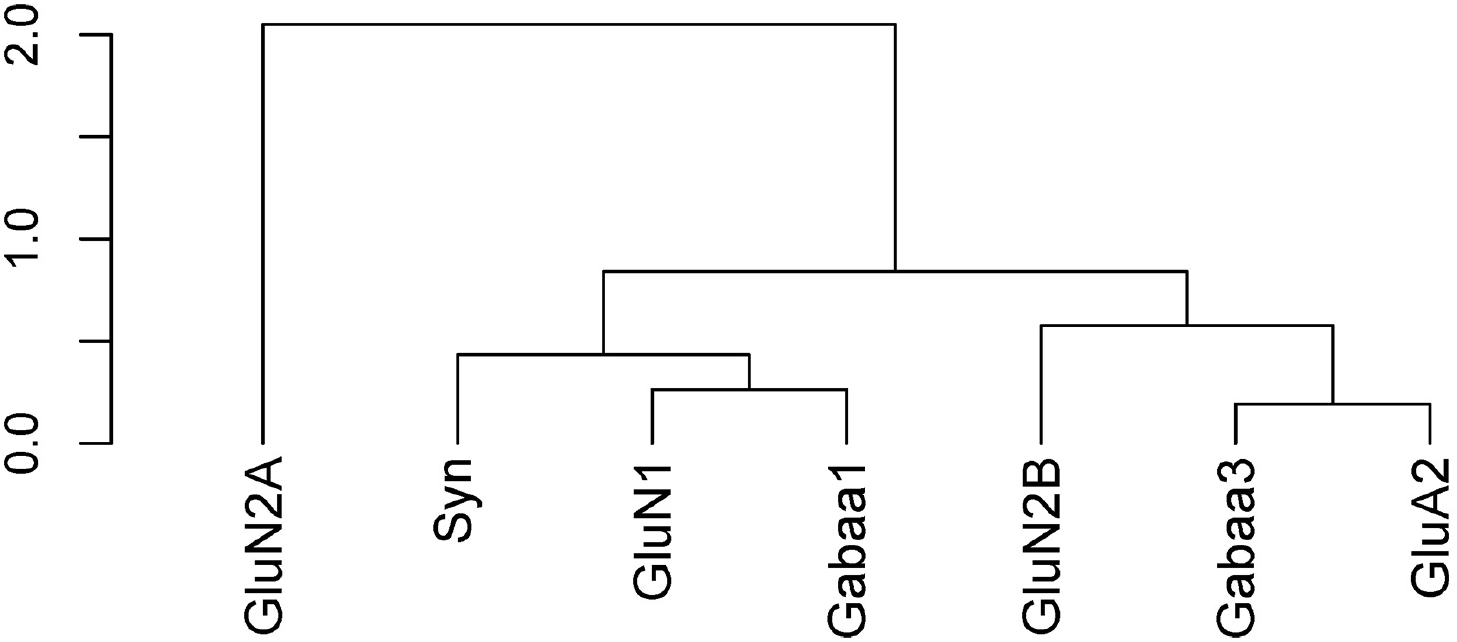
Initial output of hierarchical clustering the Normal distance matrix of proteins. The distance (y-axis) represents how closely related proteins are, with smaller values indicating a smaller distance and therefore a higher correlation. In this dendrogram, GluN2A is the most dissimilar to all other proteins. Depending on the height of the cut level, this dendrogram can be parse into any number of clusters (*k*).

The last step was to create the colour-coded correlation matrix with surrounding dendrogram in Figure 6 using the *heatmap.2* function from the *gplots* package (22). The correlation matrix (`NormalCorMat`) and dendrogram (`NormalDendrogram`) were the inputs for this example code (6). The colour scheme of the correlation matrix was adjusted by selecting an appropriate colour palette using the ‘col’ parameter, and the limits of the correlation matrix were adjusted to represent the range of Pearson’s R correlations using the ‘breaks’ parameter.

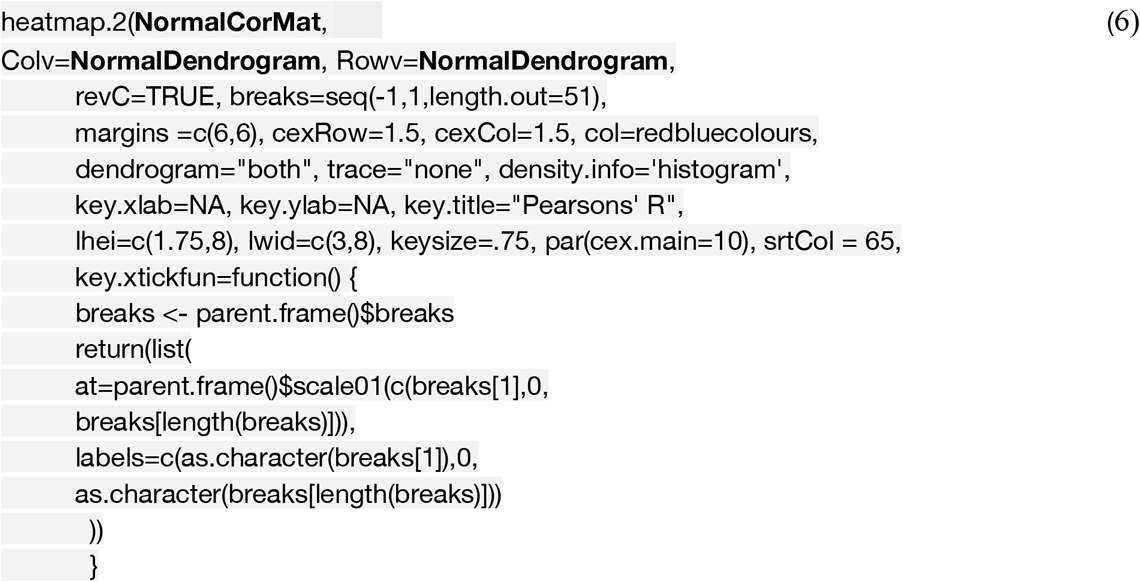

**Figure 6.**
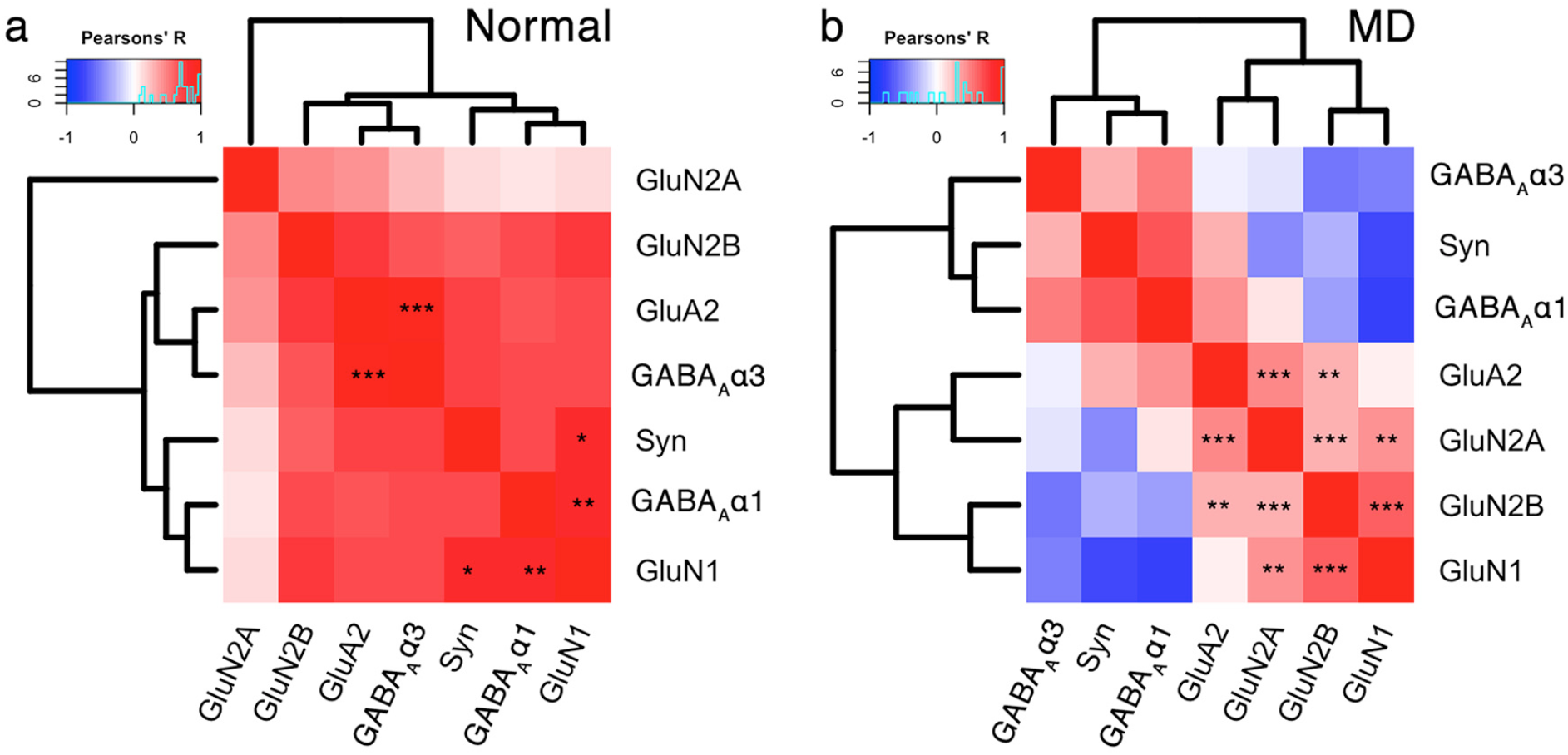
Correlation matrices for protein expression in Normal and MD animals. **a.** The output of the coding example for the Normal animals reveals most proteins have positive correlations with each other. **b**. The same code was used on the subset of MD data to create a correlation matrix, distance matrix and dendrogram. The MD plot shows that the excitatory proteins are positively correlated with one another, the inhibitory are positively correlated with one another, but excitatory proteins are negatively correlated with inhibitory proteins.

The examples above illustrate the information that can be readily visualized in a 2D correlation heatmaps. This analysis helped to identify the pairs of proteins (e.g., GluN1 and GABA_A_α1) that had different relationships after abnormal visual experience.

A final note: the dendrogram package *dendextend* (23)provides additional control for dendrogram attributes such as line style, thickness, and colour. Also, the *seriation* package (24) allows rotation of child branches to improve visualization. For example, in the MD heatmap above the first branch separates positive and negative correlations and using the *seriation* package that branch could be rotated so the blue cells are on the top left and the red cells on the top right. This control can be helpful for highlighting the pattern of correlations for particular proteins in the study.

### 3.iii) Dimension reduction

The next part of the analysis workflow uses PCA to explore the high-dimensional nature of the data. We have implemented a two-step procedure that starts by reducing the dimensionality of the data and then uses the basis vectors for those dimensions to identify the biological features that capture the variance in the data.

A note of caution: many implementations of PCA do not work well when there are empty cells in the matrix. There are a variety of approaches that can be used, including imputation to fill in the empty cells, removing runs where data are missing for one or more proteins, or averaging across runs. In this example, we averaged protein expression across runs.

This section is not an overview of PCA and we encourage readers to go to the online tutorials to learn more about applying PCA to biological data. It is important, however, to emphasize that our use of PCA is a data-driven approach to protein analysis because the variables (*p*) were *only* protein expression and did not include any of the categorical information such as treatment condition, cortical area or age.

This coding example (7) starts by extracting the columns (3-9) from `my.data` that contained protein expression.

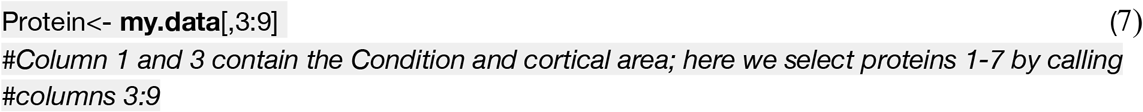

The first step for performing PCA was to center and scale the data so that proteins with abundant expression did not obscure proteins with smaller but still significant variations in expression. Each protein was scaled and centered producing a standard deviation of ±1, and a mean of zero. Scaling data in R was done using the following example (8) of the base function *scale*:

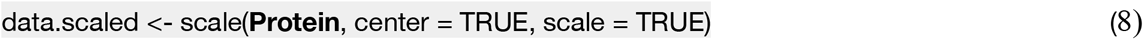

There are a variety of packages in R to do PCA, and here we used the *PCA* function in the *FactoMineR* package (25). That package produces eigenvalues and comes with excellent visualization tools to aid exploration of the relationships between principal components and features in the data set.

First, we ran PCA on the scaled data set `data.scaled` and saved the result as the object `pca`.

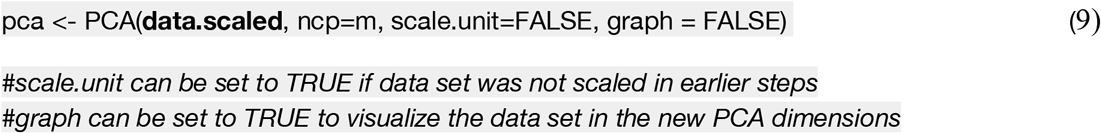

Principal components (PC) returned by this function are the set of orthogonal vectors in the object pca identifying the variance in data.scaled. The eigenvalues represent the magnitude of the variance captured by each PC vector, and the eigenvalue is largest for PC1 and successively less for each subsequent PC. An in-depth explanation of PCA and eigenvectors can be found here (6).

The eigenvalues for each PC were identified by consulting the `pca` object as follows:

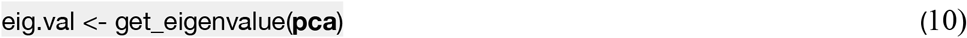

The first step of dimension reduction was to identify how much variance was captured by each PC, then rank the PCs from largest to smallest, and lastly, retain the set of PCs that capture a significant amount of the variance. Start with a Scree plot (Figure 7) showing the amount of variance explained by each of the PC dimensions. The following coding example (11) demonstrates how to consult the pca object to create a scree plot.

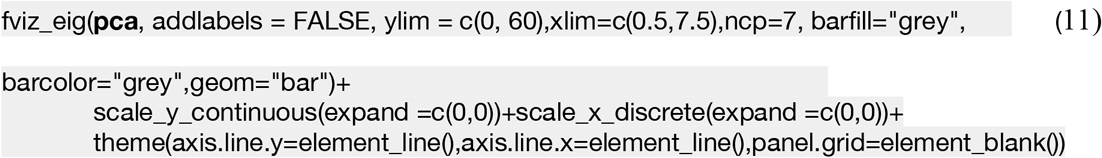

**Figure 7.**
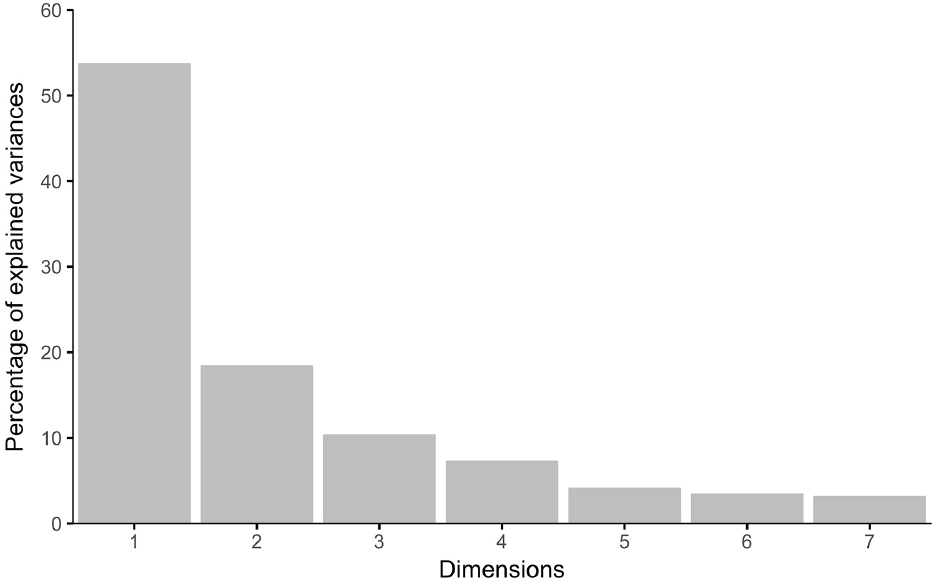
Scree plot of the percentage of explained variance captured by each principal component. The amount of cumulative explained variance over the first 3 dimensions is >80%. Typically, the first 3 dimensions are used at the cutoff value for reducing dimensionality. The remaining 4 dimensions each contain <7% variance.

The Scree plot (Figure 7) shows the decreasing magnitude for the variance explained by the 7 PC vectors. A variety of methods have been used to identify the significant dimensions (26,27) and here we used the simple rule to retain successive dimensions until the amount of variance explained was ≧80%. In this example, Dim1-3 explained 82% of the variance.

Once the significant dimensions were identified they were used to select candidate plasticity features driving the variance of each dimension. The 3 significant PC vectors can be represented by the weighted contribution from each the 7 proteins that together make up the basis vectors (Figure 8). Those were used to understand which proteins drove the variance in the data. That information was stored as XY coordinates in the `pca` object and it was called with the following coding example (12).

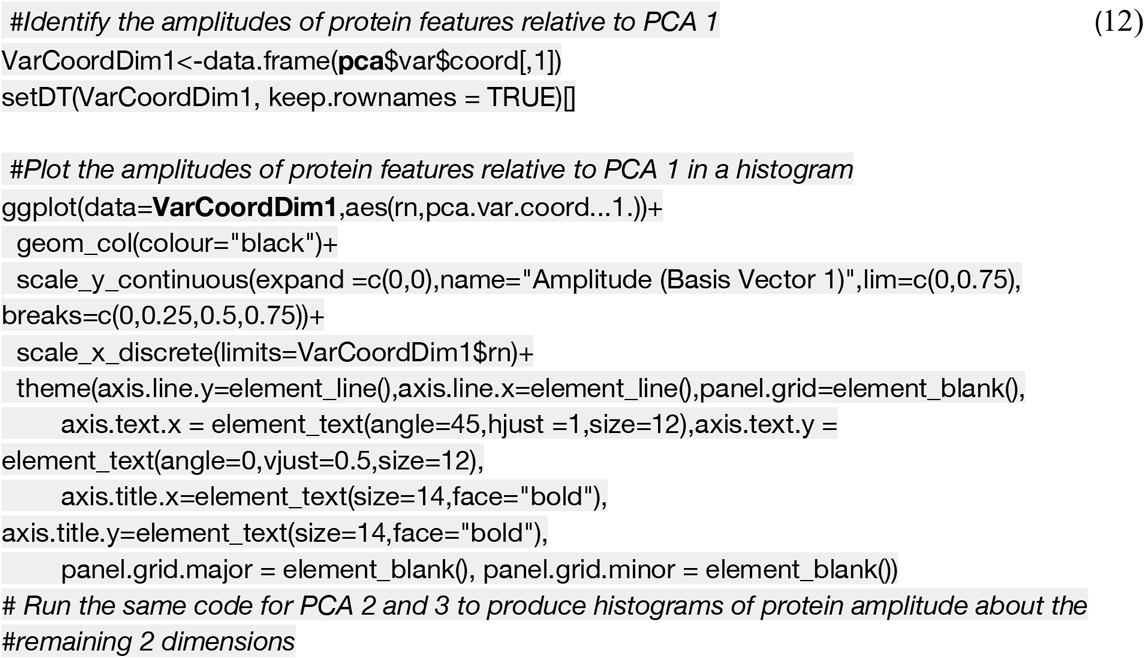

**Figure 8.**
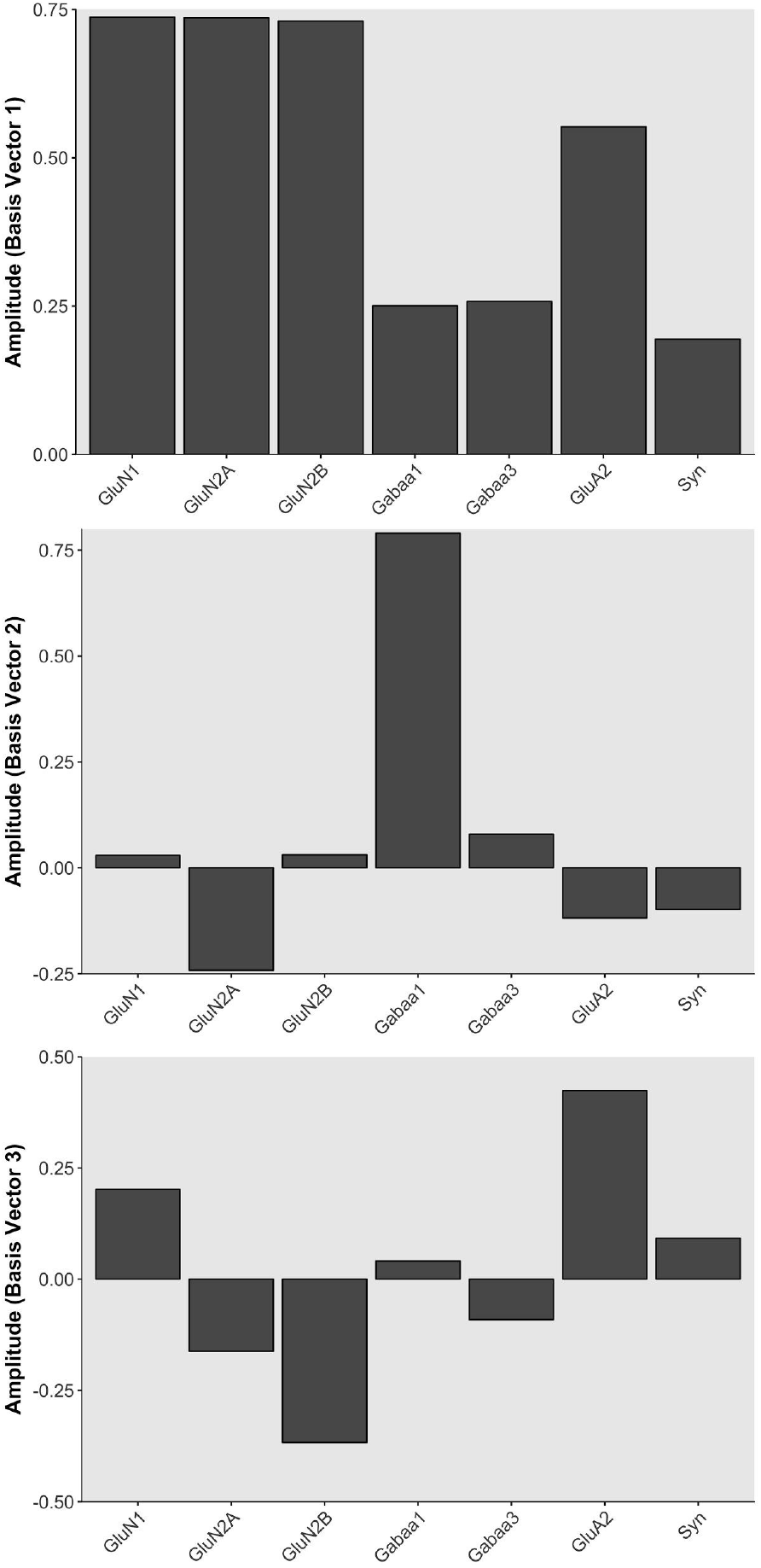
The basis vectors for dimensions 1-3 showing the protein vector amplitudes. The vectors were used to identify candidate plasticity features (top). For example, the basis vector for PCA 1 showed that all proteins move in the same direction suggesting that this dimension reflects protein sums (middle). The basis vector for PCA2 showed that GluN2A and GABA_A_α1 contribute in opposite directions, suggesting that this dimension reflects a balance between those proteins (bottom). The basis vector for PCA3 reflected a balance between GluN2B and GluA2.

We used multiple steps to identify candidate plasticity features from each of the basis vectors. Starting with PC1, we noticed that the weights for all the proteins were positive, so 3 features candidate were made from the sum of all proteins, the sum of the glutamatergic proteins, and the sum of the GABAergic proteins. Next PC2 and PC3 were inspected, these basis vectors had both positive and negative weights suggesting that along these dimensions the expression of some proteins increased while others decreased. This inspection revealed some pairs of proteins (e.g., GluN2A:GluN2B) that are known to change in opposite directions with different forms of visual experience. This step also identified novel pairings (e.g., GABA_A_α1:GluN2A) that were also included as candidate features. Continuing this approach, we identified 9 candidate features from the 3 basis vectors, and it is important to note that all were combinations or proteins rather than individual proteins. Thus, this approach to using PCA analysis can be described as an initial dimension reduction followed by expansion into candidate plasticity features. Importantly, the expansion steps will identify both novel features and ones that have been well studied thereby facilitating interpretation of the results within a biologically relevant framework.

### 3.iv) Feature selection

The features were validated by determining the correlation between each of the 9 candidate features and the 3 dimensions. This was done by calculating the 9 candidate features for all of the samples using the protein expression data and correlating those with the eigenvalues for the 3 dimensions. Bonferroni correction was done to adjust the significance level for the multiple correlations and features that were significantly correlated with a dimension became the plasticity features used in subsequent stages of the workflow.

The validation of candidate features was done in R, by storing the new features in a matrix `NewFeatures`, centering and scaling those data, then correlating the eigenvalues with the `NewFeatures` matrix. The function *corr.test* from the *psych* package (28) was used for that step. The strength of the significant correlations was visualized with a custom 2D matrix created using the *geom_tile* function from the *gplots* package (22).

The new features were centered, scaled and stored as a new data matrix:

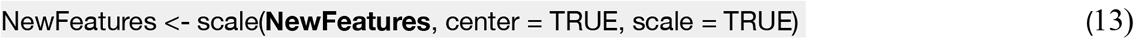

Next, the coordinates for all data points in PCA space were stored in another matrix by consulting the `pca` object as follows:

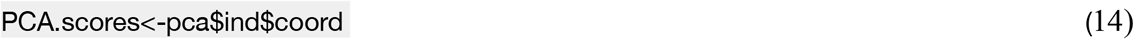

Finally, the correlations between the two matrices `PCA.scores` and `NewFeatures` were determined and visualized using the following R coding example (15):

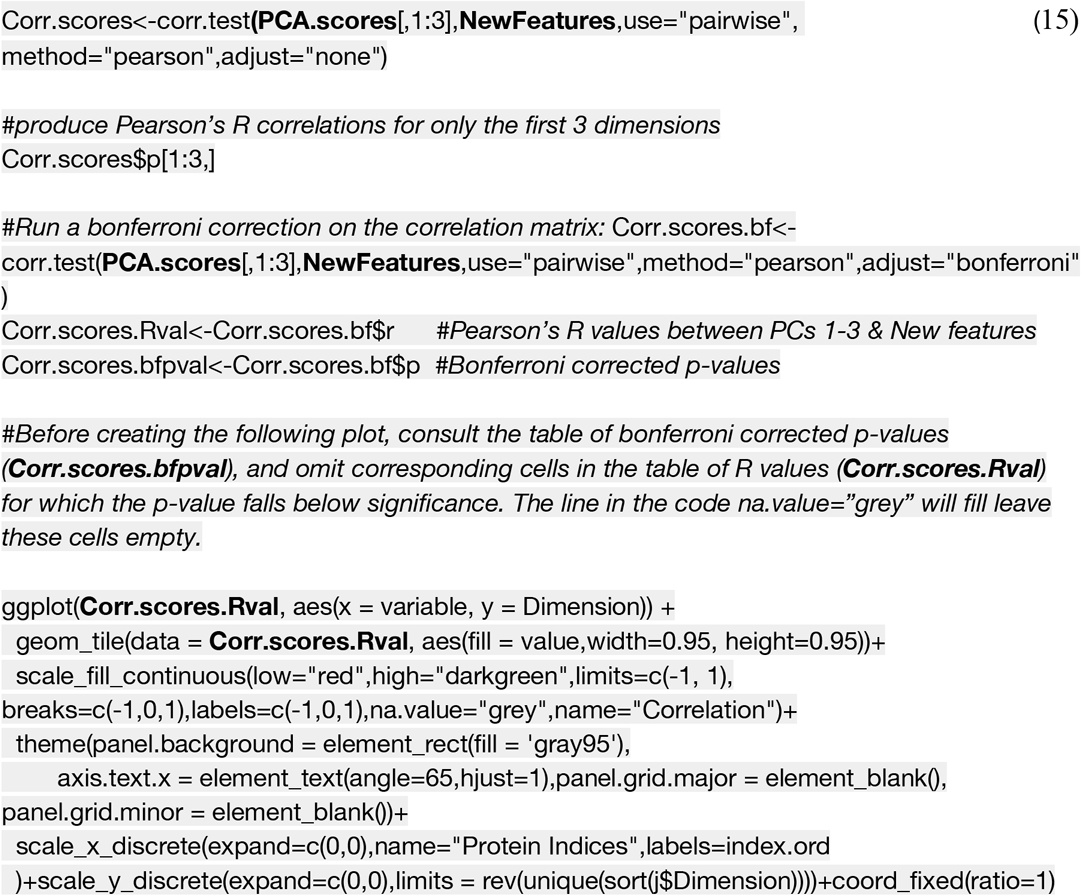

**Figure 9.**
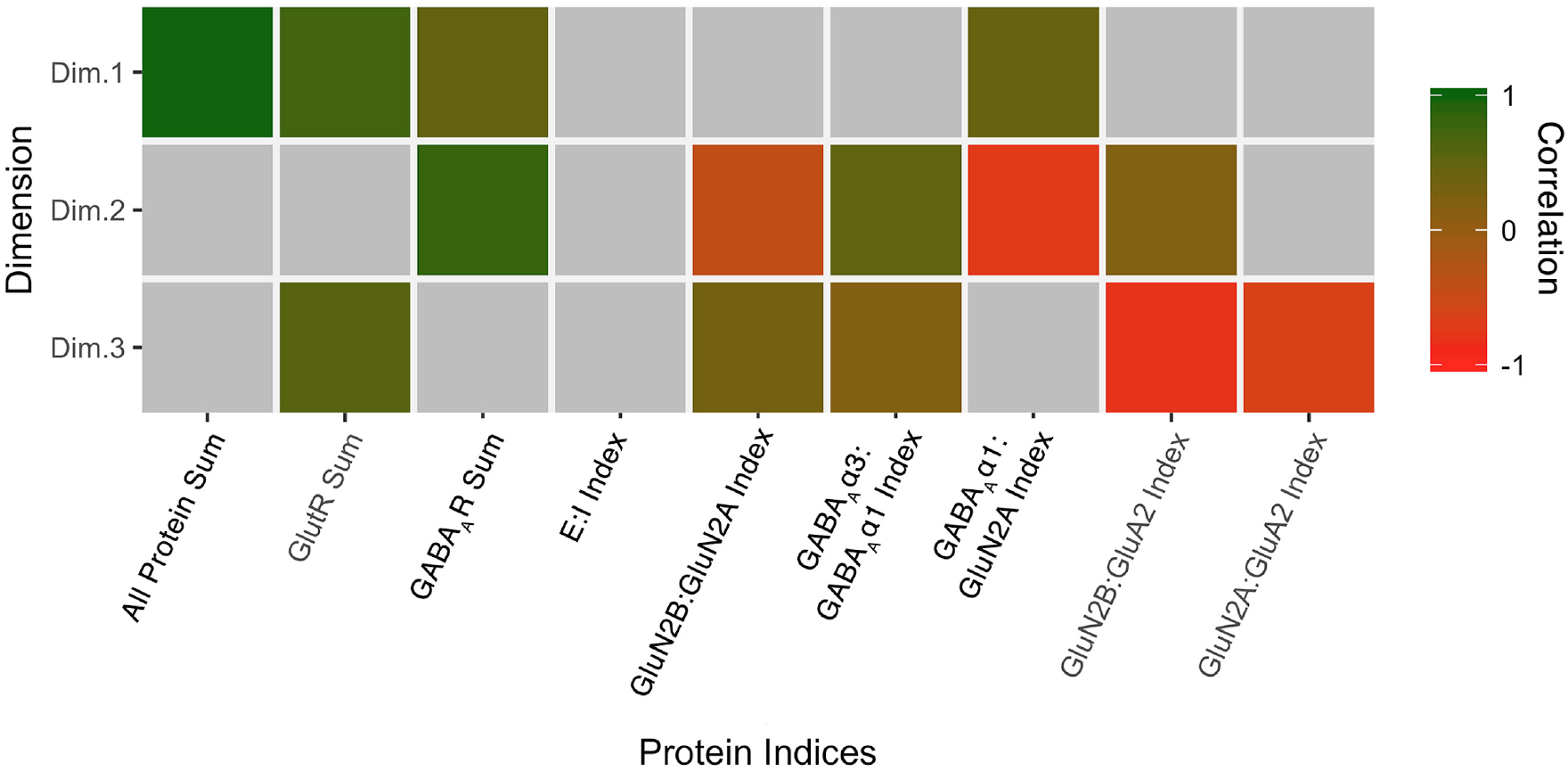
Correlation between the plasticity features (columns) identified using the basis vectors for PCA dimensions 1-3. Coloured cells are significant, grey cells are non-significant. Bonferroni corrected correlations (green = positive, red = negative).

The plot of correlations between the 3 significant PCA dimensions and the candidate features was used to validate the selection of features that will used as the input to the next stage in the analysis workflow (tSNE and clustering). In the example above, all of the candidate features except a measure of the E:I balance was significantly correlated with at least one of the dimensions. Interestingly, none of the features were correlated with all 3 dimensions demonstrating that multiple plasticity features are necessary to capture the variance in the data.

The whole collection of plasticity features was combined to form the plasticity phenotype that became the input to the next step in the analysis workflow.

A final note: the new plasticity features can be analyzed using univariate statistics to determine if there are significant differences among rearing conditions. That analysis is particularly important when feature selection identifies combinations of proteins that are not typically studied since they may provide new insights into the neurobiological mechanisms that differentiate among groups reared with various forms of visual experience.

### 3.v) tSNE and clustering

This step used tSNE analysis to preserve both the global and local arrangement of in the plasticity features. Also, tSNE revealed good clusters were revealed because of the artificial scaling of the distance between data points with similar patterns of features.

In this example, the inputs to tSNE were the validated plasticity features (`NewFeatures`) without any of the information that identified the source of the sample (e.g., cortical region, rearing condition, or age).

The following coding example (16) demonstrates how to perform a tSNE analysis using the *tsne* function from the *tsne* package (29).

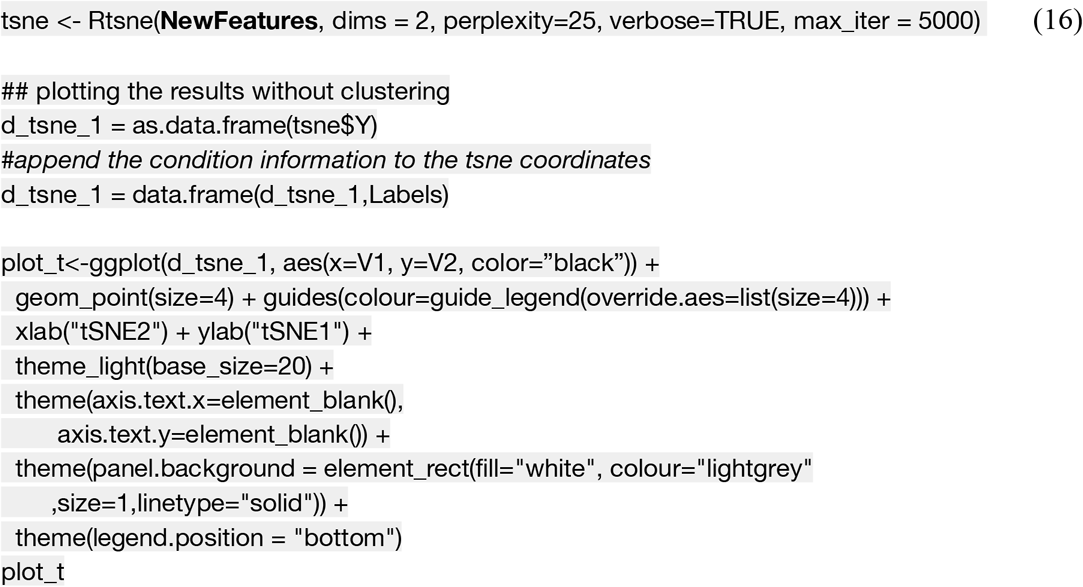

The first step in the tSNE analysis reduced the plasticity features from each sample to tSNE XY coordinates (Figure 10). Those coordinates were used as the input to *K*-means clustering analysis to identify and then visualize clusters in the data set.

**Figure 10.**
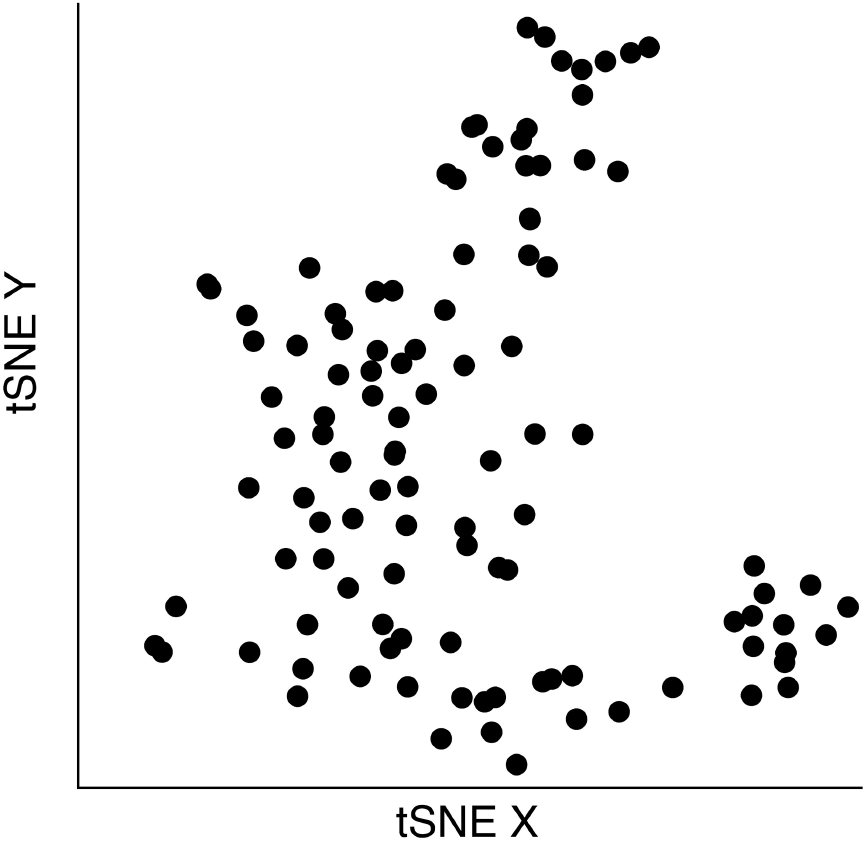
tSNE output when run on NewFeatures identified by PCA. The dimensions for X and Y in tSNE have no units, but the distance between objects is scaled to position similar samples nearest to one another.

Both *K*-means and hierarchical clustering algorithms require the number of clusters *k* as a parameter. A good method for choosing the number of clusters is to measure the within groups sum of squares (WSS) for a range of *k*, plot that information and then determine the inflection point.

In the example data set there were 9 rearing conditions (e.g. normal, monocular deprivation etc.) so we chose a range for *k*, 2 to 15 clusters, that encompassed the number of conditions.

The following coding example (17) determined the WSS and plotted it for the *k* clusters:

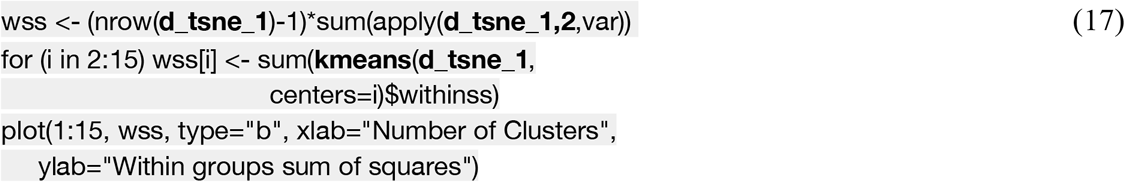

**Figure 11.**
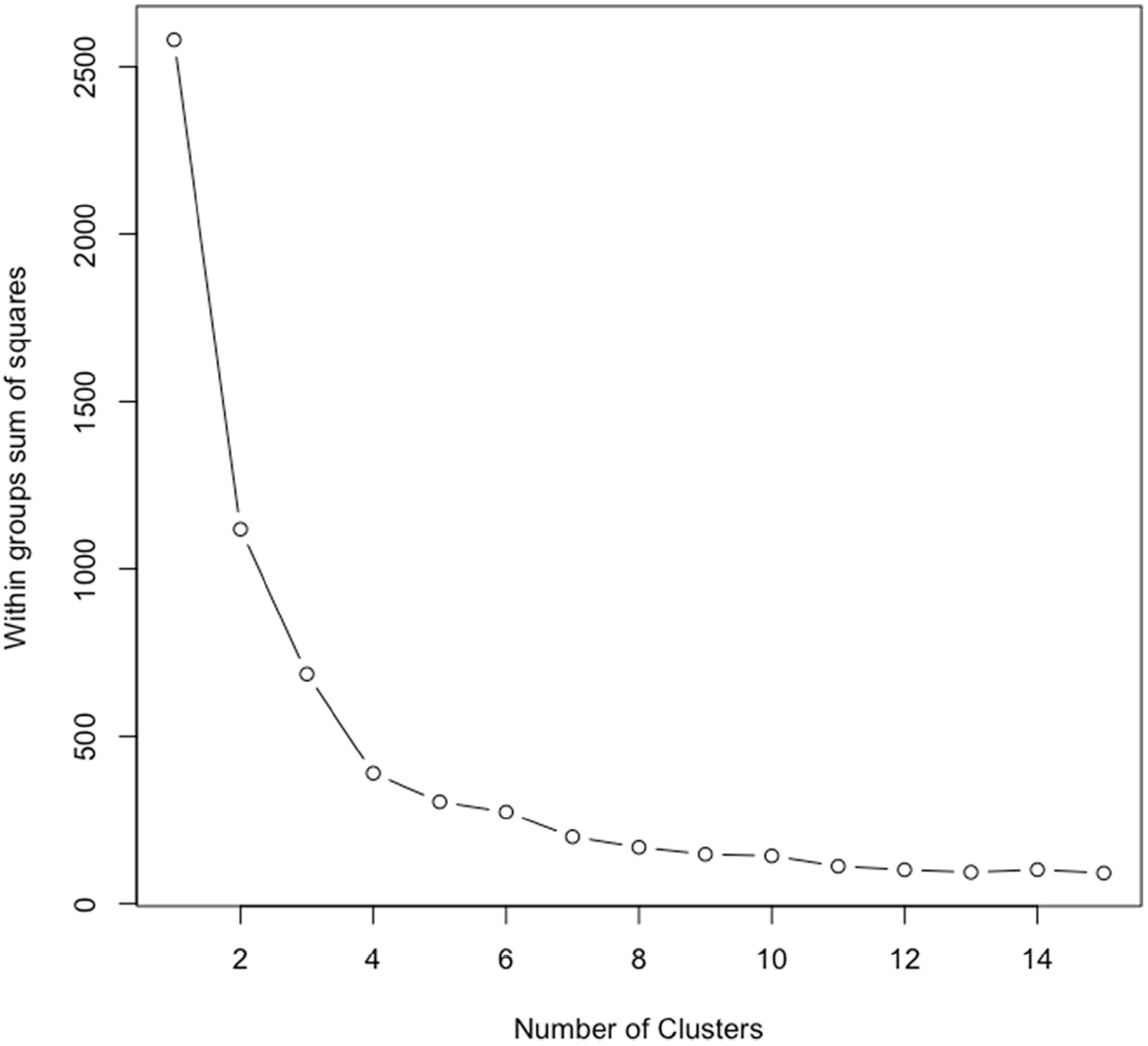
Within group sum of squares for hierarchical clustering of tSNE coordinates in Figure 9. Sum of squares decreases as cluster number (*k*) increases. Increasing k has little effect on sum of squares after a certain threshold is met. In this plot it appears that 6 or 7 clusters represents that threshold point. To accurately identify optimal cluster number using this “elbow method” we apply an exponential decay function to the curve and identify the 3τ, the point where 85% of the change has occurred.

The optimal number of clusters was selected by fitting an exponential decay curve to the WSS data the finding the number of clusters corresponding to point where the curve plateaued (4τ) (*k*=6). This approach is called the “elbow method”, where 4τ is the point of inflection, or elbow, of the curve.

Next, *K*-means clustering for *k*=6 was done on the output from the tSNE analysis (Figure 12a). The clusters were assigned different colours to visualize the samples in each cluster. Some clusters (green and yellow) were spatially separated on the tSNE plot, while others (e.g., orange and blue) were adjacent. The following coding example (18) plots the clusters identified in the tSNE representation of the data as different colors, but these colors can be manipulated to represent other characteristics of the data (e.g. cortical area, treatment condition).

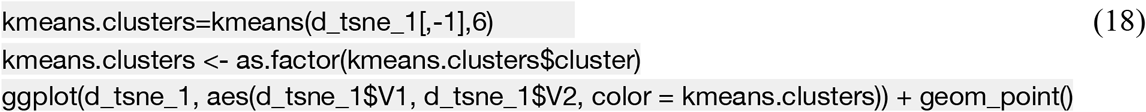

**Figure 12.**
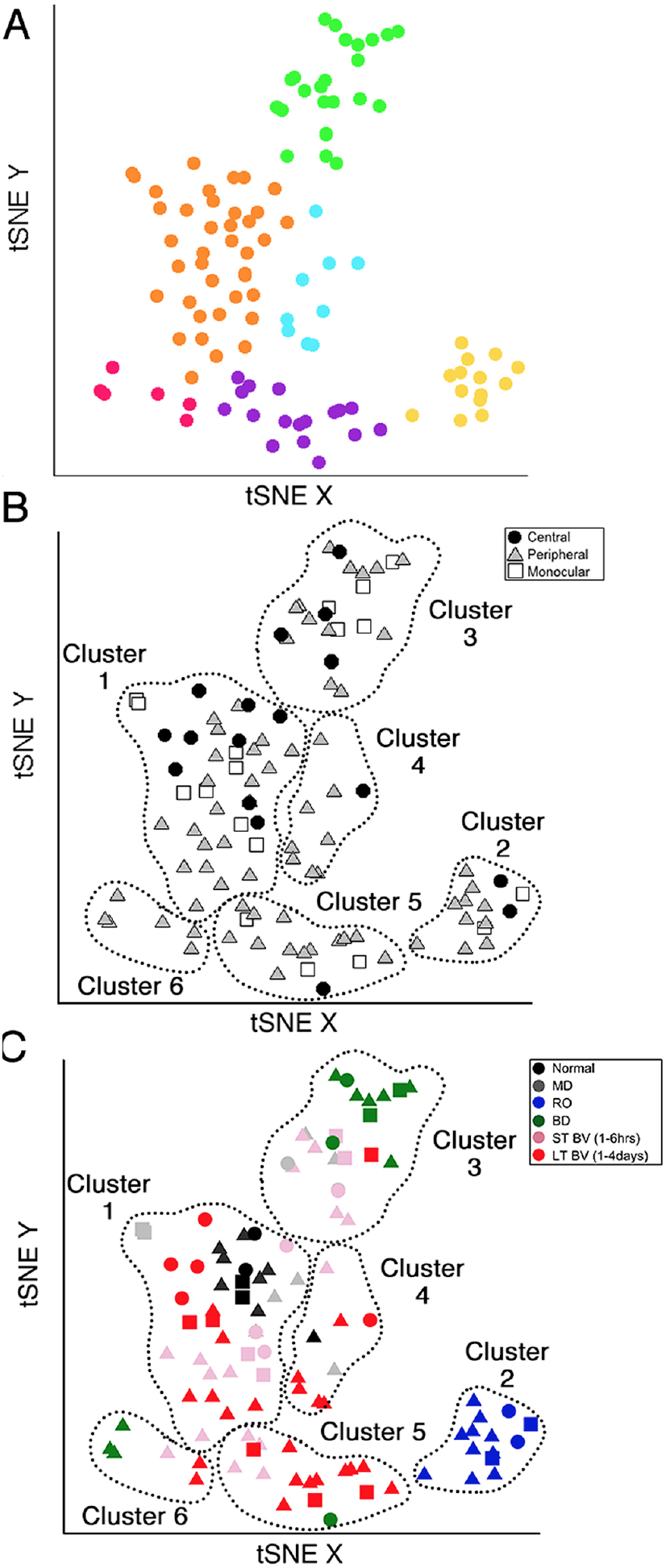
*K*-means clustering applied to tSNE reduced data. **a**. The clusters were colour-coded to visualize the 6 clusters identified in the data. **b**. Samples were annotated for cortical region (black circles=central, grey triangles=peripheral, white squares=monocular) and dashed lines were drawn around the clusters identified in a. **c**. Samples were colour-coded according to the rearing condition, the shapes were the same as in b.

The number of samples in each cluster ranged from 5 (magenta) to 38 samples (orange). We annotated each sample based on the visual cortical region (central, peripheral, or monocular) (Figure 12b) and rearing condition (Figure 12c) to analyze cluster composition and determine if the clustering reflected one of those parameters. For example, cluster 2 contained samples from only one rearing condition (reverse occlusion) and cluster 1 contained almost all of the normally reared cases but it also had samples from other rearing conditions. Thus, this step identified clusters and provided some evidence that the rearing condition was driving changes in the plasticity phenotypes. The tSNE clustering, however, did not reveal which features from the phenotypes were separating the samples into different clusters or grouping them into the same cluster.

Annotating each sample by the cluster, visual cortical region, and rearing condition was an essential first step for using the plasticity phenotypes to identify and explore *subclusters* in the data. That process identified 13 subclusters in the example data set (Table 3).

**Table 3.** Subclusters identified from the tSNE plot

| Subcluster | Rearing Condition | tSNE Cluster | Region |  |  |
| --- | --- | --- | --- | --- | --- |
|  |  |  | C | P | M |
| Normal 1<sub>C,P,M</sub> | Normal | 1 | X | X | X |
| LTBV1<sub>C,P,M</sub> | Long term BV recovery | 1 | X | X | X |
| MD1<sub>P,M</sub> | Monocular deprivation | 1 |  | X | X |
| STBV1<sub>C,P,M</sub> | Short term BV recovery | 1 | X | X | X |
| RO2<sub>C,P,M</sub> | Reverse occlusion | 2 | X | X | X |
| STBV3<sub>C,P,M</sub> | Short term BV recovery | 3 | X | X | X |
| MD3<sub>C,P</sub> | Monocular deprivation | 3 | X | X |  |
| BD3<sub>C,P,M</sub> | Binocular deprivation | 3 | X | X | X |
| LTBV4<sub>P,M</sub> | Long term BV recovery | 4 |  | X | X |
| LTBV5<sub>P,M</sub> | Long term BV recovery | 5 |  | X | X |
| STBV5<sub>P</sub> | Short term BV recovery | 5 |  | X |  |
| LTBV6<sub>P</sub> | Long term BV recovery | 6 |  | X |  |
| BD6<sub>P</sub> | Binocular deprivation | 6 |  | X |  |

A final note: In this workflow, dimensionality reduction and feature selection were performed before tSNE analysis and clustering. Although this is a common approach for analyzing high-dimensional data in neuroscience it is important to remember that PCA preserves the features with variance that is aligned with the orthogonal dimensions. Thus, features with more subtle but important variance away from the PCA dimensions will not be included in subsequent clustering (9). We will return to this issue in the section 4 where we present a second analysis workflow to handle data with more features.

### 3.vi) Identifying and exploring subclusters

In this section, we describe how to analyze and visualize subclusters using the features that comprise the plasticity phenotypes.

First, the features and tSNE results were combined in R by appending the object containing the tSNE dimensions and clusters (`d_tsne_1`) to the plasticity features (`NewFeatures`). Now each sample had both the clustering information from the tSNE analysis and the feature data from PCA. Next, the data were organized into subsets according to the subclusters. For example, all of the data points for Normal samples in cluster 1 were subset as follows:

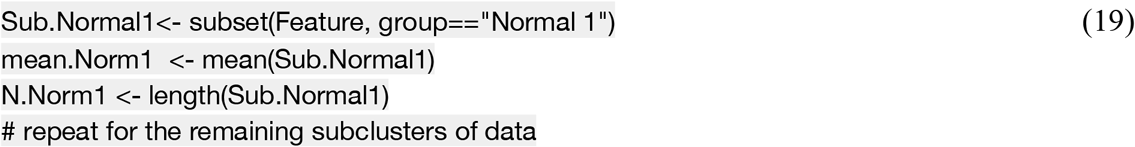

Now, univariate analyses were done to compare plasticity features between subclusters. Boxplots were made with the *boxplot* function in the *vioplot* package (30) and bootstrap tests were used to determine which subclusters were significantly different from the normal subcluster (Supplemental Information).

The significant deviations from normal were colour-coded in the boxplot to facilitate visualizing subclusters that had above (red) or below (blue) normal expression for a feature. To include the colour-coding of boxes a column with the information from the significance tests was added to the subset data (e.g., `Sub.Normal1`). For example, clusters that were significantly greater than `mean.Norm1` were identified with the label ‘red’, clusters that were significantly less than `mean.Norm1` with the the label ‘blue’, and those that were not significantly different than `mean.Norm1` with the label ‘grey’. That updated collection of subset data were stored in an object called `Clusters.Subsetted`, and the original cluster designation was stored as `Subset.Names`. The following coding example (20) was used to create boxplots of the subsetted data for the feature GluN2A:GABA_A_a1.

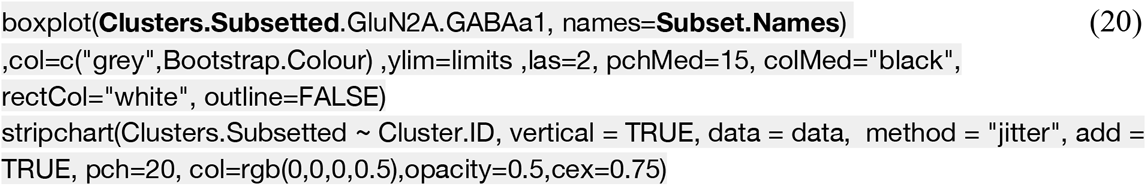

**Figure 13.**
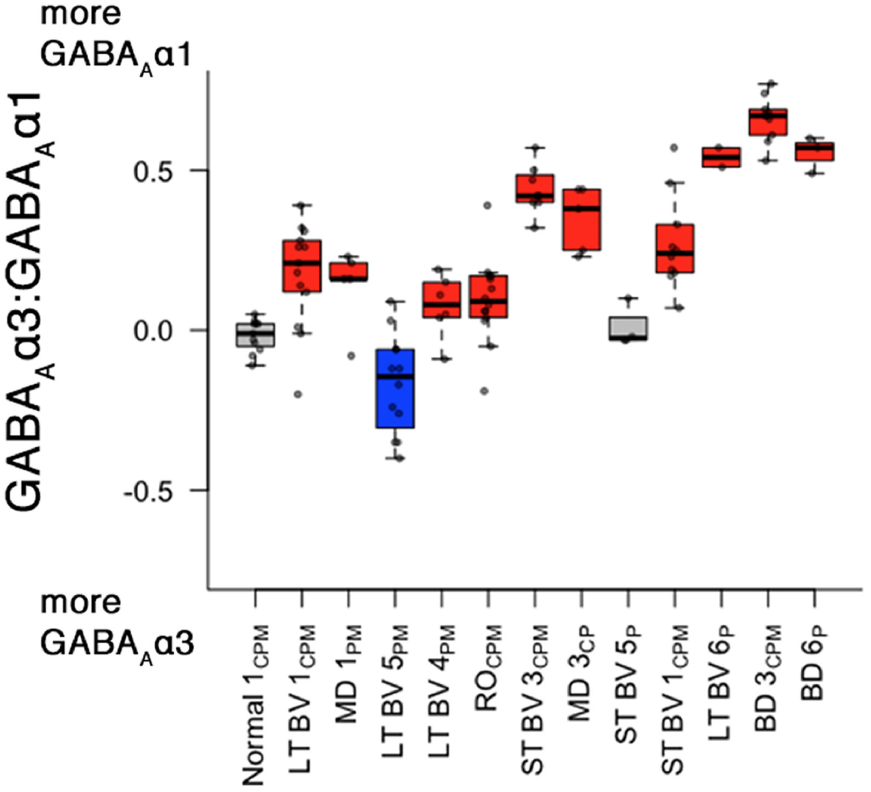
Boxplot for the GluN2A:GABA_A_α1 feature that identified subclusters with expression different from normal. The original 6 clusters were divided into 13 subclusters, annotated by the original conditions. Boxplots were drawn around the mean expression on the GluN2A:GABA_A_a1 balance. Boxes were coloured relative to normal (red for significantly above, and blue for significantly below). Scatterplots were drawn on top of each boxplot showing the observations within each cluster.

The boxplots were useful for highlighting significantly different subclusters for individual features. For example, the plot above identified 8 subclusters that were significantly greater than normal (red). However, it is daunting trying to synthesize all of the significant differences for 9 features and 13 subclusters using just that approach. Instead, we calculated the pairwise correlations between subclusters using the plasticity phenotypes, ordered the subclusters using hierarchical clustering and visualized these in a 2D heatmap. The steps were the same as explained in coding examples (2)-(6), except the input data for the plasticity phenotypes was `NewFeatures` and the 13 subclusters identified in `Clusters.Subsetted`.

**Figure 14.**
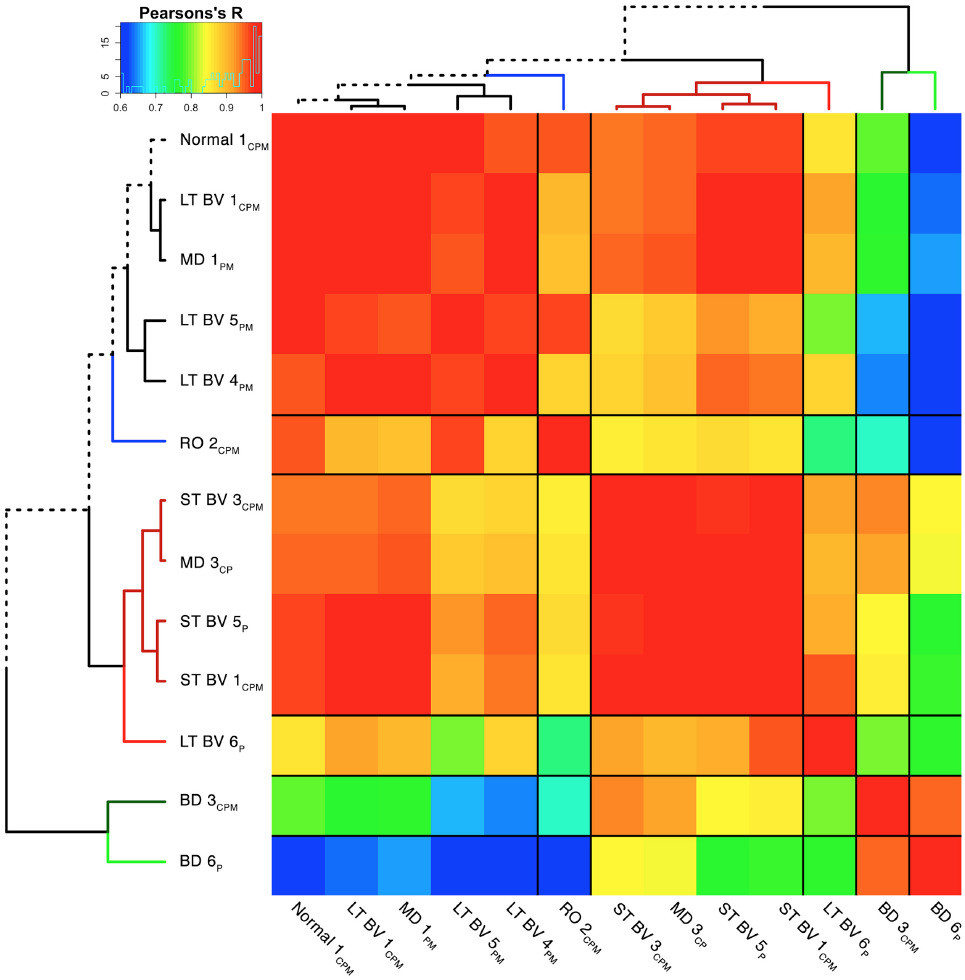
Correlation matrix for plasticity phenotypes between the subclusters of the data. Matrix of Pearson’s R correlations between the means across 8 plasticity features in each subcluster. Clusters were reordered according to the surrounding dendrogram. The dendrogram positioned similar clusters close together, and moved dissimilar samples to the periphery. Inset demonstrates counts across the range of Pearson’s correlations, while the color gradient ranges from low positive correlations (blue) to high positive correlations (red).

The correlation matrix for the plasticity phenotypes showed the strength of similarity or dissimilarity among the subclusters. Here, the surrounding dendrogram ordered subclusters for some rearing conditions (e.g., LTBV) on the same branch as the Normal subcluster, while other conditions (e.g., BD) were far from Normal branch. This analysis revealed which subclusters had similar plasticity phenotypes but did not clarify if that was based on the entire pattern of the features in the phenotype or if a smaller number of features drove the clustering.

### 3.vii) Construct and visualize plasticity phenotypes to identify similar and different features among subclusters

In the last step for this workflow we describe visualizing the plasticity phenotypes, ordering them using the dendrogram from the hierarchical clustering, and comparing phenotypes to identify differences among rearing conditions.

A display was created to show each of the feature and the whole pattern of the plasticity phenotype so that it was easy to compare the subclusters for similarities and differences visually (Figure 15). The visualization had a series of colour-coded horizontal bands where each band represented a feature, and together the 9 bands represented the average plasticity phenotype for a subcluster.

**Figure 15.**
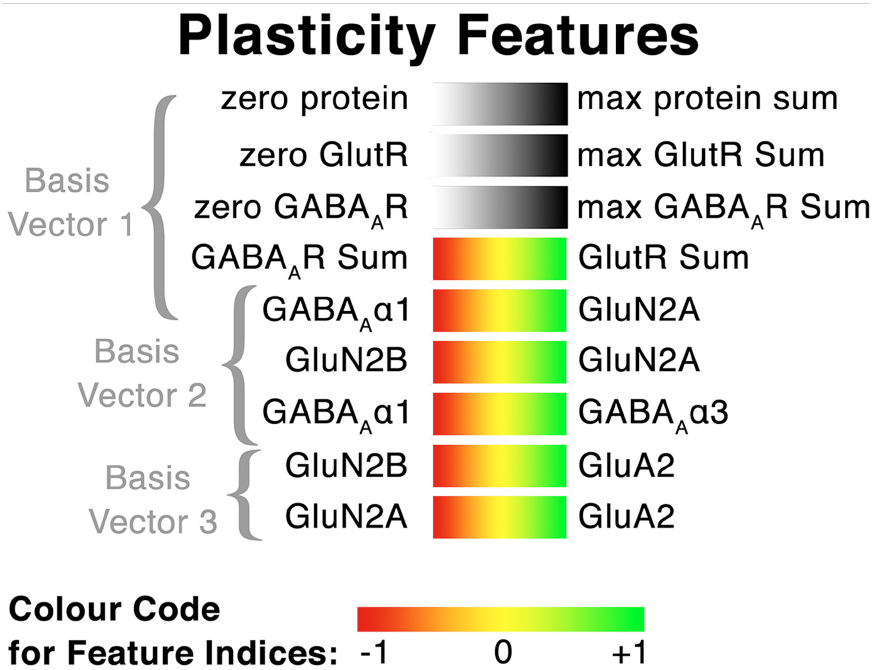
Legend for the features comprised in the plasticity phenotype and the colour-code. The top three bars (from protein amplitudes about PCA basis vector 1) represent protein sums and use greyscale (white to black) for zero to the maximum protein sum. The next 6 bars represent the feature indices identified with the basis vectors from PC2 and PC3 and use a colour-scale (red to yellow to green) for the shift from one protein to the other.

The plasticity phenotypes were visualized in R, using the *geom_tile* function in the *ggplot2* package (31). First, the feature mean was determined for each subcluster then the limits of the colour scales were set by finding the maximum and minimum expressions for a feature across all subclusters. Finally, the subcluster mean was converted to the corresponding RGB score. The following coding example (21) was used to map the mean for each feature in the Normal condition onto a color scale:

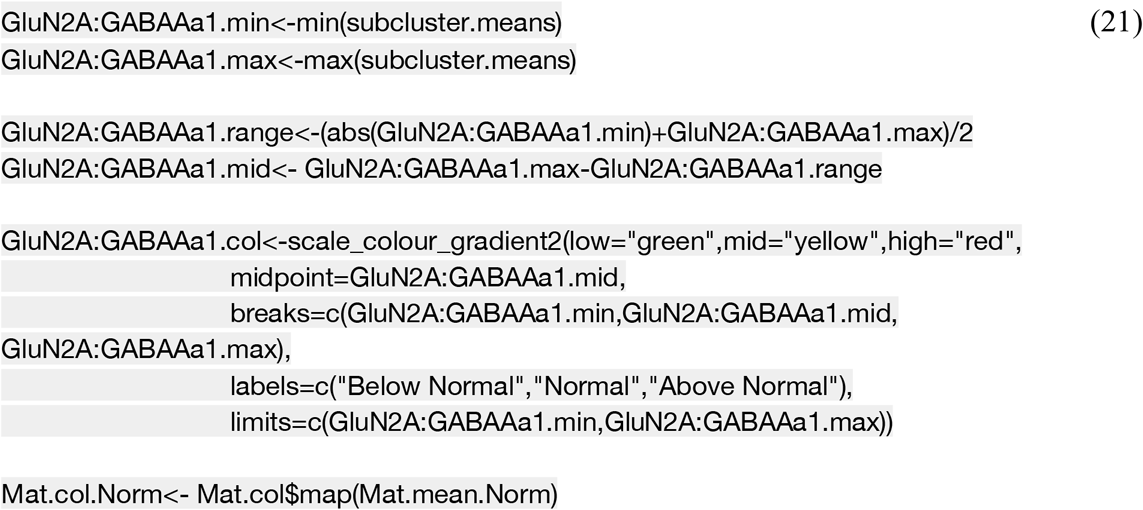

That list of colour-codes for each subcluster was stored in a new matrix called `Colour.Table`. The matrix will be consulted in the code below (22) to call the correct colour for each horizontal bar in the plasticity phenotype.

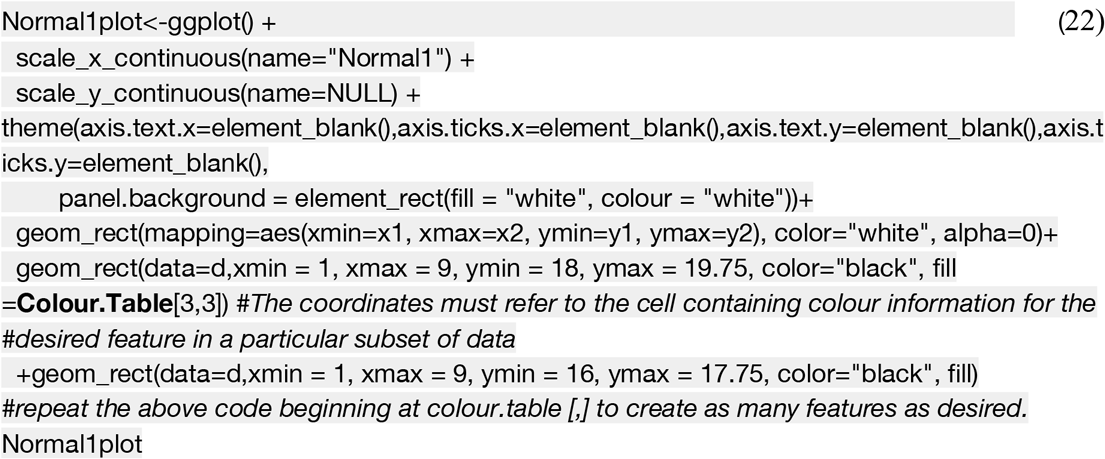

**Figure 16.**
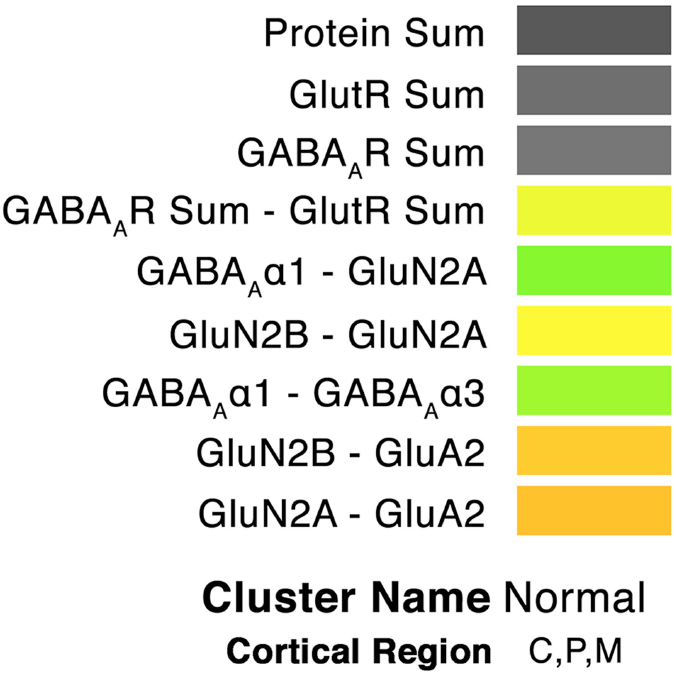
Example plasticity phenotype for the Normal subcluster. The top three bars depict high levels for the protein sums, while the next 6 bars represent the balance between the protein pairs for that feature using a green-to-red colour scale. The greenish bars indicate features that are biased toward first protein (e.g., GABA_A_a1 versus GluN2A), reddish bars are biased toward the second protein (e.g., GluN2A versus GluA2) and yellow bars reflect roughly equal expression of the two proteins (e.g., GluN2B versus GluN2A).

The figure below (Figure 17) illustrates the power of this tool for visualizing the patterns and features in the plasticity phenotypes that group or separate subclusters.

**Figure 17.**
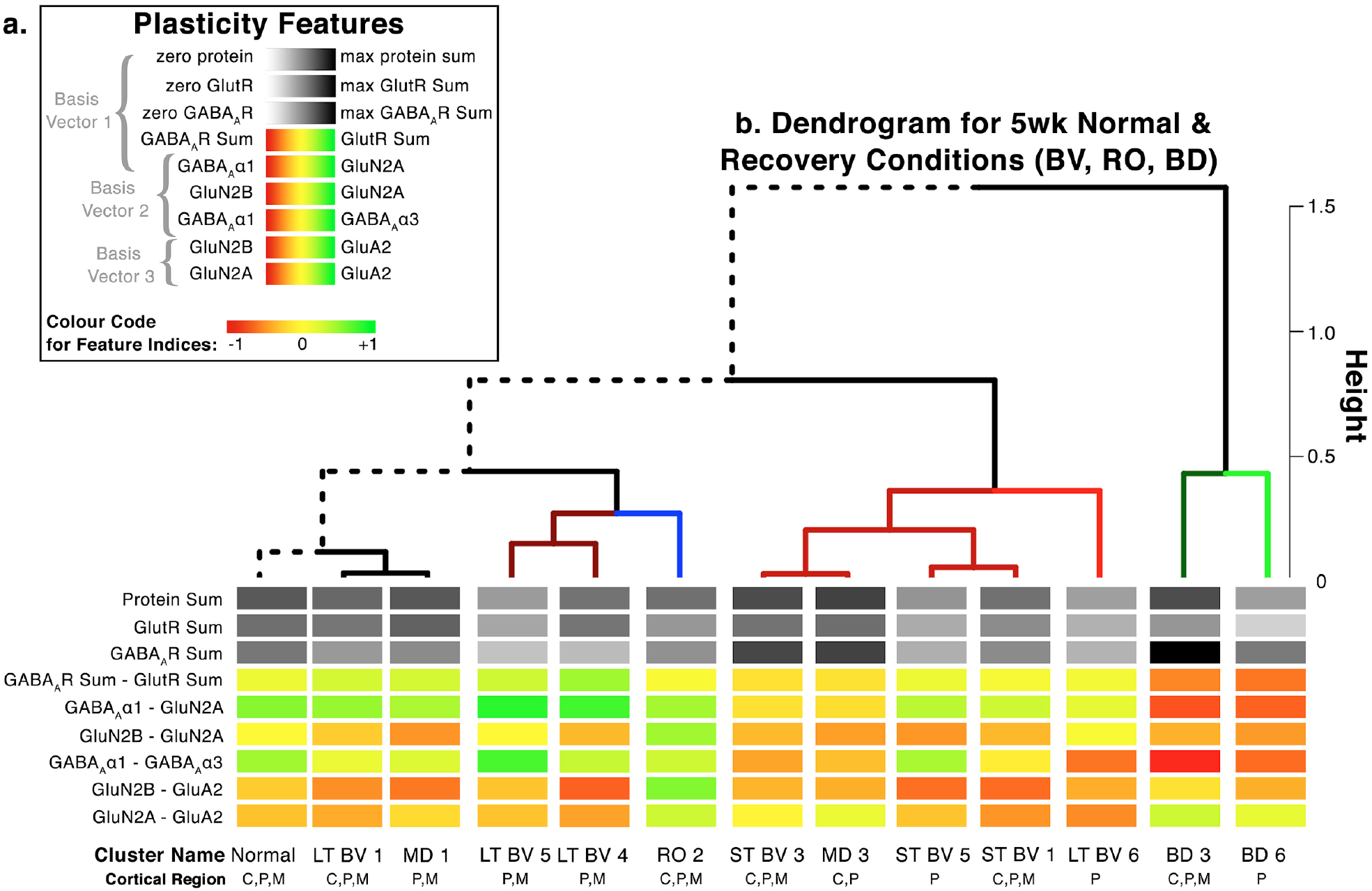
Example of the orderly progression of plasticity phenotypes across subclusters. The subclusters are noted below the phenotypes, and are arranged according to the dendrogram

The subclusters in this example were ordered using the same hierarchical clustering dendrogram as in Figure 14, and the average plasticity phenotype for a subcluster was displayed at the end of its branch in the tree. The figure provides a strong visual impression of the phenotypic similarity among subclusters located nearby in the tree (e.g., normal and LTBV) and differences for subclusters that are further away (e.g., normal and BD). Thus, this tool supports linking the output of high-dimensional analyses with neurobiologically meaningful insights. For the example data set, the visualization revealed the patterns of neural proteins changes driven by different forms of visual experience.

Finally, an alternative color scheme can be used to fill the horizontal bars in order to identify differences between the features in each phenotype. We performed a Bootstrap analysis comparing expression of each feature to the Normal subcluster of data points and colored each bar with a mean expression significantly greater than the normal subcluster red, and each bar with a mean expression significantly less than the normal cluster blue. Features that did not significantly differ from Normal subcluster were left empty. The result of that analysis is below.

**Figure 18.**
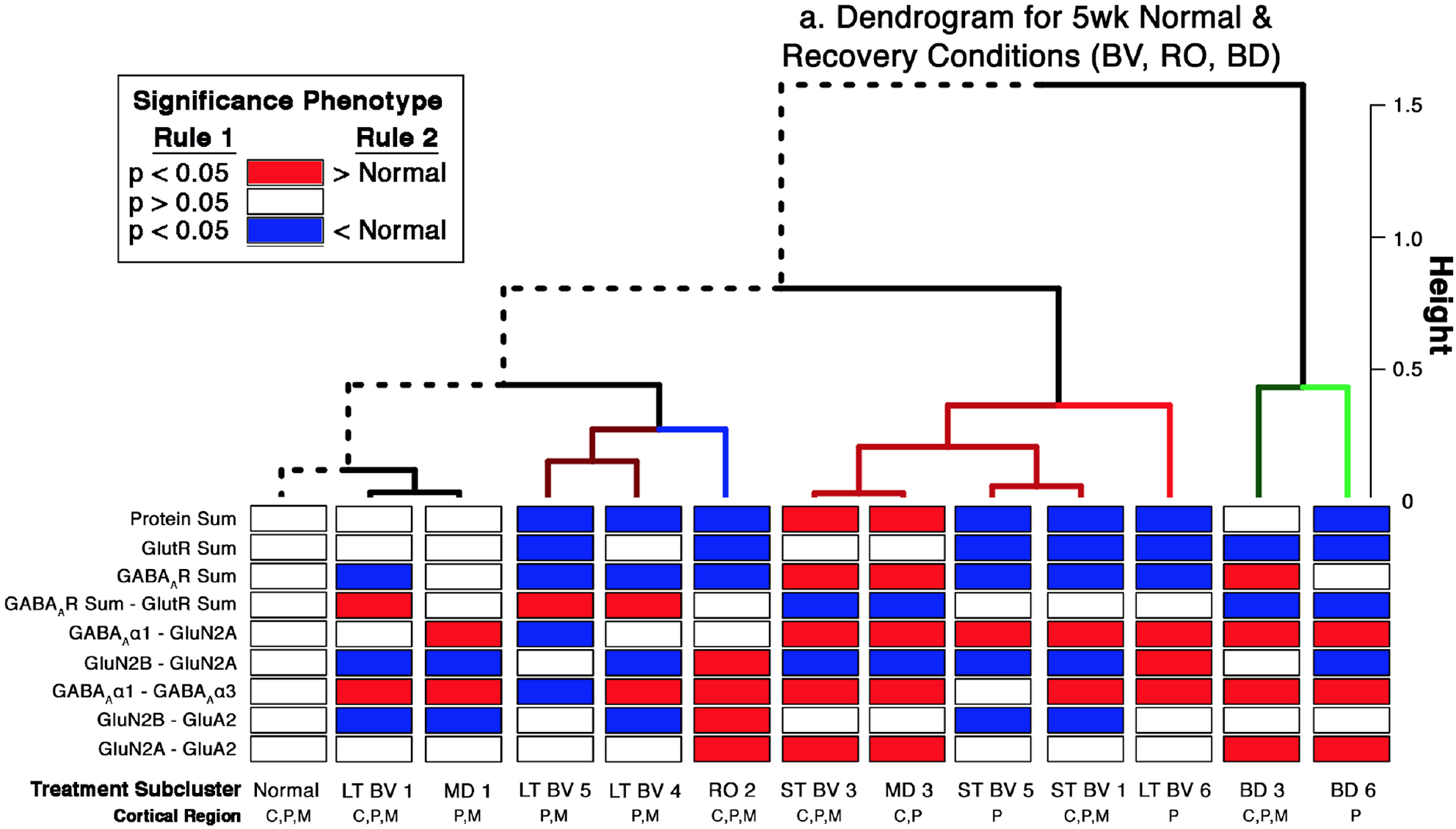
Identifying features of the plasticity phenotypes in the subclusters that are different from normal. The subclusters are noted below the phenotypes and are arranged according to the dendrogram. The features in each subcluster phenotype are color-coded to indicate if the feature is significantly greater than (red) or less than (blue) the 5wk normal condition (left most phenotype).

## 4. Using sparse high-dimensional clustering to study human visual cortical development: Clusters, features, and plasticity phenotypes

The goal of testing sparse high-dimensional clustering methods was to determine if they reveal age-related clusters in the expression of proteins in human visual cortex that provide meaningful insights into the neurobiological development of human cortex.

The second workflow describes the application of sparse high-dimensional clustering, transformation and identification of features using PCA, and exploration of cluster composition using plasticity phenotypes (Figure 19). This workflow moved away from using tSNE and instead applies sparse high-dimensional clustering for analyzing the protein data. The data set used here was from a series of studies examining the development of synaptic and nonsynaptic proteins in human visual cortex (13-18). The data set created a *n*x*p* matrix comprised of *n*=403 rows of observations and *p*=23 columns of protein variables (Tables 4 & 5). The data were collected from the same tissue samples, across a series of different experiments, and each experimental data set was stacked vertically creating an initial matrix of 9,269 cells and an initial N= 1,831 data points. The empty cells created by stacking experimental data sets (7,438 empty cells) were reduced by converting protein expression across the 23 protein variables to a sample average for each of the 31 samples, creating a final matrix of 713 cells and a final N=651 data points.

**Table 4.** Observations (*n*)

| Categories | Specific | Total |
| --- | --- | --- |
| Age | 2 days - 79 years | 31 |
| WB Runs | 2-5 | 2-5 |

**Table 5.**
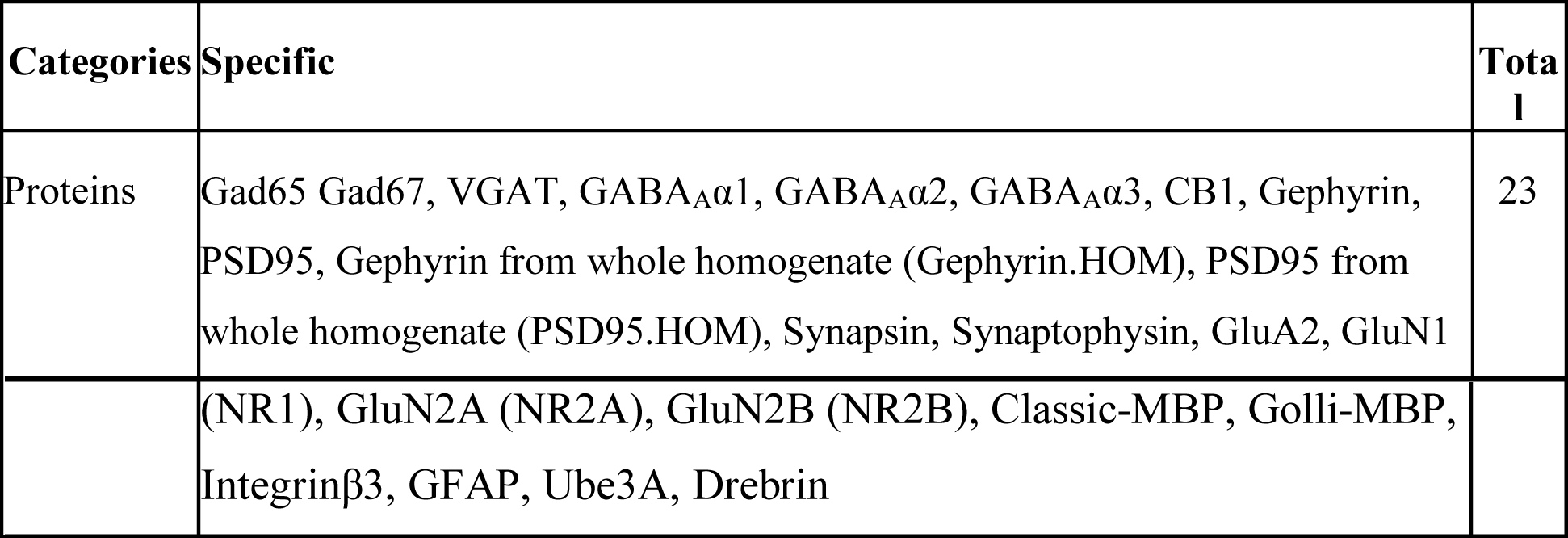
Variables (*p*)

| Categories | Specific | Total |
| --- | --- | --- |
| Proteins | Gad65 Gad67, VGAT, GABA<sub>A</sub>α1, GABA<sub>A</sub>α2, GABA<sub>A</sub>α3, CB1, Gephyrin, PSD95, Gephyrin from whole homogenate (Gephyrin.HOM), PSD95 from whole homogenate (PSD95.HOM), Synapsin, Synaptophysin, GluA2, GluN1 | 23 |
| | (NR1), GluN2A (NR2A), GluN2B (NR2B), Classic-MBP, Golli-MBP,<br>Integrinβ3, GFAP, Ube3A, Drebrin | |

**Figure 19.**
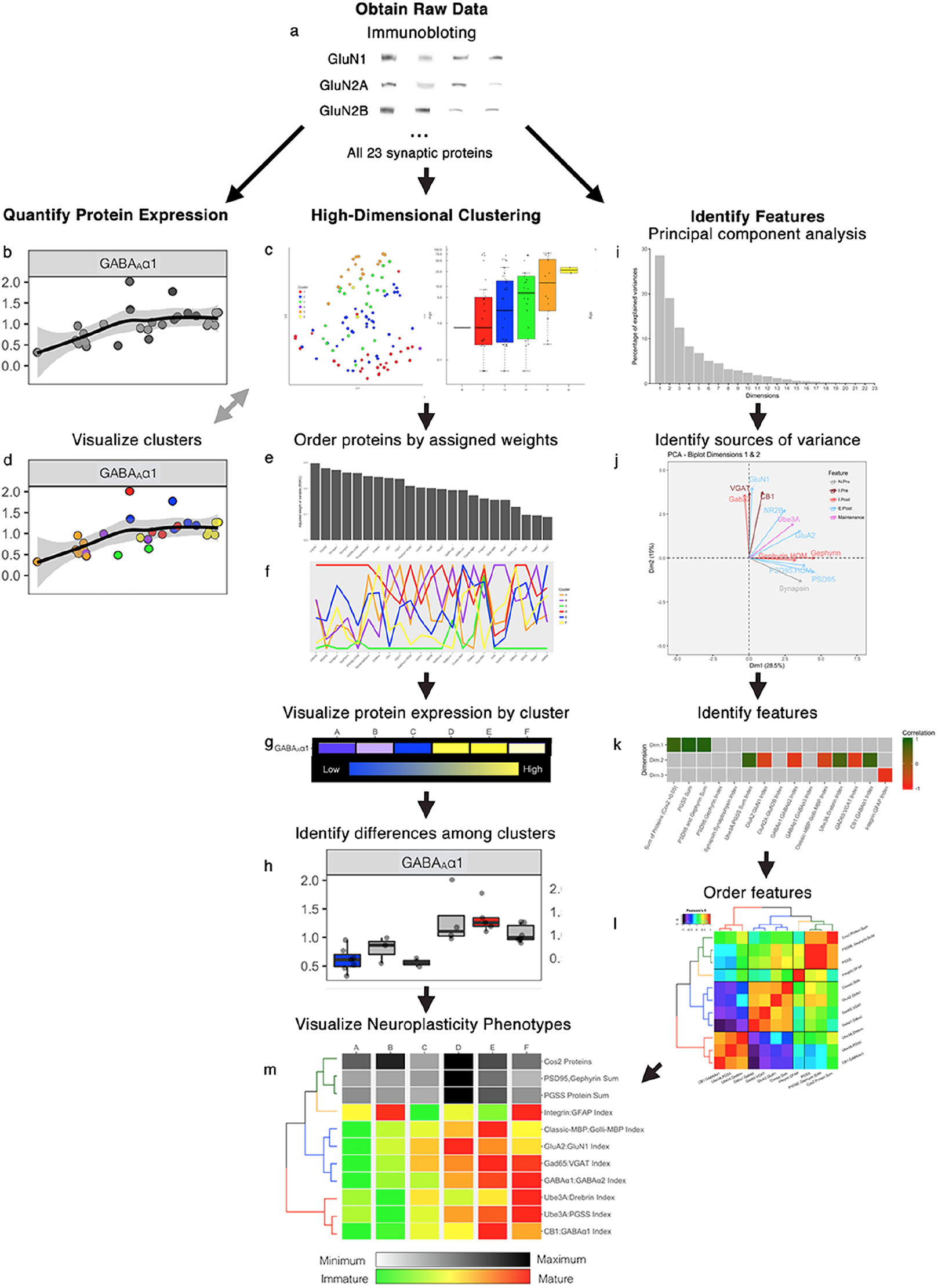
Protein analysis workflow for p>20 and p≅n. **a**. Protein expression was collected across 23 proteins using immunoblotting (N_funal_=651). **b**. Developmental trajectory of each protein visualized using scatterplots. **c.** We explored multiple high-dimensional clustering methods to identify underlying patterns across 23 proteins. A tSNE representation of the data was used to visualize the results of each clustering attempt (left), and boxplots of cluster median age were used to visualize developmental separation of clusters (right). **d.** Cluster information was fed into the developmental trajectories to color clusters in scatterplots. **e-f.** The weights attached to each protein after high dimensional clustering were used to order the protein variables and used to make a parallel coordinates plot of the average scaled protein expression for each cluster. **g.** Protein motifs of the same scaled data in f were created as a simple visual inspection tool to identify differences across clusters. **h.** Boxplots of clusters with significantly high or low expression relative to the full course of development. **i**. Principal component analysis (PCA) was performed to identify significant dimensions. **j.** Biplots of protein vectors visualize variables with the greatest variance. **k.** Identify candidate features and transform the data using the results of PCA and biplot analysis. **l-m**. Correlation matrix of new plasticity features and the surrounding dendrogram (**l**) were used to order the presentation of neuroplasticity phenotypes (**m**).

### 4.i) Challenges identifying clusters in a data set with many features

Clustering of high-dimensional data is challenging because of sparsity in the data set, and for that reason, it is common to use dimension reduction (e.g., PCA) as the first step to identify a smaller number of features that are correlated among the points and drive variance in the data. Then the subsequent clustering focuses on those features. That approach, however, breaks down when the features that differentiate clusters are not orthogonal (9) or, as in the current data set, represent subtle developmental shifts where some proteins are correlated with some features while others are correlated with other features. These are important considerations when studying cortical development because even a small change in protein expression can have a significant impact on neurobiological function and those changes may be missed by some approaches to high dimensional data analysis.

To overcome the problem that dimension reduction may prune off too much information, or miss more subtle changes in protein expression, we tested a set of sparse clustering algorithms using our human V1 development protein data set as the input. Those algorithms were designed to cluster high-dimensional data with strong sparsity, and here we show that a recent application (RSKC- (32)) based on the adaptive sparse clustering algorithm from Witten & Tibshirani (2010) performed best for clustering the developmental changes in protein expression. The Witten & Tibshirani algorithm was explicitly designed for sparse clustering of high-dimensional data when the structure of the data set is either *p* ≈ *n* or *p > n.* That is precisely the structure of the data set used here (*p*=23, *n*=31) furthermore that is generally the case for genomic or proteomic studies of human brain development where measurements are made for many genes or proteins (*p*) from a smaller number of cases (*n*).

It is important to note that sparsity in the context of high-dimensional cluster analysis does not mean empty cells in the data set but rather that for any pair of points in a high-dimensional space there are probably a few dimensions that separate them. Furthermore, those dimensions are probably not orthogonal hence the need to use a clustering algorithm that can adaptively search all dimensions of the data for both local and global features that characterize clusters in the data set.

### 4.ii) Sparse high-dimensional clustering to identify communities in the data

Here we describe and compare four sparse high-dimensional clustering methods for analyzing the development of human visual cortex. High-dimensional clustering is an active area of statistics research (e.g., (33)) with many new algorithms and approaches published yearly but the more commonly used ones are reviewed in Parsons et al., 2004.

The 4 clustering methods that we tested were selected because they were developed to find clusters in sparse high-dimensional data. The first two methods, CLIQUE (34) and PROCLUS (35), use projected clustering to discover dense regions, or subspaces of correlated points and find clusters in the corresponding subspace. The CLIQUE algorithm is a bottom-up approach moving from lower to higher dimensionality subspace, but it does not strictly partition points into unique clusters so a data point may be assigned to more than one cluster. CLIQUE is also prone to classifying points as outliers and excluding them from the analysis. PROCLUS was developed to address the partitioning problem and uses a 3-step top-down approach to projected clustering based on medoids. The steps involve initializing the number of clusters (*k*) and the subspace search size (number of dimensions to consider), then iteratively assigning medoids to find the best clusters for the local dimensions, and a final pass to refine the clusters. PROCLUS has better accuracy than CLIQUE in partitioning points into clusters but the *a priori* selection of cluster size is not easy and necessitates the iterative approach to finding clusters. Furthermore, by restricting the subspace search size, some essential features may be omitted from the analysis.

Both CLIQUE and PROCLUS were developed with large data sets, 2-3 orders of magnitude larger than the data set used here and that higher information content can lessen the partitioning, outlier, and feature omission problems. For our purposes, we also needed to test sparse clustering designed for smaller data sets where *p* ≈ *n* or *p > n*, but that also met the criteria introduced above:

1. use as many features as possible to identify clusters
2. group all of the samples into distinct clusters containing at least 3 samples
3. do not exclude points as outliers

Those criteria led us to select two more approaches to sparse hierarchical clustering, SPARCL and RSKC.

The sparse hierarchical clustering method SPARCL was developed by Witten & Tibshirani (2010), and clusters data points using an application of the lasso regression method to select local subsets of features adaptively. Those subsets are applied by scaling the weight of each variable, the proteins in our data set, to reflect the impact of each protein on the features and the reweighted proteins are the input to *K*-means hierarchical clustering. In our application, all of the reweighted proteins were part of the clustering but it is possible to drop variables from the clustering component of this approach. The adaptive feature selection of SPARCL focuses on the subset of proteins that underlie the differences among clusters and that process is similar to removing noise from the data. Finally, SPARCL makes it easier to draw meaningful conclusions about why data points are in a cluster because clustering is determined using the subset of features responsible for differences among the data points.

The Witten & Tibshirani sparse clustering algorithm has many strengths for analyzing data sets with *p* ≈ *n* or *p > n*; however, it can form clusters containing just one observation (36). A recent extension, Robust and Sparse *K*-means Clustering (RSKC), addresses the issue of small clusters by assuming that those are caused by outlier data points. RSKC uses the same clustering framework as SPARCL, except that it is *‘robust’* to outliers (32). RSKC iteratively identifies clusters in the data, then identifies clusters with a small number of data points (e.g., n=1) and flags these data points as potential outliers. The outliers are temporarily removed from the analysis, and clustering proceeds as outlined above for SPARCL. Once all clusters have been identified, the outliers are re-inserted in the high-dimensional space and grouped with the nearest neighbour cluster. Thus, RSKC identifies meaningful clusters in the data and includes all of the data points.

### 4.iii) Applications

Here we compare four high-dimensional clustering approaches (PROCLUS, CLIQUE, SPARCL, RSKC) using a subset of the proteins in the human visual cortex develop data set. Then describe the application of RSKC to the full set of proteins in the data set.

First, we tested the two density projection clustering methods that use either top-down (PROCLUS) or bottom-up (CLIQUE) clustering methods with all of the observations (*n*=31) and 7 of the proteins from the human visual cortex development data set. The outputs were plotted in 2D using tSNE, and the data points were colour-coded according to the clusters identified by each method. Finally, to determine if the clusters represented developmental changes in the data set we plotted boxplots showing the median ages for each cluster.

#### PROCLUS

The PROCLUS clustering methods was implemented in RStudio using the *ProClus* function in the *subspace* package (37). We explored clusters between *k*=2-9 and the example code below is for *k*=2 clusters.

The data file was read into an object called `my.data2` and the clustering function *ProClus* was called. tSNE was run on `my.data2` to visualize the data points in 2D and the points were color-coded using their cluster identification determined from *ProClus*. The results were saved in `my.proclus.tSNE` and that object was used to create the tSNE plots.

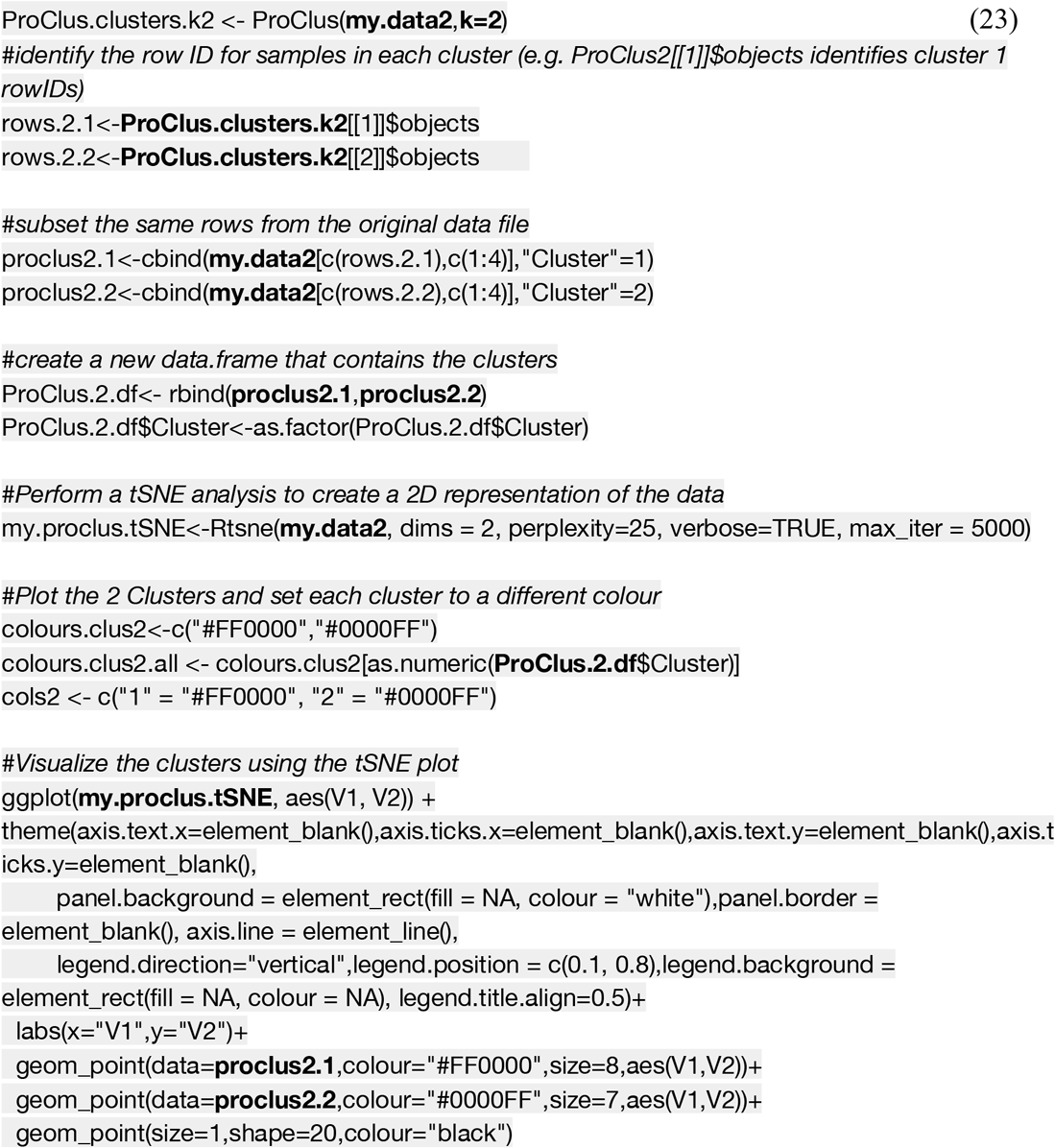

In this example, PROCLUS identified clusters but some observations (small grey dots) remained without a cluster designation and were interpreted as outliers (Figure 20A). The outliers remained even after stepping through a range of cluster numbers (*k*=2-8) and some of the clusters had only one or two data points (Figure 20Ai-iii). Furthermore, there was only a weak progression in the median age of the clusters (Figure 20Bi-iii). Thus, the iterative top-down feature identification and cluster border adjustments of PROCLUS performed poorly for analyzing the human visual cortex development data set.

**Figure 20.**
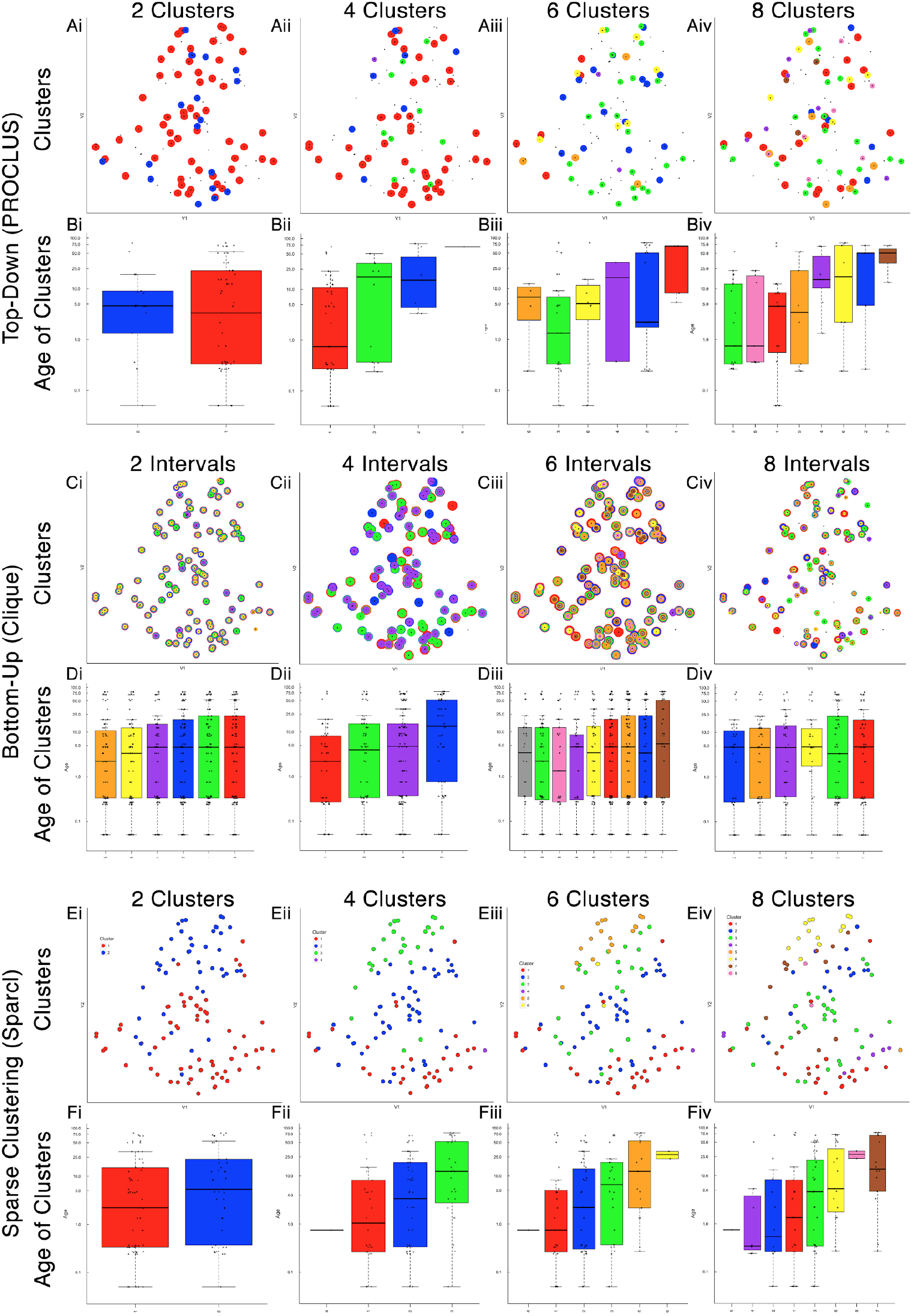
Top-down, bottom-up and sparse subspace clustering. **A-B.** Top-down PROCLUS subspace method across range of cluster numbers (2,4,6,8). The clusters are visualized in tSNE representations of the data by color-coding each data point with its cluster identity (Ai-iv) and in histograms showing the median age of the samples in each cluster (Bi-iv). **C-D.** Bottom-up Clique subspace clustering method for a range of ‘intervals’. Different clusters are visualized as coloured dots in a tSNE representation of the data (Ci-iv) and as histograms depicting the mean age of the samples (Di-iv). **E-F.** Sparse clustering after varying the inputted k cluster number (2,4,6,8). Different clusters are visualized as coloured dots in a tSNE representation of the data (Ei-iv) and as histograms depicting the mean age of the samples (Fi-iv). The colours in scatterplots and histograms represent the cluster designation for all plots.

#### CLIQUE

Next, the bottom-up clustering method CLIQUE was tested to determine how well this iterative approach to building clusters performed with the developmental data.

The *CLIQUE* function from the *subspace* package (37) was used to test clustering with the same data set tested with PROCLUS (`my.data2`). CLIQUE requires an input value for the interval setting (xi) because the intervals divide each dimension into equal width bins that are searched for dense regions of data points. Here we tested a range of intervals (xi=2-8) that produced 4-9 clusters.

The coding example below (24) used xi=2 as the input interval value and that produced 6 clusters that were visualized in a tSNE plot using `my.clique.tSNE` object (Figure 20 C&D).

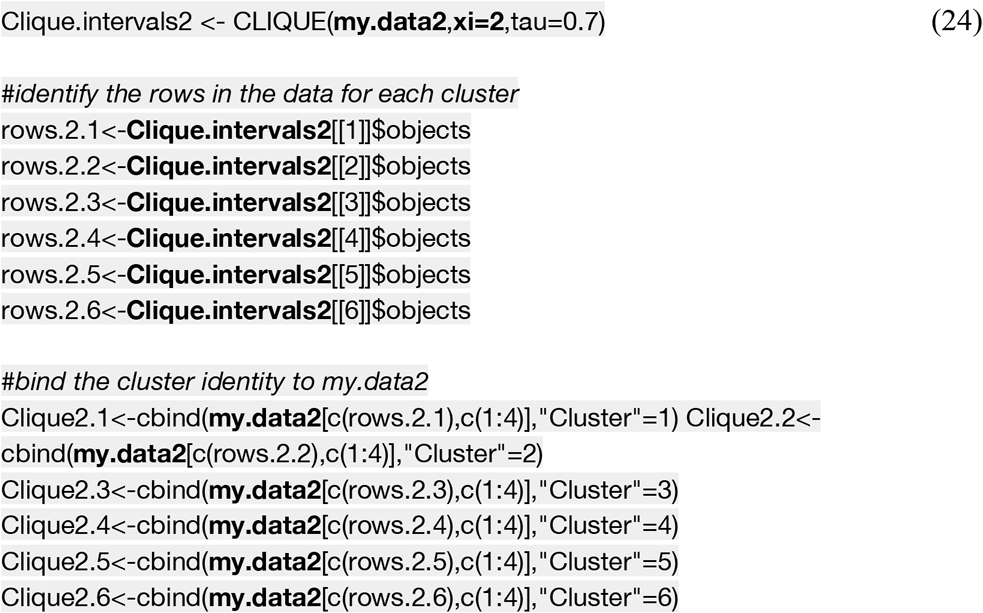

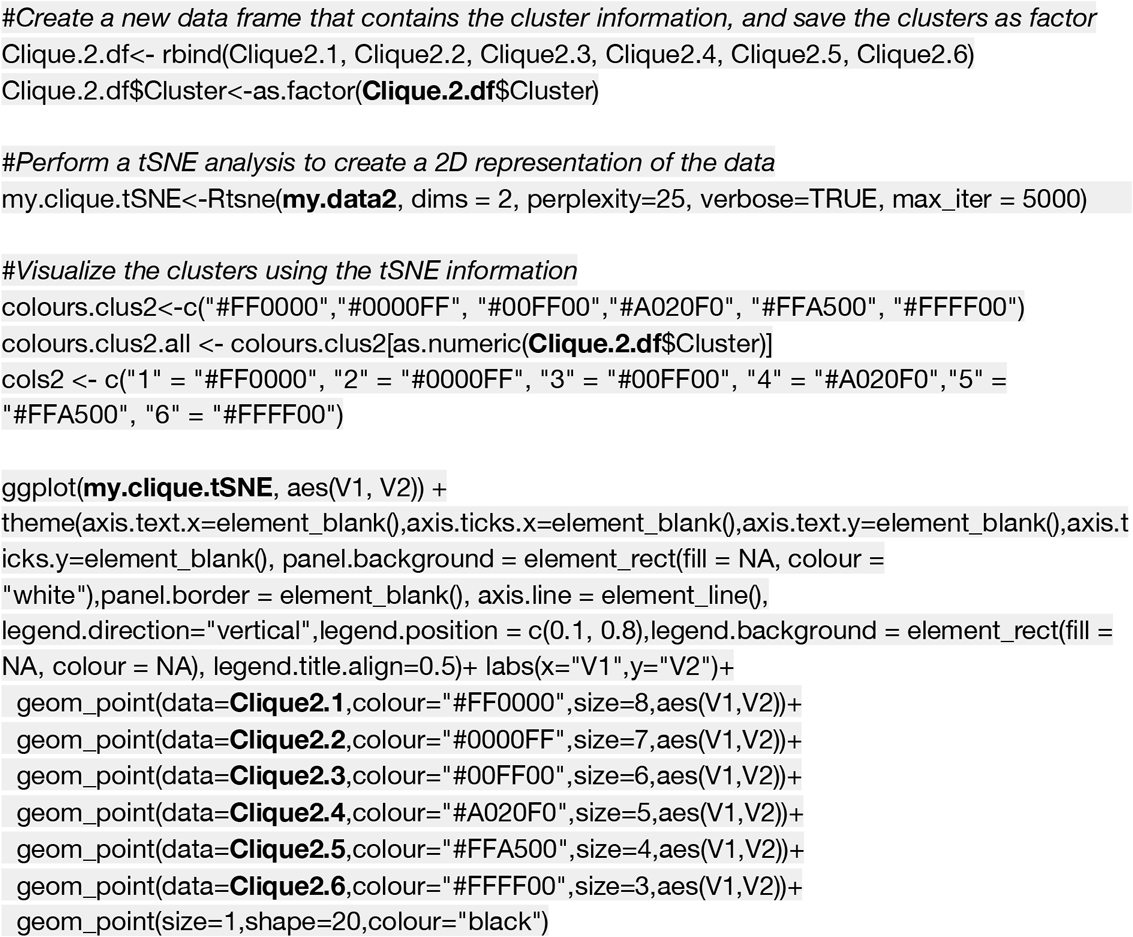

CLIQUE allows data points to be in more than one cluster and to visualize the multi-cluster identities we plotted the data points using concentric color-coded rings. CLIQUE placed all of the data in the developmental data set into multiple overlapping clusters for all of the interval settings (xi=2-8, Figure 20C). That poor partitioning of points resulted in no differences in the median cluster age (Figure 20D). Thus the iterative bottom-up clustering of CLIQUE performed poorly for clustering the developmental data set and did not reveal age-related clustering.

Comparing the top-down PROCLUS and bottom-up CLIQUE methods showed that neither approach was appropriate for analyzing the human visual cortex developmental data set. PROCLUS performed somewhat better because some of the parameters resulted in clusters with a progression in the median cluster age. That better clustering may have been because PROCLUS used a subset of the proteins for each iteration; however, the number of data points treated as outliers was unacceptably high.

We tested a third algorithm, sparse *K*-means clustering SPACRL (36) that adaptively finds subsets of variables capturing the different dimensions and includes all samples in the clusters. SPARCL searches across multiple dimensions in the data and adjusts the weight of each variable based on their contribution to the clustering. The term ‘sparse’ here refers to the selection of different subsets of proteins to define each cluster, but all samples are assigned to a cluster.

#### SPARCL

To implement sparse *K*-means clustering, we used the *Kmeans.sparsecluster* function in the *sparcl* package (38). We explored a range of *k* clusters between *k*=2-9. The *sparcl* package also includes a function to help determine other input variables such as the boundaries for reweighting the variables (*wbounds*) to produce optimal clustering.

This coding example (25) identifies the optimal *wbounds* setting for *k=*2 clusters:

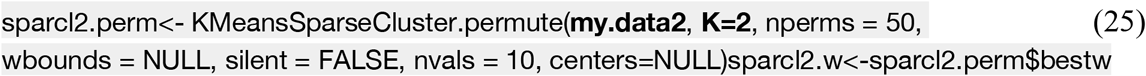

The coding example below calculates sparse clustering for *k*=2 clusters with the *wbounds* found above:

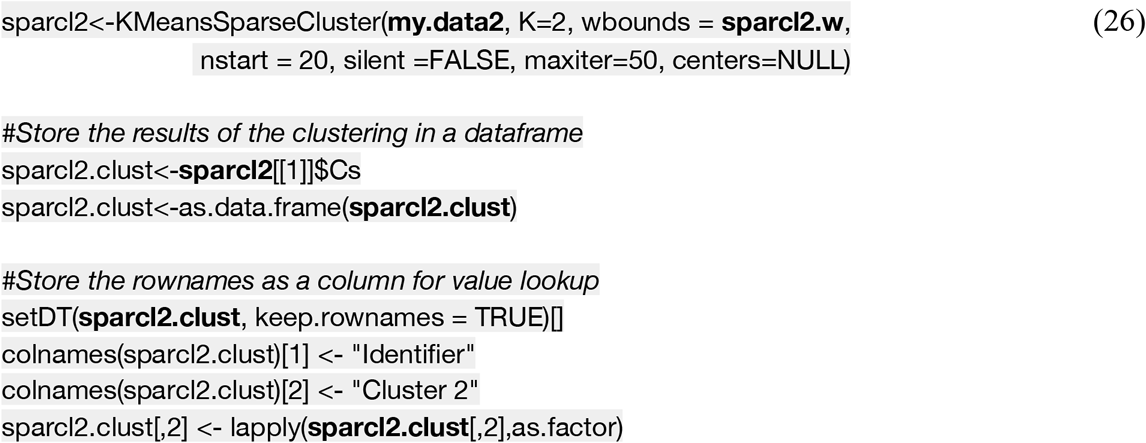

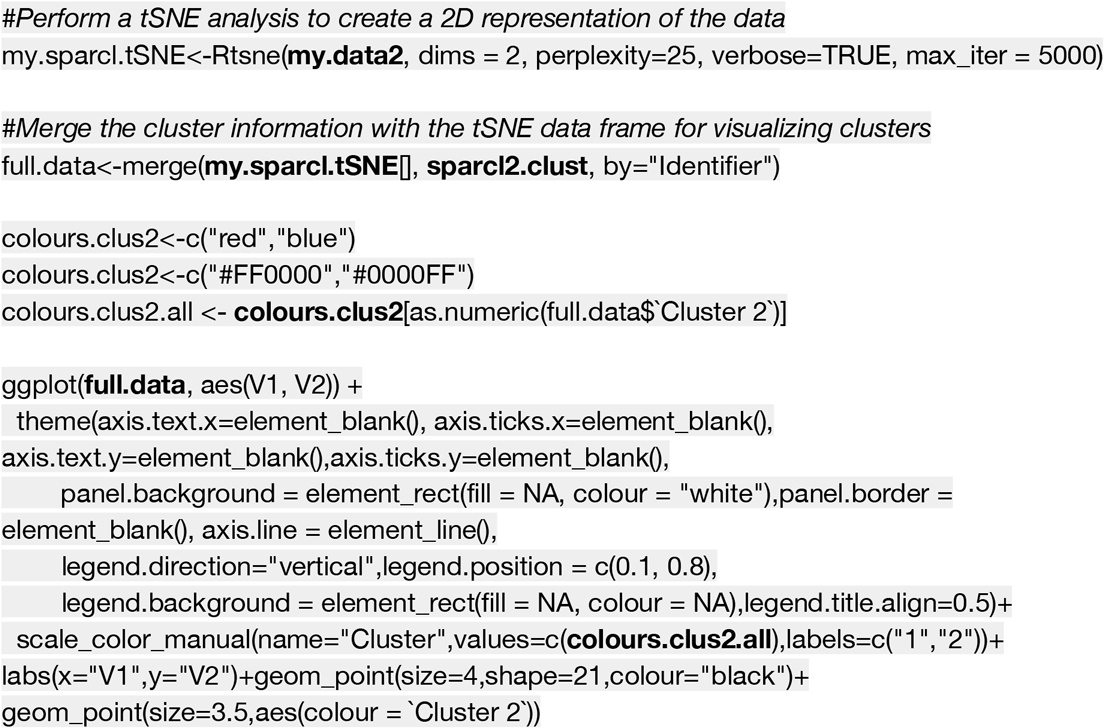

#### RSKC

The clusters found with SPARCL showed good partitioning of the data, and with 4-8 clusters, there was a progression of the median cluster age (Figure 20 E, F). However, some clusters had a small number of data points (e.g. n=1). To address that problem we turned to a modified version of the SPARCL algorithm called Robust and Sparse K-means clustering (RSKC) (32). The RSKC algorithm was designed to be robust to the influence of outliers that can drive other algorithms to create clusters of n=1. RSKC operates by iteratively omitting outliers from cluster analysis, assigning all remaining samples to clusters, and then reinserting outliers to the analysis by grouping them into the nearest-neighbouring cluster.

We used information from the SPARCL analysis to set the the number of clusters to *k*=6 for testing the RSKC method. The *RSKC* function in the *RSKC* package (32) was implemented for this analysis and the code below was used to analyze `my.data2`.

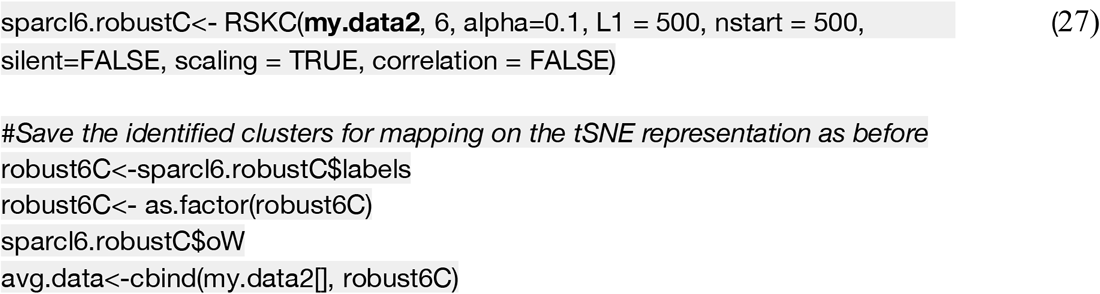

The initial test of RSKC on the subset of data gave the best clustering so RSKC was rerun using the full set of 23 proteins. The output from the RSKC method showed a clear progression in the median age of the 6 clusters from the youngest samples in cluster A to the oldest samples in cluster F (Figure 21). Furthermore, the boxplot showed minimal overlap of the cluster ages.

**Figure 21.**
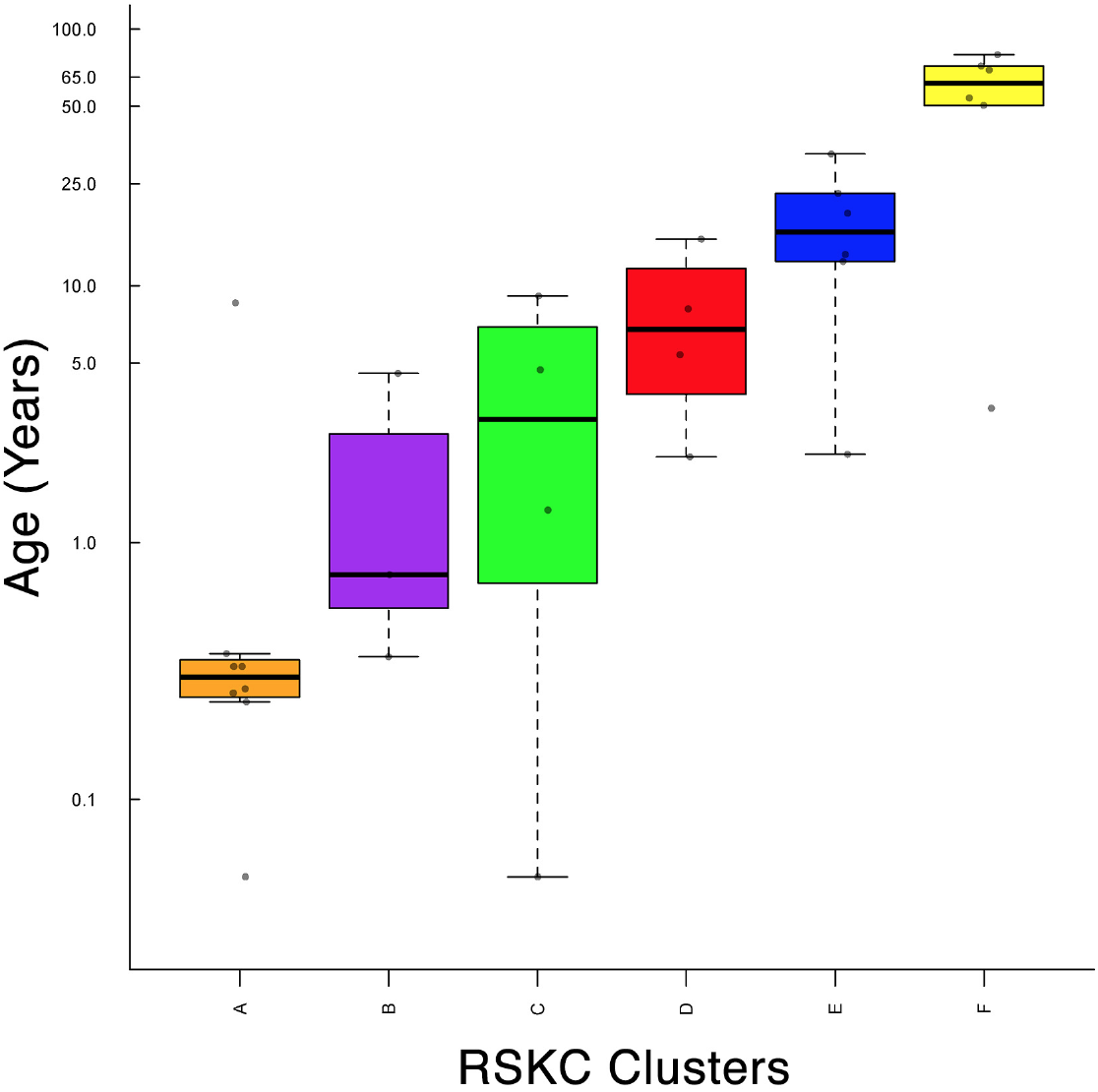
Robust and sparse k-means clustering (RSKC) applied to 23 proteins from the human data set. Sample ages were reduced to group averages to reduce crowding. Boxplots were ordered from youngest (orange) to oldest (yellow).

RSKC used all of the data, partitioned all of the samples into a cluster, and none of the clusters had fewer than 3 samples (Figure 21). Thus, RSKC clustering met the criteria we set for selecting a good approach to analyze the visual cortex development data set. Importantly, we analyzed the age progression of the clusters by comparing the observed pattern with a Monte Carlo simulation that randomly assigned the samples randomly into one of the 6 clusters. The simulation showed that it was highly unlikely for the progression of clusters ages shown in Figure 21 to occur by chance.

In section 4.iv we explain how to explore the RSKC clusters to identify neurobiological features that changed across development.

It is important to note that the WSS metric used in the first workflow to assess the number and quality of the clusters cannot be used with the sparse clustering methods. The lack of a cluster quality metric is because all the data are used with the 4 sparse high-dimensional clustering algorithms, so the number and quality of clusters vary depending on the number of dimensions that are considered. As a result, there is no suitable metric to assess the quality of clusters in high-dimensions and the different sparse clustering methods must be compared by adjusting the input variables iteratively (Proclus=k, CLIQUE=xi, SPARCL=k, RSKC=k).

### 4.iv) Exploring cluster content

Cluster content was explored with two methods. First, information from the RSKC clustering was used to color-code the samples by their cluster ID and the developmental trajectory was plotted for each protein (Figure 19b). Next, the proteins were ordered by their assigned weights from the RSKC clustering (Figure 22), and the normalized expression of each protein was plotted for the six clusters (Figure 23 & 24).

**Figure 22.**
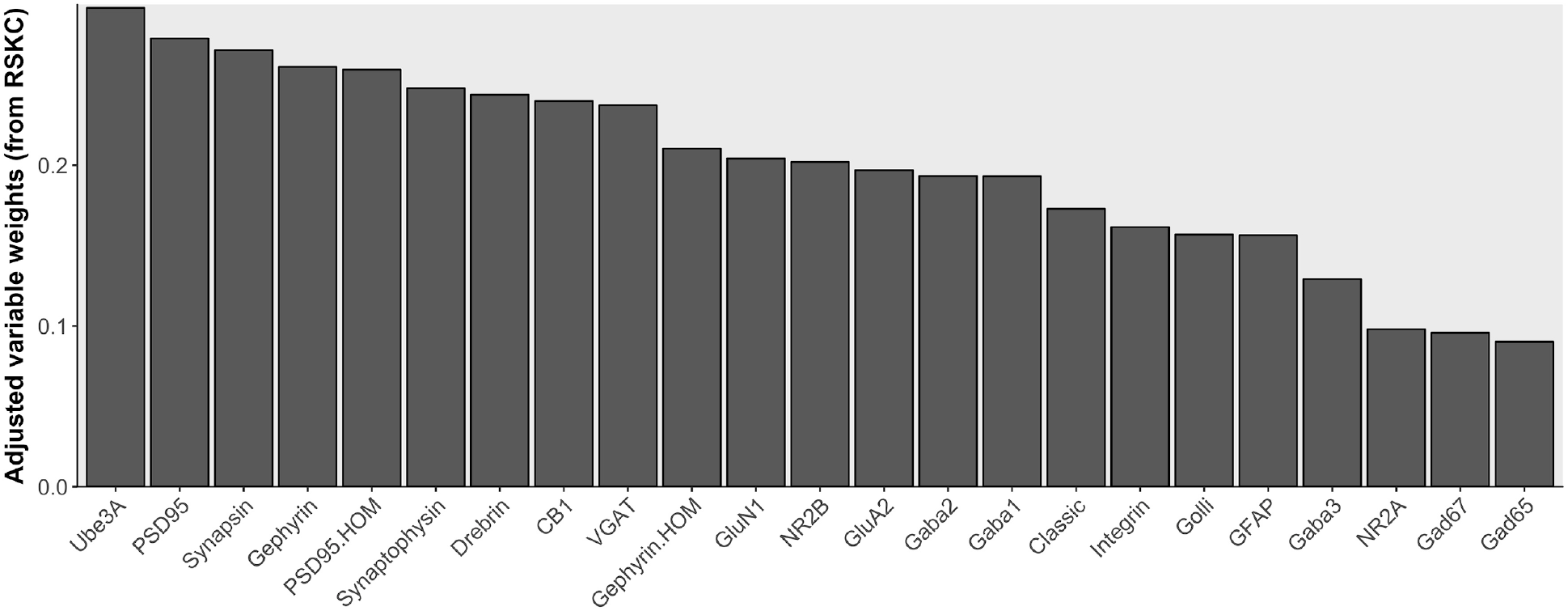
Adjusted weights for the proteins from RSKC. The proteins are ordered using the adjusted weights from RSKC.

**Figure 23.**
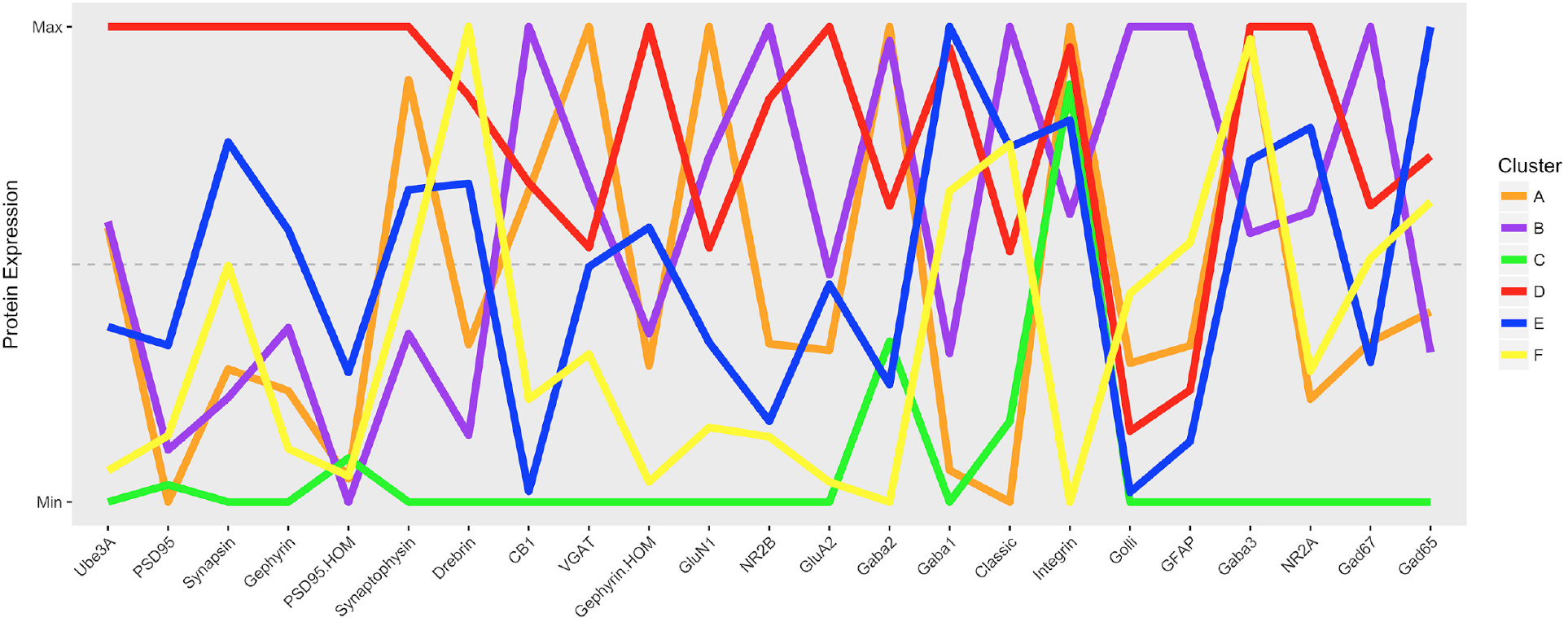
Parallel coordinates of the protein expression in each cluster. The expression is scaled to the average maximum and minimum expression across all 6 clusters. The colour of the lines is the same conventions for cluster identity as in Figure 21. Proteins were ordered from largest to smallest RSKC weights.

The second method to study cluster content (Figure 19 i-k) started with identifying features using PCA as described in section 3.iv. Then hierarchical clustering was used to order the features and finally, plasticity phenotypes were constructed for each cluster to visualize the neuroplasticity features that change during development of human visual cortex.

Together these approaches to exploring the cluster content provide an understanding of cortical development at the level of individual proteins and higher-level combinations of proteins that define features in the plasticity phenotype.

The following code consults the RSKC object sparcl6.robustC to retrieve the weight for each protein and then graphs the proteins in descending order of the weights.

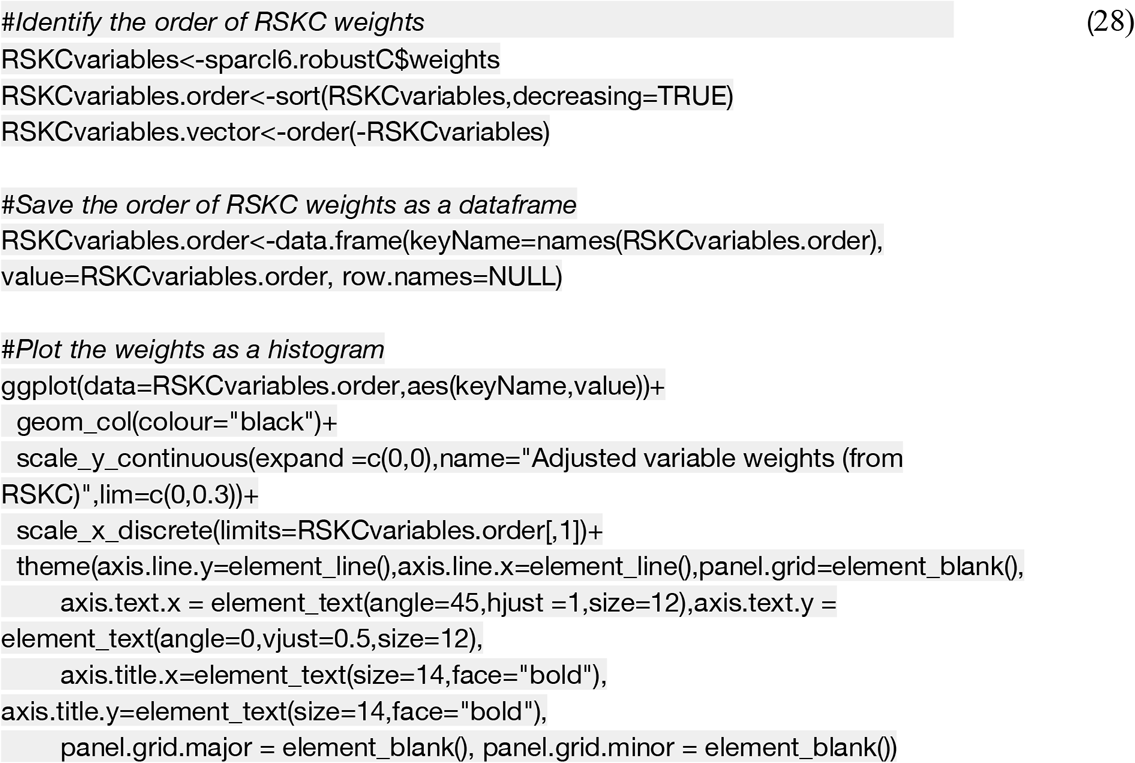

Visualizing the RSKC adjusted weights for the 23 proteins showed an almost 3-fold range in the weighting of the proteins with Ube3A having the maximum and GAD65 the minimum weight (Figure 22). That ordering of the proteins was used along with 2 visualizations of protein expression to explore the influence of different proteins in the clusters. First, we created a parallel coordinate plot for the 6 clusters and 23 proteins that scaled mean protein expression relative to the maximum and minimum cluster average using the *ggparcoord* function in *ggplot2* package (31).

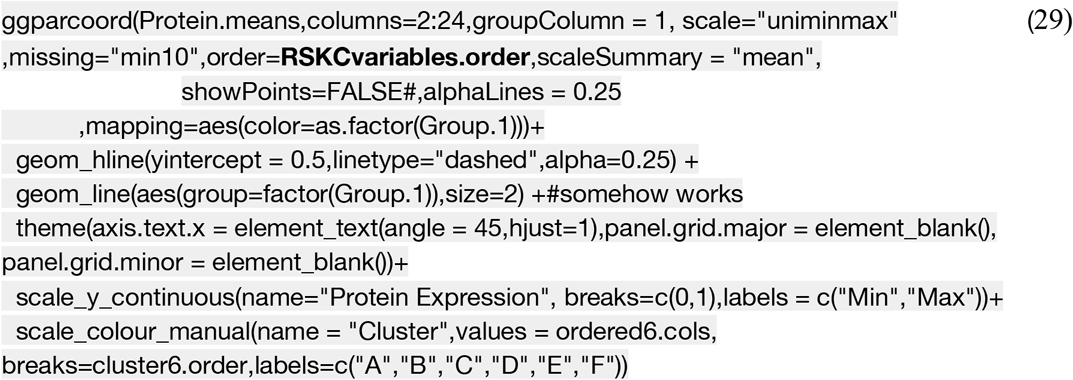

The parallel coordinates plot in Figure 23 showed that the 6 clusters had unique patterns of peaks and valleys in protein expression, and most of the protein peaks were claimed by only one cluster. For example, Cluster D had peak expression for the first 6 proteins (Ube3A, PSD95, Synapsin, Gephyrin, PSD95HOM and Synaptophysin), then cluster F peaked for Drebrin, B for CB1, A for VGAT and so on. Interestingly, the 4 proteins that had peaks from multiple clusters peaks included the 3 GABA_A_R subunits and Integrinβ3.

The information from the parallel coordinates graph can be used to construct a list of protein peaks and valleys for each cluster where those patterns reflect the protein motif for each cluster. Here we visualized the protein motifs for the 6 clusters using the *geom_tile* function in the *gplots* package. The coding for the motif visualization was similar to the method outlined in coding examples (21) and (22) except here it was constructed using the average protein expression.

Exploring the cluster content with the protein motifs helped to differentiate the groups of clusters that shared protein peaks. For example, the two youngest clusters A and B shared the GABA_A_α2 peak, but the older clusters D, E, and F shared the GABA_A_α1 peak. The motifs visualization facilitated the comparison of protein expression within are across clusters since all of the 138 protein averages (6 clusters X 23 proteins) were represented in one figure.

Next, we quantified protein expression in each cluster, plotted those in boxplots and used a bootstrap analysis to determine clusters that had expression significantly above (red boxes) or below (blue boxes) the median protein expression level. The boxplots for each protein were generated using the geom_boxplot function in the *ggplot2* package (31).

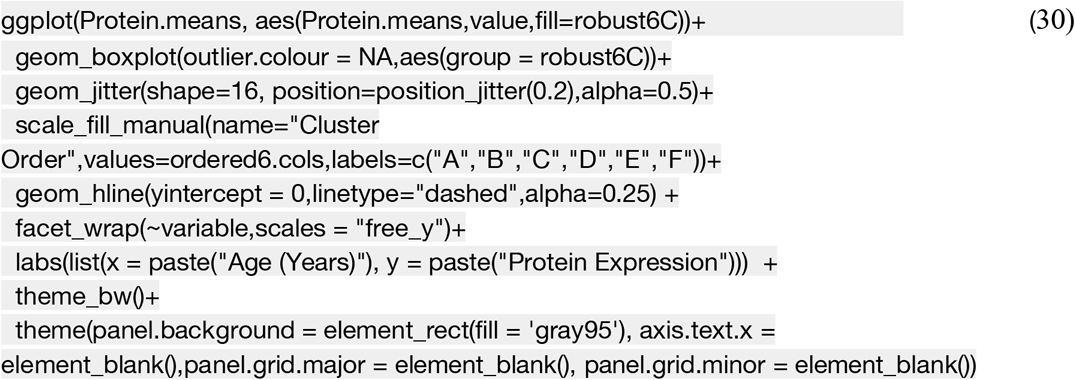

Using the protein motifs to visualize protein expression by cluster (Figure 24) and the color-coded boxplots to quantify protein expression (Figure 25) helped with efficiently exploring significant patterns in the clusters. That approach, however, did not readily identify combinations of proteins that represent higher-dimensional features in the data set. To explore if high-dimensional features identify age-related changes in the clusters we returned to the use of PCA.

**Figure 24.**
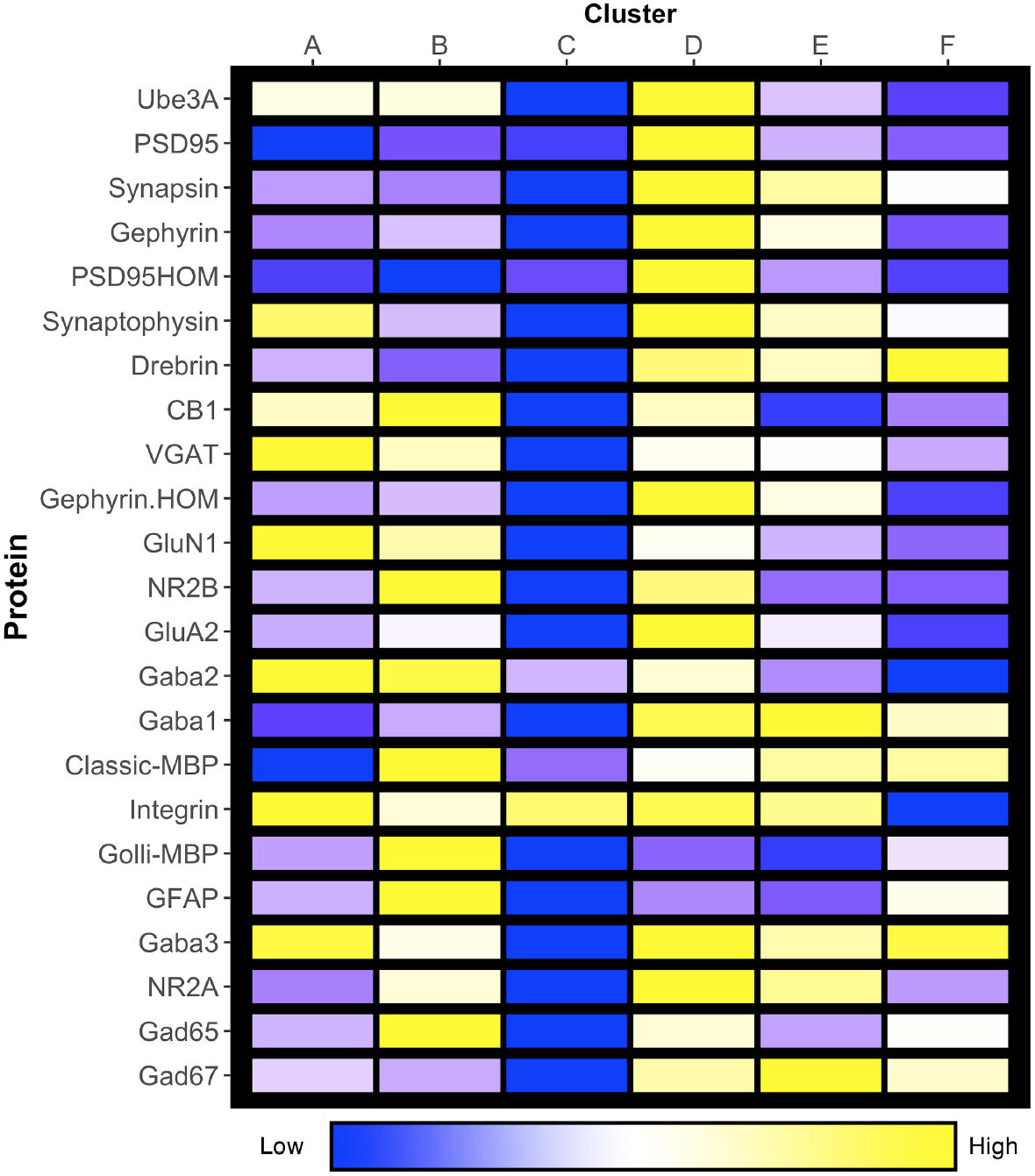
Protein expression motifs for the 6 clusters. The average protein expression across clusters was scaled from the minimum (blue) to the maximum (yellow). The proteins were ordered according to their RSKC adjusted weights.

**Figure 25.**
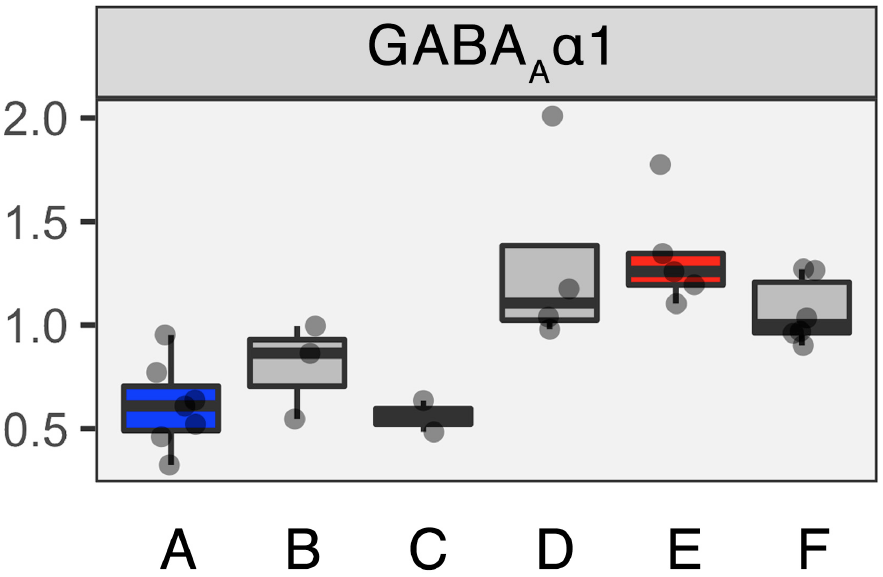
Protein expression per cluster. Each boxplot represents a different cluster of data points, and the dashed line overlaps balance (0) between each protein. Blue boxes represent clusters with median expression below the lowest 25% of the data, red boxes represent clusters with median expression above the upper 75% of the data, and grey represent those with median expression within the middle 50% of the data.

### 4.v) Candidate high-dimensional feature selection

To identify candidate high-dimensional features in the clusters we applied PCA to the human visual cortex developmental protein data set. The PCA found the significant dimensions and we used those to explore the proteins contributing to those dimensions. Also, we compared the proteins correlated with PCA dimensions to their adjusted weight from RSKC to uncover the hidden features that supported better clustering with RSKC.

The first step repeats the PCA described for the coding examples (8)-(11) in section 3.iii. We used the *PCA* function from the *Factominer* package to analyze and display the percentage of variance explained by each of the 23 dimensions (Figure 26). For this data set, the first 3 dimensions captured >60% of the variance which is often used as a cutoff for identifying the important dimensions.

**Figure 26.**
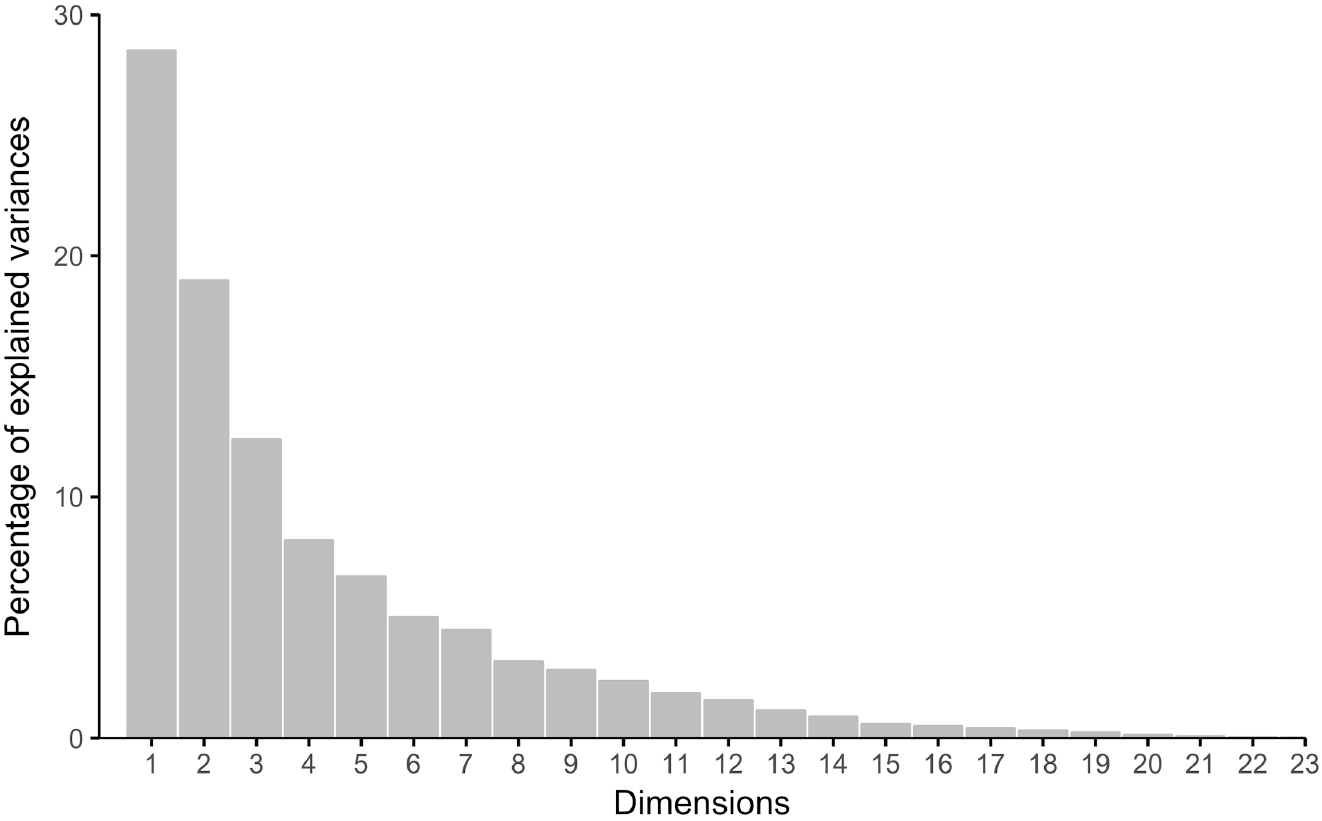
Scree plot of the percentage of explained variance captured by each principal component. The variance captured by the each of the first 3 dimensions is large (>10%), while subsequent dimensions gradually becomes less. The cumulative variance of the first 3 dimensions is 62%. Typically, a cumulative variance > 60% is an acceptable cutoff for identifying the important dimensions in PCA.

To interpret the representation of each protein on the PC dimensions we calculated pairwise correlations between the vector for each protein and the vector for each of the PC dimensions (Pearson’s correlation)(Figure 27A). Next, we determined the quality of representation for each protein on the dimension (cos^2^)(Figure 27B). The R code to calculate and plot those metrics is found in the FactoMineR package, and can be called by consulting the pca object:

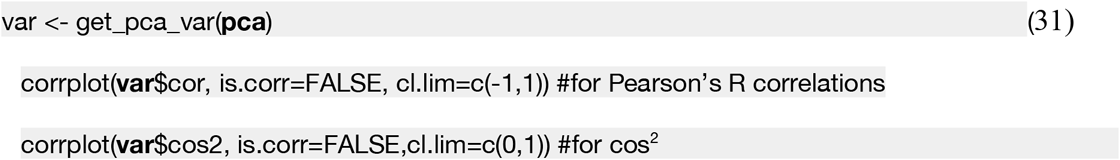

**Figure 27.**
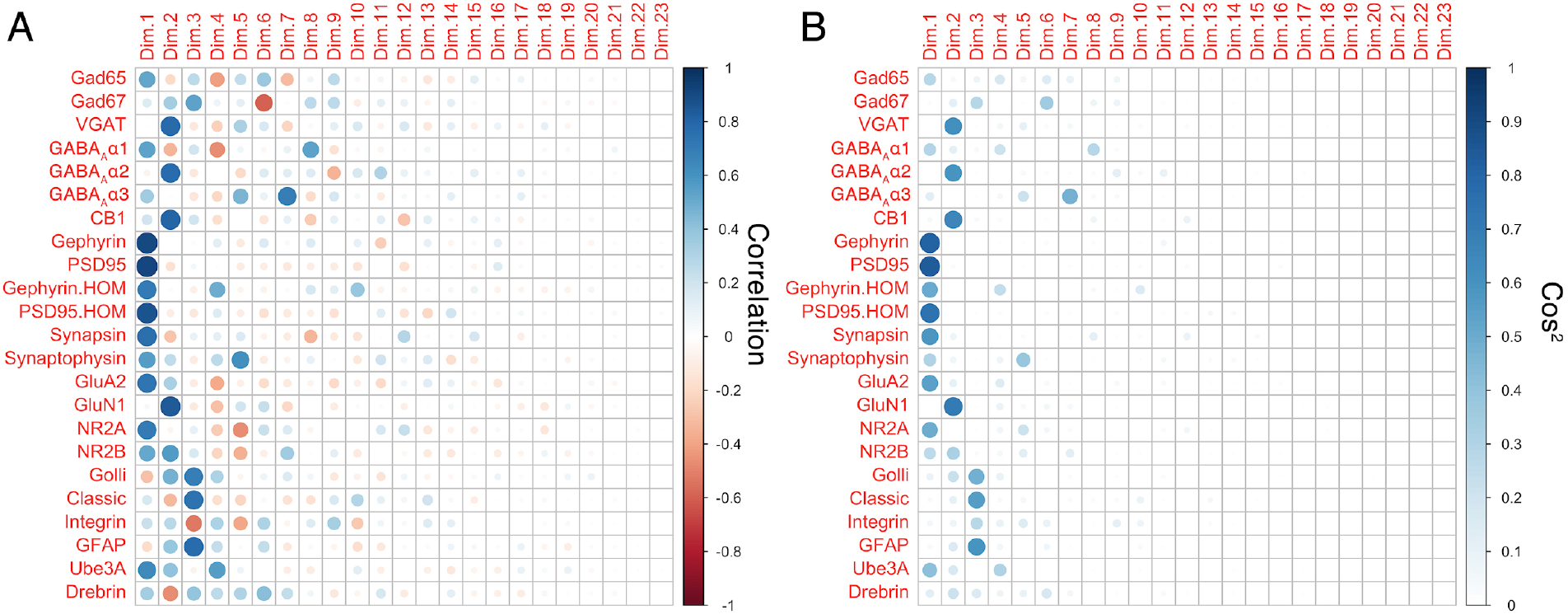
The correlation (A) and cos^2^ (B) matrix showing the relationship between each protein and the 23 principal component dimensions. **(A)** The circles in each matrix represent the strength (size and saturation) and direction (colour) of the correlation between proteins and dimensions. (B) The circles represent the goodness-of-fit, cos^2^ for the protein on PC dimensions. Most of the proteins have the strongest correlation and best fit with at least one of the first 3 dimensions (Dim.1, Dim.2, Dim.3).

The figures showing the correlations and cos^2^ for each protein with the 23 dimensions used the size and saturation of a circle to represent the strength of the relationship, and the color of the circle indicated the direction for the correlations (Figure 27A, 27B). Inspection of the plots showed that most of the proteins contributed to the first 3 dimensions but also identified a few proteins that had their most substantial contribution to a higher order dimension. This example helps to illustrate where PCA might lead to dropping variables from subsequent cluster analysis. It is important to note that the workflow described in this section ran PCA and sparse high-dimensional clustering in parallel with each analysis using the full protein data set so that the results from the two approaches could be compared to unpack the higher order features in the data set.

In this example, the first 3 dimensions were chosen because they captured >60% of the variance in the data (Figure 26), and most of the strong cos^2^ values and Pearson’s R correlations (Figure 27). The quality of the representation for each protein on the first 3 dimensions was analyzed by summing up the cos^2^ values and plotting that information as a histogram ranking the protein from lowest to highest cos^2^ value (Figure 28). The following code was used to visualize the sum of cos^2^ for the first 3 dimensions:

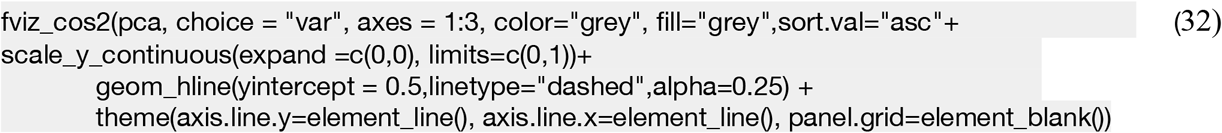

**Figure 28.**
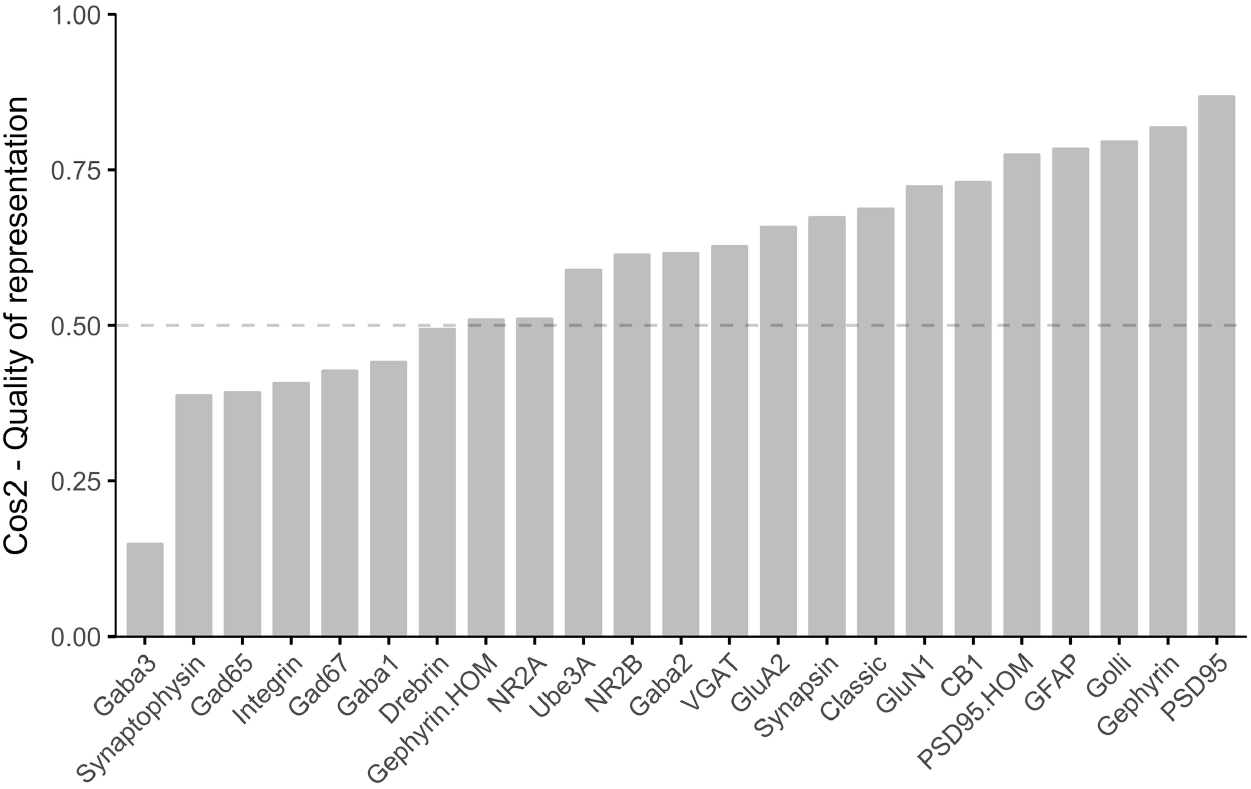
Sum of cos^2^ across the first 3 dimensions. Across all 23 dimensions, sum of cos^2^ for each protein is 1. Dashed line represents cos^2^ =0.5 cutoff, used for feature selection. 16 proteins fall above that cutoff and 7 fall below. The 7 that fall below can be removed form analysis (feature selection).

The cos^2^ can be plotted for individual dimensions. The example code below (33) uses the parameter `axes` to select the cos^2^ data for dimension 1.

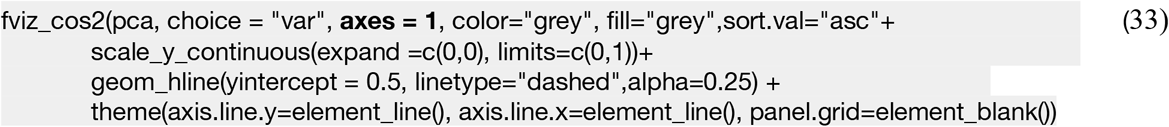

The cos^2^ value (e.g. 0.5) can be used as a cutoff to select a subset of proteins for further analysis but here were used the cos^2^ ranking (high to low) from the first 3 PCs (Figure 29) to compare with the ranking of each protein from the RSKC reweighting (Figure 22). This step was used to identify proteins that RSKC adaptively increased or decreased the weighting (Figure 28) and showed that 5 proteins were increased (Synapsin, Gephyrin.hom, Drebrin, Ube3A, Synaptophysin) while 4 proteins were decreased by RSKC (Golli-MBP, GFAP, Classic-MBP, GluN2A) (Figure 30). Those differential rankings provided some insight into the proteins that were important for the RSKC clustering but less influential for the PCA. It also illustrated the need for using multiple methods when studying high-dimensional cluster content.

**Figure 29.**
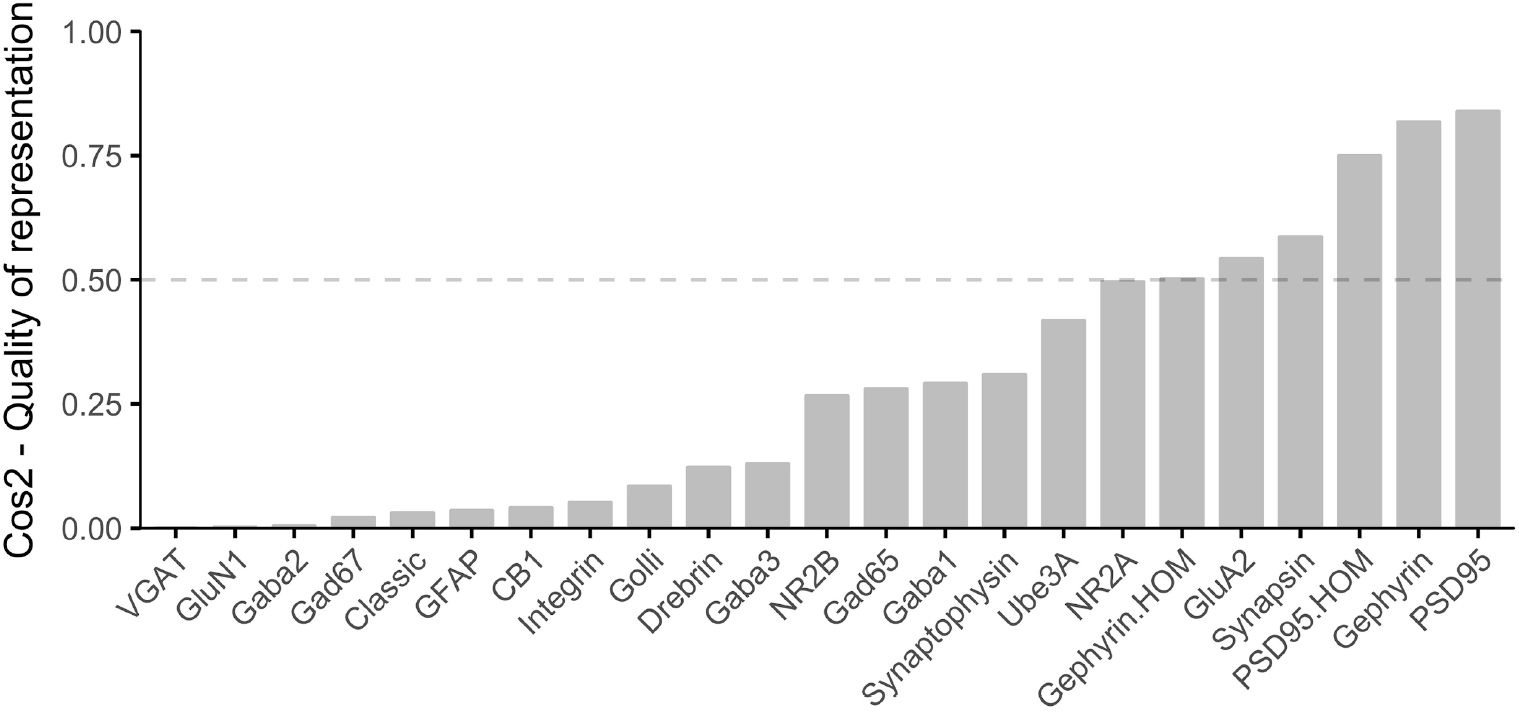
Sum of cos^2^ across dimension 1. The sum of cos^2^ for each protein in Dimension 1. Dashed line represents cos^2^ =0.5 cutoff, used for feature selection.

**Figure 30.**
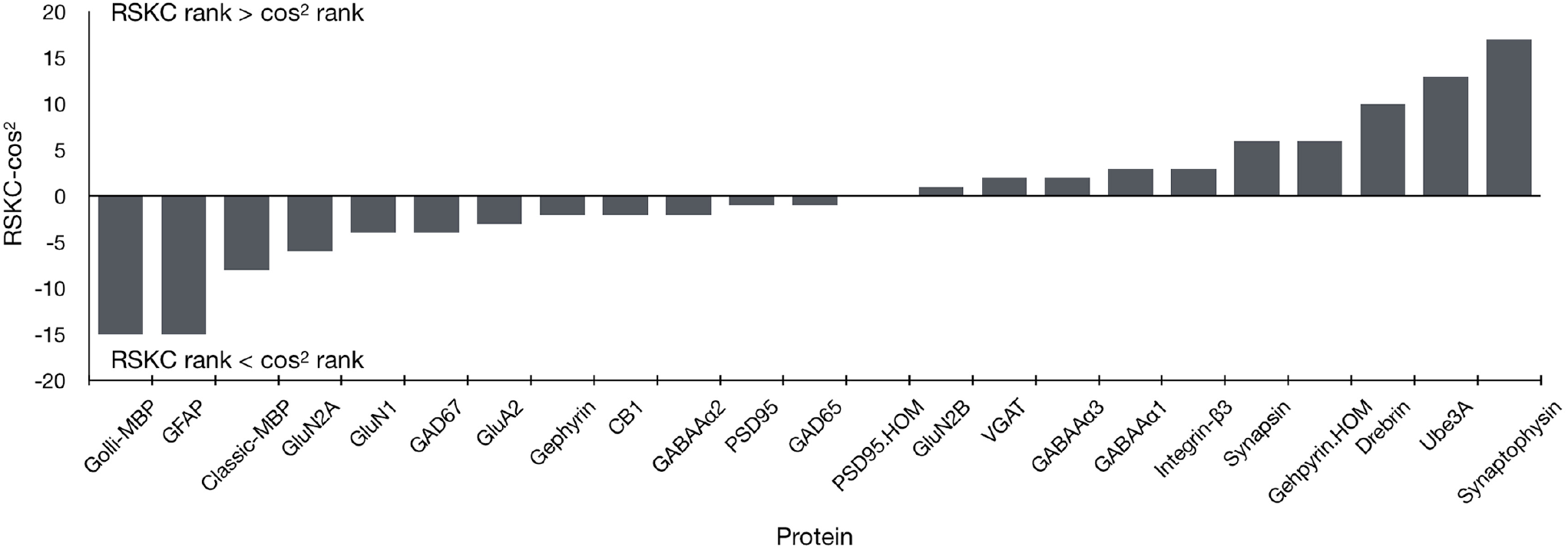
Ranking of proteins using RSKC reweighted values relative to cos^2^ representation. Positive values on the y-axis indicate a higher ranking using the RSKC reweighted values than with cos^2^ representation, and therefore have greater influence on clusters. Conversely, negative values indicate proteins that were ranked higher by cos^2^ representation than after RSKC reweighting.

Biplots were also used as another approach to visualize the contribution of different proteins to the PC dimensions. The example biplot (Figure 32) showed the vectors for select proteins (cos^2^>0.5) along dimensions 1 and 2. We used the *fviz_pca_biplot* function from the *FactoMineR* package to superimpose the protein vectors in PCA space.

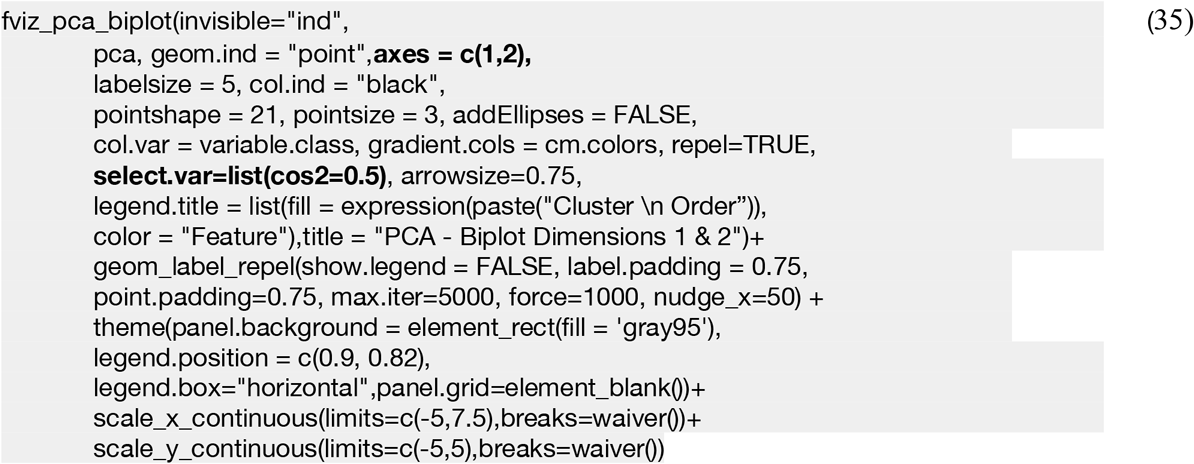

The pair of PC dimensions examined in a biplot can be changed using the values in ‘axes = c(1,2)’ parameter.

The length and angle of the vectors in Figure 31 was informative for identifying the proteins that contribute to each of the PC dimensions. Groups of proteins with similar vectors were correlated while orthogonal vectors indicate little correlation between those proteins. That information was used to guide the selection of proteins for candidate feature.

**Figure 31.**
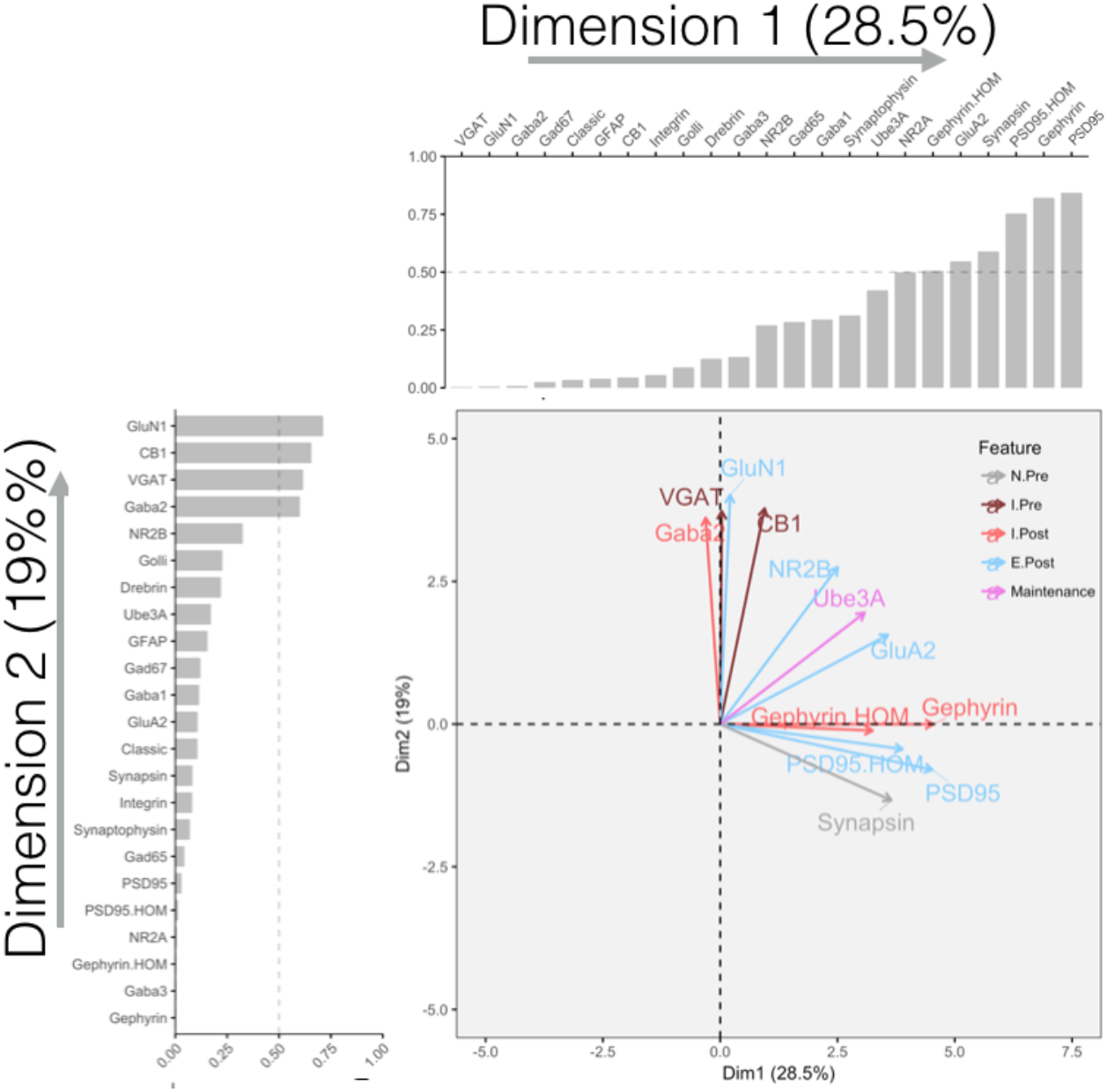
Biplot shows the protein vectors along dimension 1 (horizontal axis) and dimension 2 (vertical axis). Surrounding histograms depict cos^2^ values for the proteins along dimension 1 (top) and dimension 2 (left). Proteins with a summed cos^2^ above 0.5 (dashed line) across both dimensions are shown. Vector directions marked by arrowheads point in the direction of high protein expression, while low protein expression extends in the opposite direction through the point of origin (not shown). The length of the protein vectors indicates the variance of the protein along the dimension and the angle of the vector indicates the quality of the representation on a dimension. Parallel vectors indicate proteins with similar expression while perpendicular vectors have no relation.

We also plotted the individual samples onto the biplot by removing the line of code ‘invisible=“ind”’. The samples were color-coded by their RSKC cluster ID to visualize those clusters in PCA space (Figure 32). That biplot helped identify the relationship between the 6 clusters PC dimensions 1 and 2. For example, clusters C and D differed along dimension 1 while clusters A and F differed along dimension 2.

**Figure 32.**
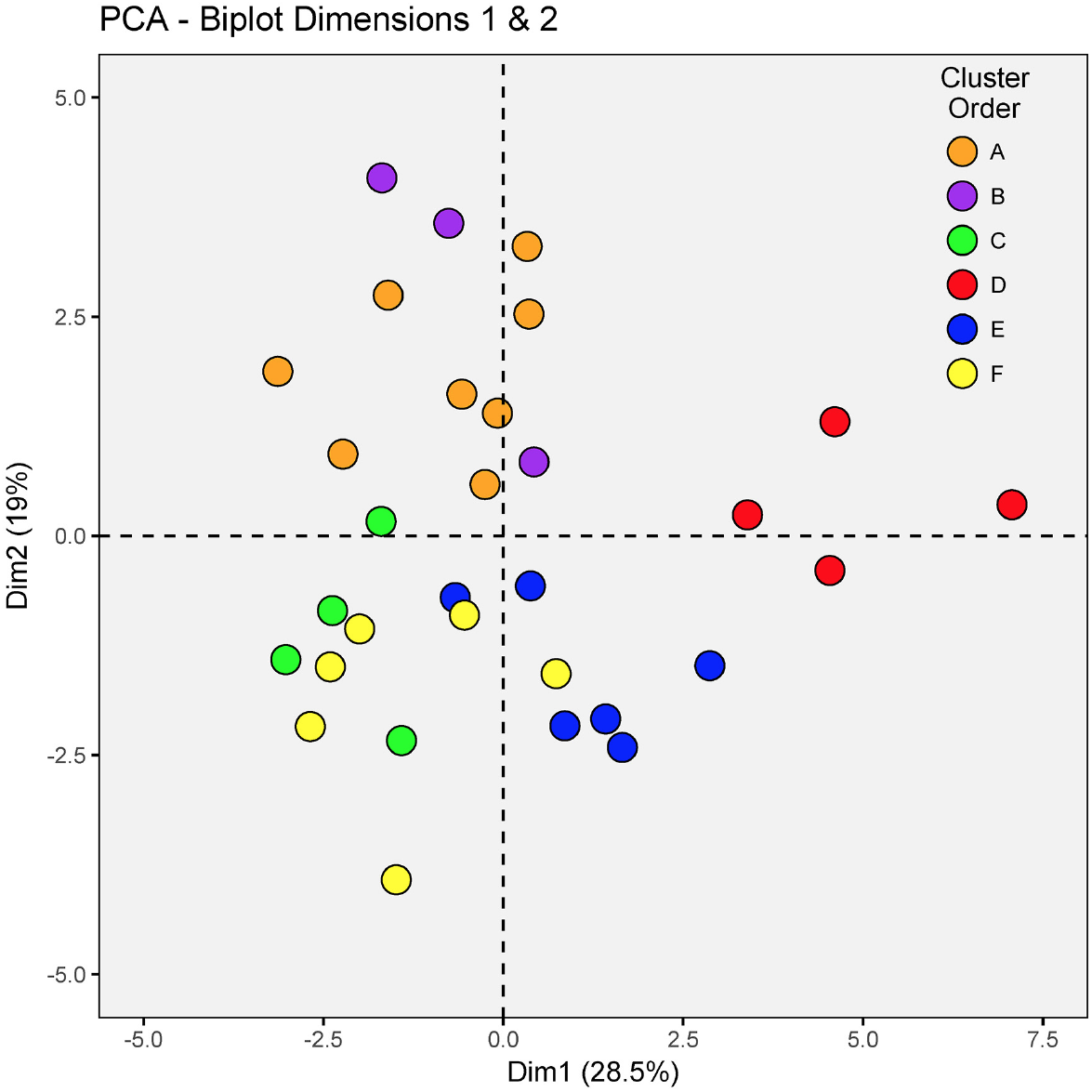
Biplots demonstrate the relationship between proteins and samples. Biplot of dimension 1 and 2 that shows the individual samples (dots) colour-coded according to the six clusters (A-F) they are grouped in. The axes conventions are the same as in figure 31.

In the next section we use the information gleaned from the cos^2^, differential RSKC-cos^2^ rankings and biplots about proteins that contribute to high-dimensional features in the data set to help select plasticity features.

### 4.vi) Converting protein expression into candidate plasticity features

Here we describe how the basis vectors from PCA and information from the previous section were used to select combinations of proteins for candidate plasticity features. The steps for using the basis vectors to identify candidate features follow those presented in section 3.iv but we also included information from the differential cos^2^-RSKC rankings to ensure that proteins reweighted by RSKC were appropriately considered for candidate features. Once candidate features were identified, they were validated by correlation with the first 3 PC dimensions.

First, the basis vectors were plotted to show the amplitude of each protein about the PC dimension (example for PC2, Figure 33). The code below plots the basis vector for the second PC:

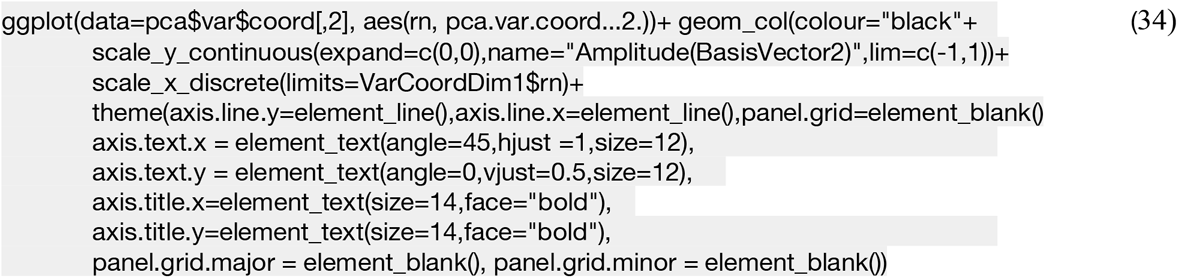

**Figure 33.**
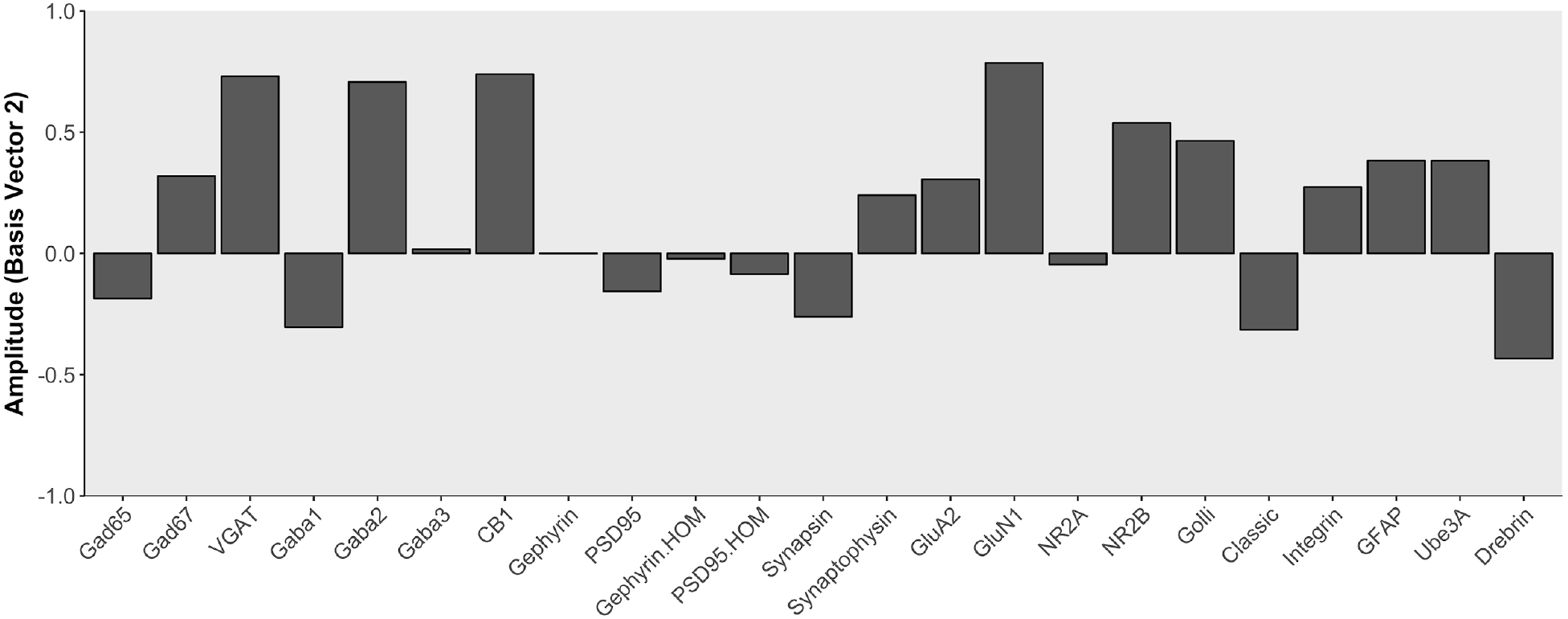
Amplitude of each protein about the second principal component. Protein amplitudes are obtained from the vectors each protein creates in PCA space. The length of these vectors is plotted about the mean of the basis vector for each PCA dimensions, and the direction (positive or negative) is arbitrarily assigned. Here the basis vector being examined is basis vector 2, and proteins (x-axis) with greater variability about this vector have larger amplitudes, while proteins with less variability about basis vector 2 have smaller amplitudes. When bars move in the same direction, it means proteins change together, and when they move in opposite directions it means protein changes are in opposition.

The basis vector plot illustrated the amplitude and direction that each protein contributed to the PC dimension. We identified proteins with the largest amplitude and pointing in the same direction and summed them. Next, we identified proteins with a large amplitude that were pointing in opposite directions and used them to calculate differential indices. Finally, we used a targeted approach based on published information about functional pairs or groups of proteins such as the GluN2A:GluN2B balance that regulates NMDAR kinetics (39), and LTP (40). Those 3 steps identified combinations of proteins that we called candidate plasticity features.

### 4.vii) Validating plasticity features

After identifying candidate plasticity features, they were calculated from the protein expression and correlated with the each of the 3 PC dimensions using the same the same steps described for the coding example (15).

The matrix of correlations between the PC dimensions and candidate plasticity features was inspected paying close attention to features that included proteins differentially weighted by RSKC and PCA. For example, although RSKC reduced the weights of the individual MBP proteins an index calculated using both proteins (Classic-MBP:Golli-MBP) was significantly correlated with PC2 (red). In contrast, RSKC also reduced the weight of GluN2A, but the candidate feature (NR2A:NR2B index) was not correlated with any of the 3 PC dimensions (grey-Figure 34). These two examples demonstrate the importance of validating the candidate features; not all features had significant correlations with a PC dimension.

**Figure 34.**
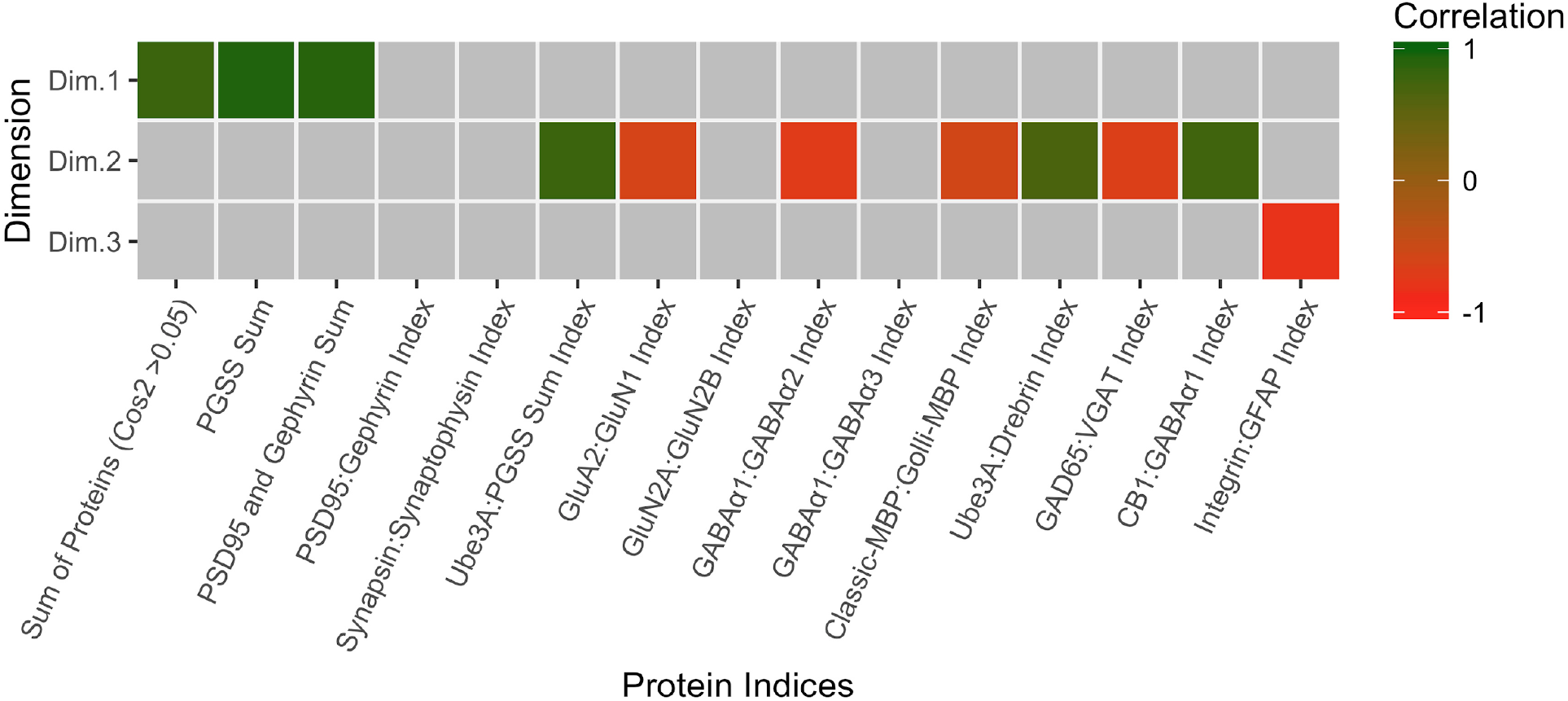
Examination of extracted feature correlation with PCA dimensions. Significant correlations between the first 3 principal components and protein sums / indices are represented as coloured cells. The colour indicates the magnitude (intensity) and direction (positive=green, negative=red) of significant correlations after Bonferroni correction

After exploring the data following the steps above 11 plasticity features were validated for this data set thereby reducing the dimensions from 23 proteins to 11 plasticity features. Furthermore, the steps transformed protein expression into features consistent with combinations of proteins known to regulate plasticity and change during development.

The next step was to use hierarchical clustering with the validated plasticity features to identify the features that had similar patterns of lifespan changes.

### 4.viii) Visualizing feature networks and building plasticity phenotypes

To determine which features had similar patterns of changes across the lifespan we calculated the pairwise correlations between features, ordered those correlations using hierarchical clustering and visualizing the output in a 2D heatmap (Figure 35). The R code used was similar to the heatmap coding example (2)-(6) explained in section 3.ii, except a new object called `NewFeatures2` was made with the 11 plasticity features.

**Figure 35.**
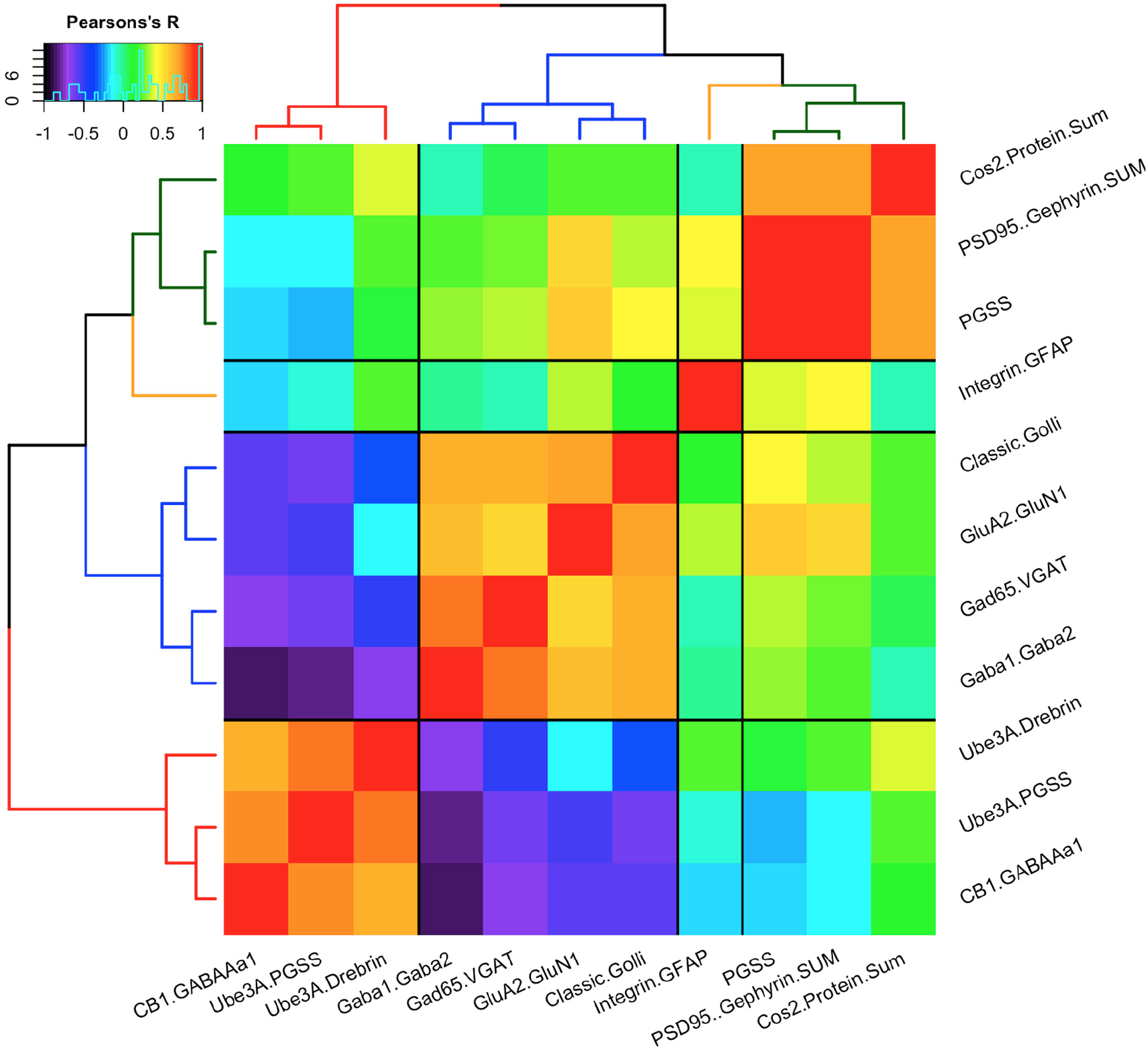
Heatmap of the significant features identified in Figure 34. Matrix of Pearson’s R correlations between the means across 11 plasticity features in each cluster. Clusters were reordered according to the surrounding dendrogram. The dendrogram positioned similar features close together, and moved dissimilar features to the periphery. Inset demonstrates counts across the range of Pearson’s correlations, while the color gradient ranges from low negative correlations (blue/black) to high positive correlations (red).

The correlation heatmap identified 4 groups of plasticity features: first, the 3 protein sums were grouped; second, the astrocyte index Integrin:GFAP was on a separate branch; third, Classic:Golli MBP, GluA2:GluN1, Gad65:VGAT, and GABA_A_α1:GABA_A_α2 were grouped; finally, markers involved with GABAergic inhibition onto pyramidal neurons that included Ube3A:Drebrin, Ube3A:PGSS, CB1:GABA_A_α1 indices were separated from the other groups.

The order of the plasticity features found above by hierarchical clustering was used for building the visualization of the plasticity phenotype (Figure 36). The 11 features that create the plasticity phenotype were visualized using the same method described in section 3.vii with the R coding examples (21) and (22) except that the object NewFeatures2 with the 11 plasticity features was the input for this step. Briefly, the code in (21) determines the color scale for each feature and the code in (22) uses those color-codes to create the visualization of the 11 features and 6 clusters.

**Figure 36.**
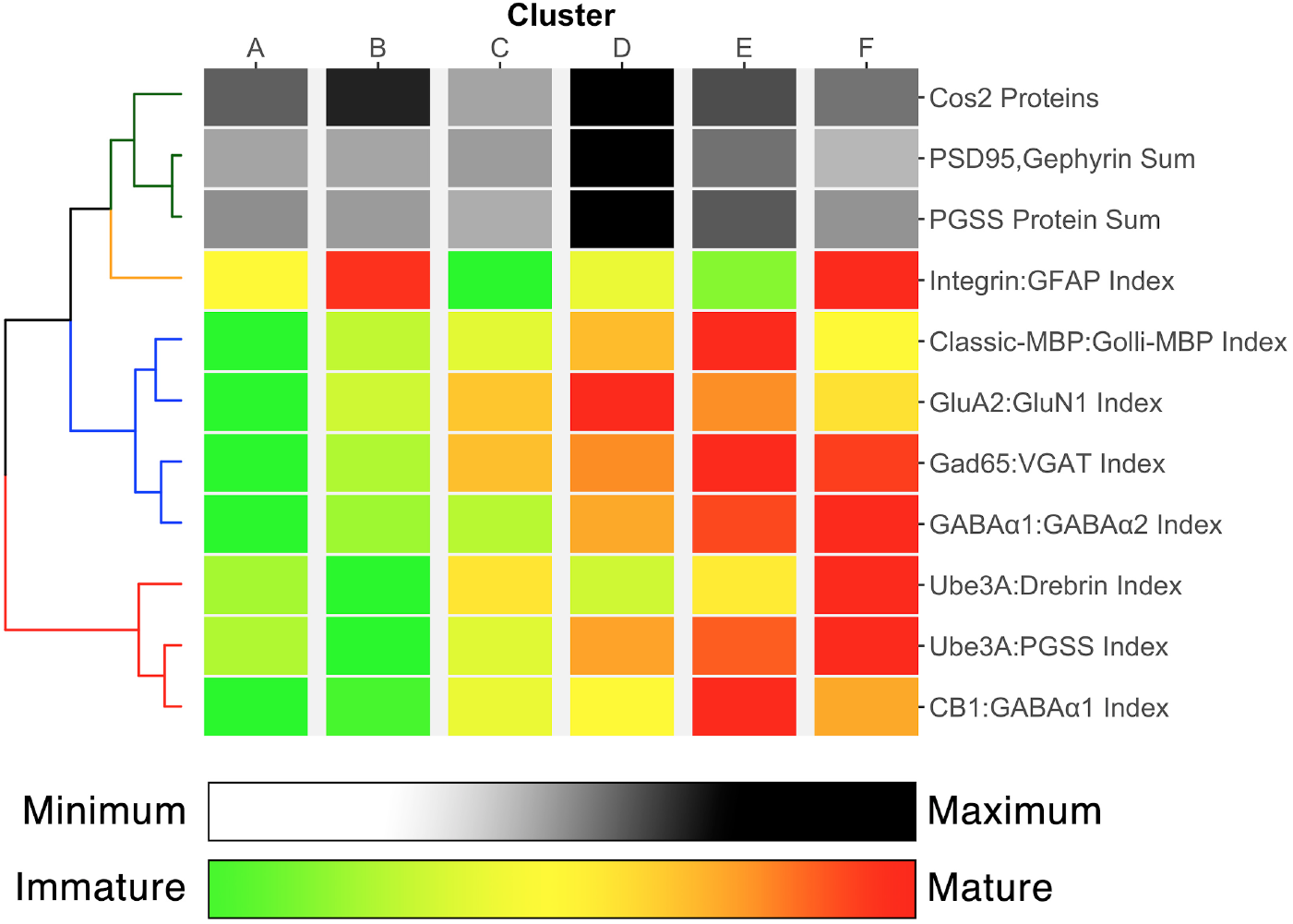
Average plasticity phenotype for each of the 6 clusters representing human visual cortical development. The order of features in the phenotypes was determined from the dendrogram in Figure 35. Most features transition gradually from an immature to mature state. The progression from light grey to dark grey in the top three bars, and from green to red in the last 8 bars indicate the pace of development.

The plasticity phenotype visualization conveys a substantial amount of information about the development of human visual cortex and helped to identify the features that partitioned samples into one cluster versus another (Figure 36). For example, two features that separated clusters A and B were the Integrin:GFAP index and the sum of cos^2^ proteins. Importantly, because some clusters had overlapping age ranges (e.g. B & C Figure 21), comparing features for those clusters can help to identify neurobiological mechanisms that contribute to heterogeneity during different stages of the lifespan.

## 5. Extensions and Discussion

Here we described two workflows and provided example R code for applying a data-driven approach that uses high-dimensional analyses to study development and plasticity of the visual cortex. The workflows were developed to help with interpreting data sets that include multiple proteins (or genes) and ages or rearing conditions where there are complex patterns of results. Also, we extended previous work by developing steps to build, visualize, and compare plasticity phenotypes that capture the neurobiological features that characterize clusters in the visual cortex data sets.

The workflow in section 3 is similar to current approaches applied to study visual cortex that used sequential steps beginning with dimension reduction (e.g. PCA, tSNE) and then cluster analysis (e.g. refs). We used this to study how different forms of visual experience changed protein expression in visual cortex and added the plasticity phenotype to determine the plasticity features that differentiated among the conditions. The plasticity phenotype is a new way of analyzing visual cortex development and plasticity that can facilitate meaningful interpretation of complex patterns of protein (or gene) changes.

The workflow in section 4 separated the cluster analysis and the steps for dimension reduction and feature identification into parallel streams. That change was needed to address the challenges of analyzing the neurobiological development of human visual cortex. Those studies typically use a small number of post-mortem tissue samples (*n*) but measure many proteins or genes (variables, *p*) resulting in a *p* ≈ *n* or *p>n* data structure that is called sparse. Here we compared a group of sparse high-dimensional cluster analysis methods and determined that a recent approach, Robust Sparse K-Means Clustering, RSKC (32), did the best job of partitioning the data into age-related clusters. We also described how to use information from both the RSKC and PCA to identify features for constructing the plasticity phenotype to characterize the neurobiological development of human visual cortex.

Both of our workflows build on established methods for high-dimensional data analysis, and each one describes the reasons for applying different approaches for analyzing the data sets. Since research on cluster analysis is a rapidly moving field, we anticipate that new methods, especially for adaptive clustering of sparse data, may perform better than RSKC. The current workflows are flexible and can be adapted to use new algorithms by exchanging a few lines in the example R code. For example, growth mixture models for cluster analysis are being developed that may be more suitable for handling the types of data structures common for studies of human cortical development (41).

Identifying clusters and exploring cluster content performed well for the example data set but both workflows need to be tested with other data sets. In particular, extension of the feature selection steps (3.iv & 4.v) should include unsupervised methods for identifying features and transforming data. Similarly, RSKC parameters in section 4.iv were selected with an exploratory process and manual inspection of cluster results that could be improved once unsupervised methods are developed.

A valuable contribution from this primer is the development and implementation of plasticity phenotypes. In particular, the plasticity phenotype visualization provides an elegant way to facilitate meaningful interpretation of the results in a single 2D plot. It is not clear, however, how well the current method for selecting features for the phenotypes will scale up for studies using larger numbers of proteins or genes. Perhaps additional methods for data reduction will need to be tested to optimize the selection of neurobiological features in larger data sets.

The R package “v1hdexplorer” aggregates the various packages and custom visualization code used in this paper, and is available for download using the function install_github(“balsorjl”/ “v1hdexplorer”). The custom visualization scripts are included in this document (e.g. coding examples (21) and (22) for visualizing the plasticity phenotypes).

## Supporting information

Supplemental bootstrap analysis

Authors’ contributions
JB Designed research, performed research, analyzed data, wrote/revised the paper; DJ analyzed data, revised the paper; KM designed research, performed research, analyzed data, wrote/revised the paper.

