## Supplemental bootstrap analysis for "A primer on high-dimensional data analysis workflows for studying visual cortex development and plasticity"

**Authors’ contributions:** JB Designed research, performed research, analyzed data, wrote/revised the paper; DJ analyzed data, revised the paper; KM designed research, performed research, analyzed data, wrote/revised the paper.

**Funding Sources:** NSERC Grant RGPIN-2015-06215 awarded to KM, Woodburn Heron OGS awarded to JB.

**Data Availability:** The data used to support the findings of this study are available from the corresponding author upon request.

### Supplemental Analysis

**Bootstrap analysis using custom R code**

To examine the probability that observations in one group (e.g. reverse occlusion-RO) were statistically different from another group (e.g. Normal animals) we performed a one-way bootstrap analysis. First, the *observed* parameters of the experimental group (RO) were used to create a *simulated* data set with 1,000,000 points that had the same mean (mean.RO) and standard deviation (stdev.RO) as the observed subset of RO data. The object sim.RO contains the simulated data set of RO samples.

###### sim.RO <- rnorm(1000000, mean.RO, stdev.RO)

Next, we modelled *experimental* data of the normal group by drawing, at random, from the *simulated* RO data set (sim.RO) the same number of samples as the *observed* subset of Normal animals (NNormal). We calculated the mean of this first *experimental* subset of data, and replaced all data points in sim.RO. This experiment was repeated 100,000 times, and stored in the object resamples.NormalRO.

###### resamples.NormalRO <- lapply(1:100000 ,function(i) sample(sim.RO, NNormal, replace = T))

Each of these 100,000 experiments were reduced to the experimental mean. This subset of data represent a simulated population of comparisons between Normal and RO animals. We calculated the mean of this simulated population, and saved it as mean.resamples.NormalRO.

###### mean.resamples.NormalRO <- sapply(resamples.NormalRO, mean)

To determine the probability that the mean of the *observed* Normal subset of data (mean.Normal) came from the observed RO subset of data (mean.RO), we compared mean.Normal against the population in mean.resamples.NormalRO. To calculate the exact probability (p-value) that the observed Normal group (mean.Normal) came from the mean.resamples.NormalRO population, we calculated the percentage of data points that fall above or below mean.Normal in the distribution of data points in mean.resamples.NormalRO.

###### Norm.v.RO.pval <- 1-(sum(mean.resamples.NormalRO<mean.Normal)/100000)

###### Norm.v.RO.pval<- if (mean.RO<mean.Normal){

###### 1-(sum(mean.resamples.NormalRO<mean.Normal)/100000)

###### } else if (mean.RO>mean.Normal) {

###### 1-(sum(mean.resamples.NormalRO>mean.Normal)/100000)

##### }

In order to compare other conditions against the normal subset of data, adjust the subset of data that is called in place of RO (eg. BD versus Normal). In order to change the direction of the comparison against Normal to be against a different group (eg. MD), exchange the comparator group for Normal in all of the above statements (eg. BD versus MD). The resultant data can be called into a text file and stored for later use when constructing histograms
